# A two-site flexible clamp mechanism for RET-GDNF-GFRα1 assembly reveals both conformational adaptation and strict geometric spacing

**DOI:** 10.1101/2020.09.13.286047

**Authors:** Sarah E. Adams, Andrew G. Purkiss, Phillip P. Knowles, Andrea Nans, David C. Briggs, Annabel Borg, Christopher P. Earl, Kerry M. Goodman, Agata Nawrotek, Aaron J. Borg, Pauline B. McIntosh, Francesca M. Houghton, Svend Kjær, Neil Q. McDonald

## Abstract

RET receptor tyrosine kinase plays vital developmental and neuroprotective roles in metazoans. <u>G</u>DNF family ligands (GFLs) when bound to cognate GFRα co-receptors recognise and activate RET stimulating its cytoplasmic kinase function. The principles for RET ligand-co-receptor recognition are incompletely understood. Here we report a crystal structure of the cadherin-like module (CLD1-4) from zebrafish RET revealing interdomain flexibility between CLD2-CLD3. Comparison with a cryo-EM structure of a ligand-engaged zebrafish RET^ECD^-GDNF-GFRα1 complex indicates conformational changes within a clade-specific CLD3 loop adjacent to co-receptor. Our observations indicate RET is a molecular clamp with a flexible calcium-dependent arm that adapts to different GFRα co-receptors, while its rigid arm recognises a GFL dimer to align both membrane-proximal cysteine-rich domains. We also visualise linear arrays of RET^ECD^-GDNF-GFRα1 suggesting a conserved contact stabilises higher-order species. Our study reveals ligand-co-receptor recognition by RET involves both receptor plasticity and strict spacing of receptor dimers by GFL ligands.

**Highlights:**

- Crystal structure of zebrafish RET cadherin-like module reveals conformational flexibility at the calcium-dependent CLD2-CLD3 interface
- Comparison of X-ray and cryo-EM structures indicate conformational differences between unliganded and liganded RET involving a clade-specific CLD3 loop
- Strict spatial separation of RET^ECD^ C-termini is imposed by each cysteine-rich domain interaction with GFL dimer
- Differences in co-receptor engagement and higher-order ligand-bound RET complexes indicate potentially divergent signalling mechanisms

## Introduction

Neurotrophic factors fulfil an essential function to support and protect both developing and mature neurons (Henderson et al., 1994). In view of their neuroprotective therapeutic potential there has been much interest in understanding how they engage and activate their cell surface receptors (Airaksinen and Saarma, 2002; Allen et al., 2013). The glial cell-line derived neurotrophic factor (GDNF) family ligands (GFLs) constitutes an important family of neurotrophic factors that include GDNF (Durbec et al., 1996), Neurturin (NRTN) (Kotzbauer et al., 1996), Artemin (ARTN) (Baloh et al., 1998b), Persephin (PSPN) (Airaksinen and Saarma, 2002; Milbrandt et al., 1998) and more recently GDF15 (Emmerson et al., 2017; Hsu et al., 2017; Mullican et al., 2017; Yang et al., 2017). Each of these soluble factors are covalent dimeric ligands and are members of the cystine-knot/TGF-ß superfamily (Hinck et al., 2016). Each GFL has a preferred, high affinity cognate GFR-alpha (GFR) co-receptor that associate as GDNF-GFRα1 (Cacalano et al., 1998), NRTN-GFRα2 (Baloh et al., 1997), ARTN-GFRα3 (Baloh et al., 1998a), PSPN-GFRα4 (Thompson et al., 1998) and GDF15-GFRAL (Emmerson et al., 2017; Hsu et al., 2017; Mullican et al., 2017; Yang et al., 2017) complexes respectively. The GFL co-receptors each consist of three related helical domains (D1 to D3) and are anchored at the membrane either through glycosylphosphatidylinositol (GPI) linkages (GFRα1-4) or by a transmembrane helix (GFRAL). The bipartite GFL-GFR complexes are recognised by the RET receptor tyrosine kinase (RTK) forming ternary RET-GFL-GFR complexes (Cacalano et al., 1998; Durbec et al., 1996; Jing et al., 1996; Treanor et al., 1996). Engagement of GFL-GFR by RET triggers RET auto-phosphorylation of critical tyrosine and serine residues to activate intracellular signalling pathways, (Ibáñez, 2013; Mulligan, 2014).

RET is unique among RTKs as it has four consecutive cadherin-like domains [CLD(1-4)] and a membrane-proximal cysteine-rich domain (CRD) in its extracellular domain (RET^ECD^) (Anders et al., 2001). The CLD domains diverge significantly, in sequence, structure and arrangement from classical Cadherins (calcium-dependent adhesion) (Anders et al., 2001; Brasch et al., 2012; Kjær et al., 2010). Cadherin-like domains CLD(1-2) form a closed clamshell arrangement with structural differences evident between higher and lower vertebrate RET orthologues (Kjær et al., 2010). Calcium ions are critical for RET folding consistent with the conservation of classical Cadherin calcium-coordinating motifs between CLD2 and CLD3 (Anders et al., 2001; Kjær and Ibáñez, 2003; van Weering et al., 1998). Biochemical efforts to map the bipartite GDNF-GFRα1 binding site within RET^ECD^ to a minimal-binding domain have implicated the entire RET^ECD^ region. This contrasts many receptor-ligand interactions RTKs that frequently map to a ∼200 aa minimal-binding domain (Lemmon and Schlessinger, 2010). Two key interactions between RET^ECD^–GFRα1 and RET^ECD^–GDNF were identified from electron microscopy structures of RET^ECD^ ligand recognition of GDNF/NRTN and GFRα1/GFRα2, though lacking a CRD structure (Bigalke et al., 2019; Goodman et al., 2014). A recent study by Li and co-workers used cryo-EM to reveal the full human RET^ECD^, including the CRD, in complex with several GFL ligands. In these structures, the D1 domain of GDNF-GFRα1 or GDF15-GFRAL complexes with RET^ECD^ were missing (Li et al., 2019). Moreover, little information about conformational changes upon ligand binding was apparent from these studies. Understanding the dynamics and conformational alterations induced on ligand-binding is important for the design of RET modulators with potential therapeutic applications.

Here we report both crystallographic and cryo-EM structures of zebrafish RET^CLD-4^ and RET^ECD^-GDNF-GFRα1a complex respectively. We observe plasticity within the zRET^CLD1-4^ and define the extent of conformational changes induced by ligand-co-receptor binding. Conformational adaptions are observed between RET and GFRα contacts even across clades, whereas a more strictly conserved interaction is observed between GFL and RET-CRD close to the transmembrane region. We also describe zRET^ECD^-GDNF-GFRα1a multimers on cryo-EM grids generating both linear and 2D arrays. Insights from this study support a dual-site clamp mechanism involving an adaptive interaction site for co-receptor recognition and an alignment interaction site between a GFL dimer and RET CRD for signalling.

## Results

### Crystal structure of zebrafish RET CLD(1-4) indicates localised flexibility

Crystals were obtained for a zebrafish RET construct spanning residues 22-504 (zRET^22-504^) with glycosylation site mutations, N259Q, N308Q, N390Q and N433Q (defined hereafter as zCLD(1-4)^red.sug.^). Diffraction data from these crystals led to a structure determination at 2.2Å resolution (Figure 1; Supplementary Table 1 and Methods). The final zCLD(1-4)^red.sug.^ model contains residues 22 to 498 and includes seven N-linked glycans well resolved in the electron density (Supplementary Figure 1). The crystals adopted the triclinic space group P1 and contained two molecules of CLD(1-4)^red.sug.^ within the asymmetric unit. Each had a similar overall structure but with different hinge angles between CLD2 and CLD3, pointing to flexibility within RET (Supplementary Figure 2).

**Figure 1:**
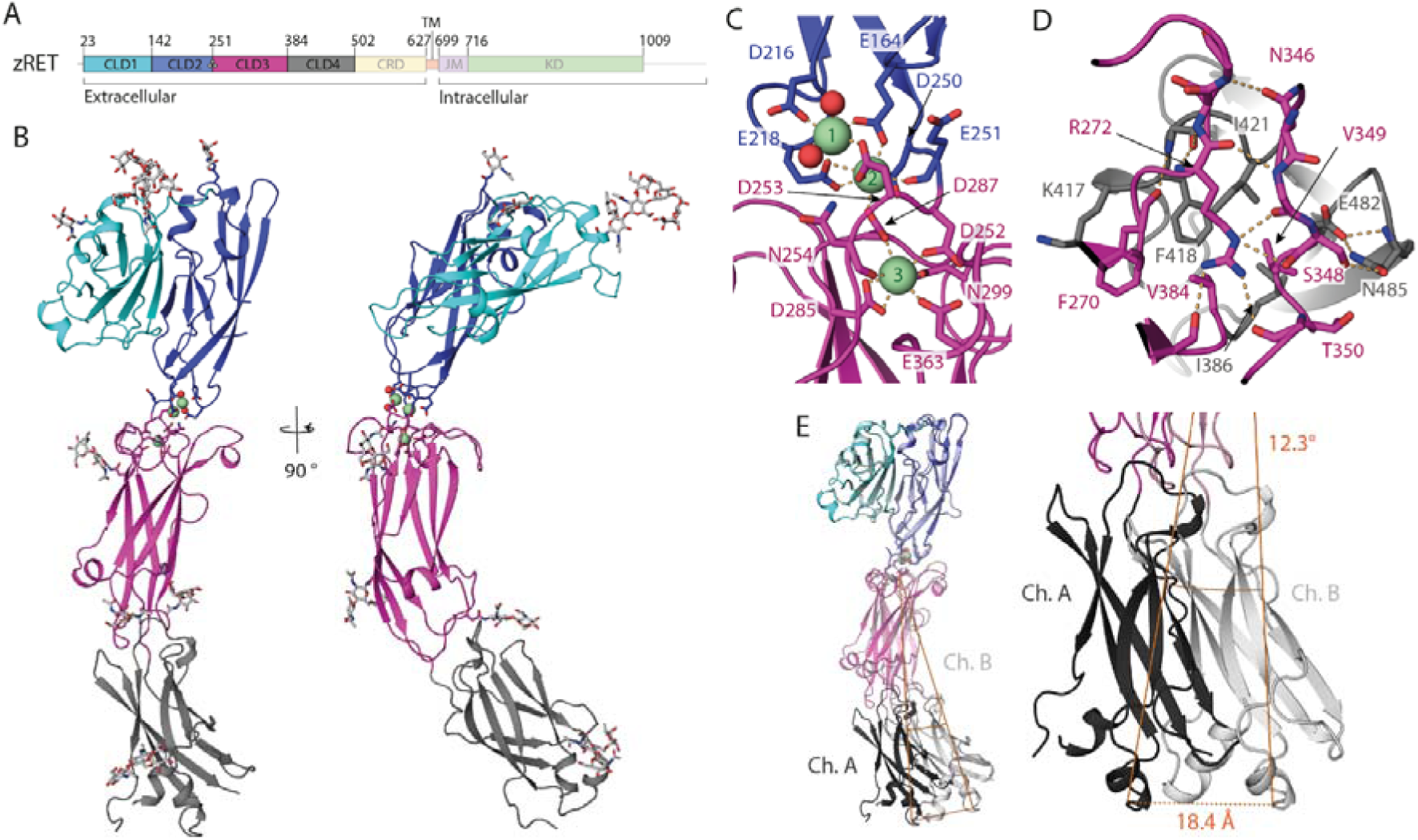
Crystal structure and flexibility of the zRET CLD(1-4) module. A) Schematic of zebrafish RET receptor tyrosine kinase; CLD cadherin like domains, CRD cysteine rich domain, TM transmembrane helix, JM juxtamembrane domain and KD kinase domain. B) Orthogonal views of zRET^CLD1-4^, with CLD1 in cyan, CLD2 in dark blue, CLD3 in magenta and CLD4 in grey. The calcium-binding site between CLD(2-3) has 3 calcium ions as green spheres with coordinating ligands shown as sticks and waters represented as red spheres. C) Closeup view of the coordination shell for the three calcium atoms between CLD2 and CLD3. D) Close-up of the interface between CLD3-CLD4 centred on R272, selected sidechains shows as sticks and dashed lines for hydrogen bonds. E) Superposition of chains A and B within the asymmetric unit, aligned through their CLD(1-2) domains, shows a shift of 18.4Å and a rotation of 12.3 ° pivoting at a hinge close to the calcium-sites. All structural images were generated in PyMOL (Schrodinger, 2015).

**Figure 2:**
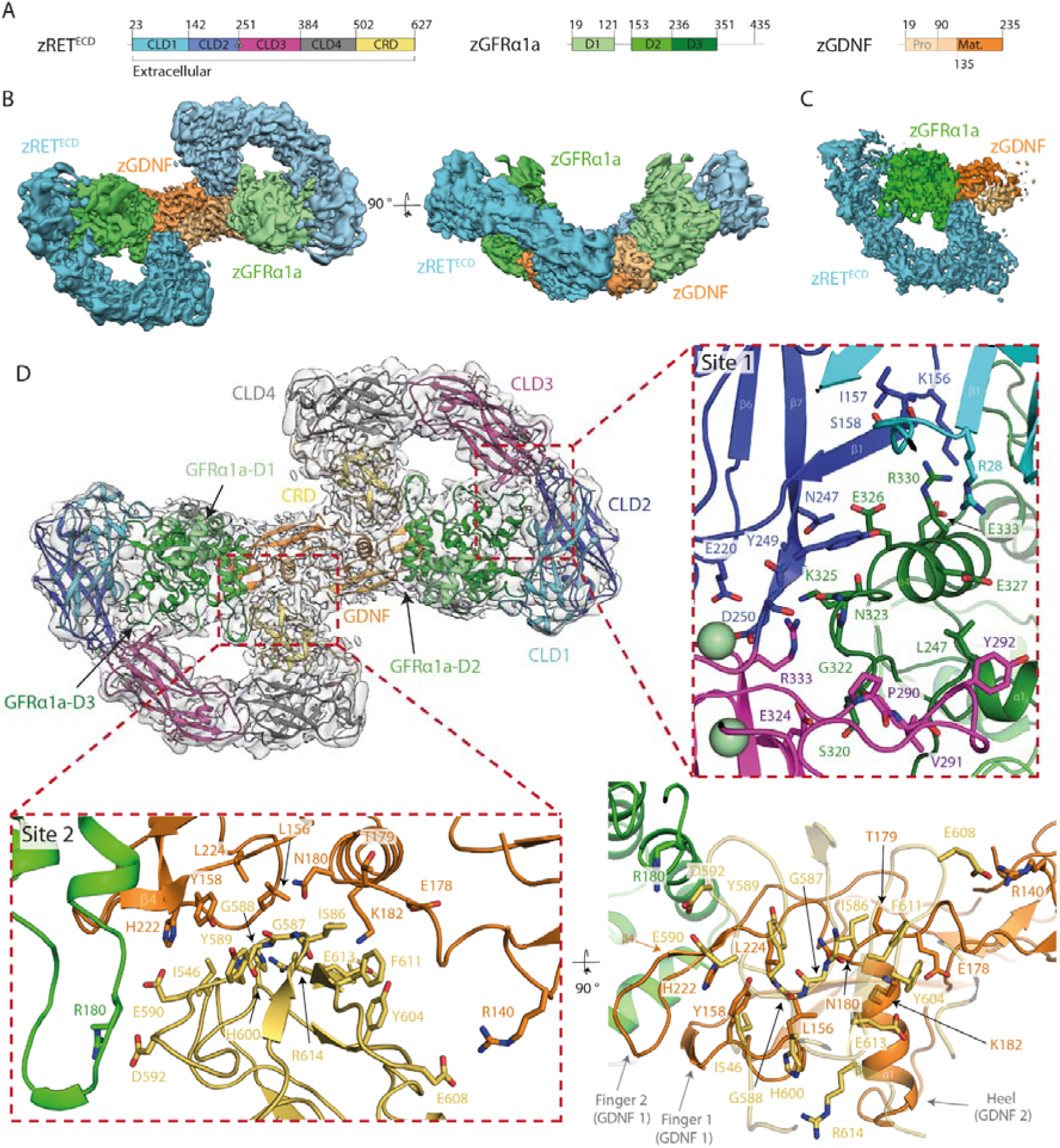
Cryo-EM structure of the zRET^ECD^-zGFRα1a^D1-3^-zGDNF^mat.^ (zRGα1a) complex. A) Schematic of zRET^ECD^, zGFRα1a^D1-3^ and zGDNF^mat.^, colour coded according to Figure 1. B) Orthogonal views of the reconstituted zRGα1a complex cryo-EM map, projecting down the approximate molecular dyad or perpendicular to it. The cryo-EM map is segmented and coloured by protein, with zRET^ECD^ cyan, zGFRα1a^D1-3^ green and zGDNF^mat.^ orange. C) Symmetry-expanded map of zRGα1a half-complex, with the map segmented and coloured by protein as per panel B). D) The final model of the zRGα1a complex, shown as a cartoon, built into the C2 symmetry map, coloured light grey, (map contoured at 0.24). The domains are coloured as per Figure 1 with zRET^ECD^ CLD1 is cyan, CLD2 is dark blue, CLD3 is magenta, CLD4 is grey, and CRD is yellow, for GFRα1a domains D1-3 are pale green, green and dark green respectively, the two molecules of zGDNF^mat.^ are orange and pale orange. Two sites of interaction between zRET^ECD^ and zGDNF^mat.^-zGFRα1a^D1-3^ complex are highlighted by red dashed boxes, labelled as site 1 (zGFRα1a-zRET) and site 2 (zGDNF-zRET). Interaction residues are highlighted as sticks and the backbone represented as cartoon. Images of the map were produced in Chimera (Pettersen et al., 2004). Image of the cartoon model in panel D was produced in PyMOL (Schrodinger, 2015).

The overall structure of zCLD(1-4)^red.sug^ showed that all CLDs have the predicted canonical seven β-strand sandwich architecture of cadherin domains (Supplementary Figure 3) (Shapiro and Weis, 2009). The amino-terminal CLD1 is packed against CLD2 in a fold-over clamshell arrangement as anticipated from human RET, while CLD(2-4) forms a “C-shape” (Figure 1B). The zCLD(1-2) clamshell has a surprisingly large overall root mean square deviation (rmsd) of 18.9 Å over 229 C-alphas when superposed with hCLD(1-2)(Winn et al., 2011). Key features contributing to this structural divergence are the different disulfide connectivity, a lack of a ß-hairpin and a longer CLD1 helix α1 between higher and lower vertebrates (Supplementary Figure 4) (Kjær et al., 2010).

**Figure 3:**
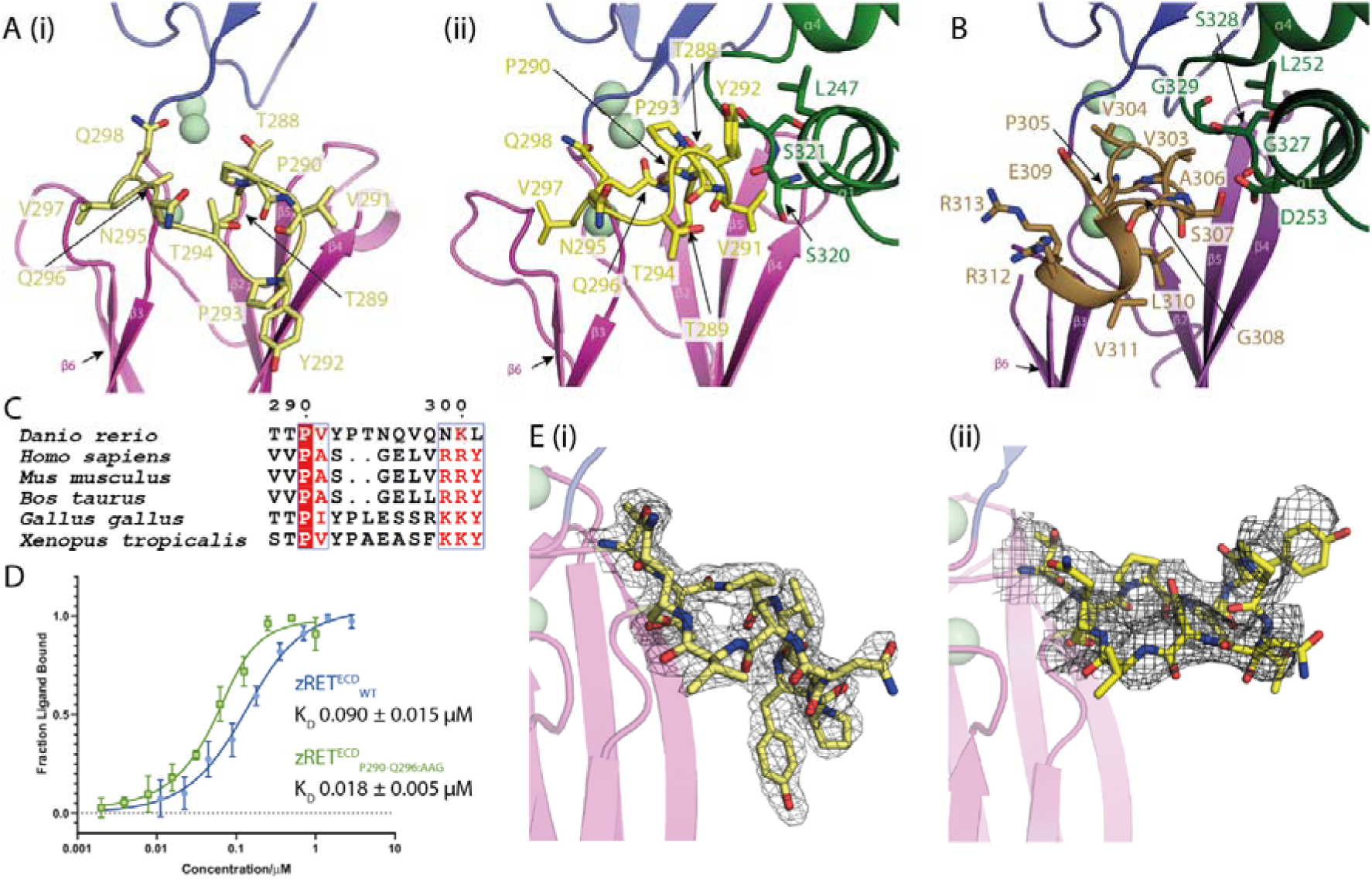
Ligand-co-receptor induced conformational changes in zRET^ECD^: A) The CLD3-β2-β3-loop shown in yellow as sticks (i) projects “downwards” in the view shown for zRET CLD(1-4) (see the orientation of Y292 sidechain) and (ii) projects “upwards” to engage GFRα1^D2^ α1 helix (green sticks) in the zRGα1a structure. B) The CLD3-β2-β3-loop from the human RET^ECD^-NRTN-GFRα2 structure (PDB 6Q2O) shown as olive coloured sticks, domains coloured as per Figure 1. C) Sequence alignment of RET CLD3-β2-β3-loop including *Homo sapiens* (Uniprot P07949), *Mus musculus* (Uniprot P35546), *Bos Taurus* (Uniprot F1MS00), *Gallus gallus* (Uniprot F1NL49), *Xenopus tropicalus* (Uniprot F7DU26) and *Danio rerio* (Uniprot O42362). Sequences are coloured by similarity using Espript (http://espript.ibcp.fr) (Robert and Gouet, 2014). D) Binding curves and calculated K_D_’s for zRET^ECD^_WT_ and mutant (zRET^ECD^_P291-Q296;AAG_) binding to zGFRα1a_2_ -zGDNF_2_ measured by microscale thermophoresis. E) (i) Electron density map calculated using m2Fo-DFc coefficients over the CLD3-β2-β3-loop, yellow sticks and contoured at 1.0 σ. (ii) Coulombic potential cryo-EM map for CLD3-β2-β3-loop from the zRGα1a complex (black mesh). Calcium ions are represented as pale green spheres. Images were rendered in PyMOL (Schrodinger, 2015).

**Figure 4:**
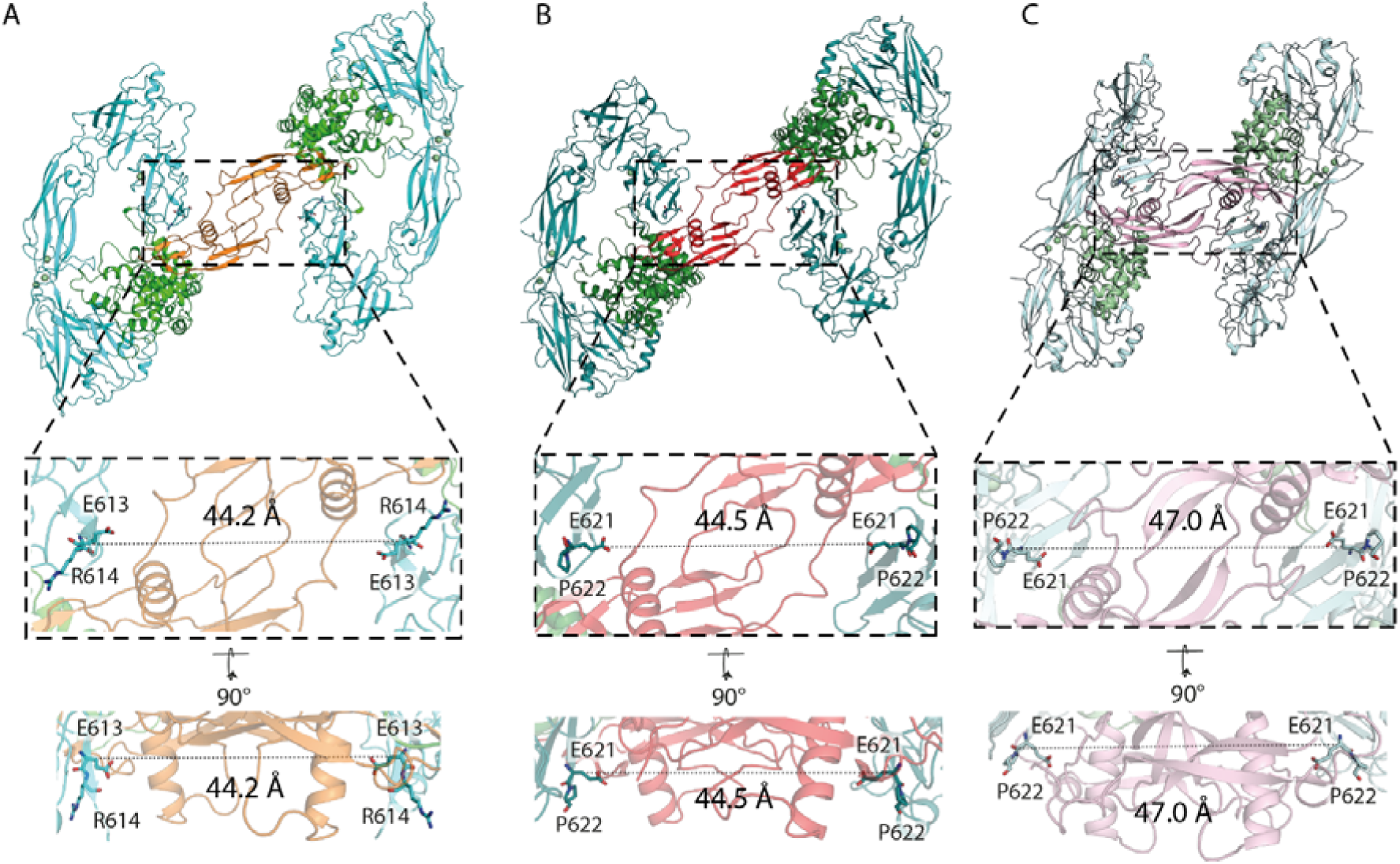
Different GFL ligands establish a conserved spacing between RET CRD-CRD pairs in the zRGα1a ternary complex. A) Separation between the Cα of E613 (equivalent to E620 of hRET) from both molecules of zRET^ECD^ within the zRGα1a structure. B) Equivalent distance between the Cα E620 from both molecules of hRET^ECD^ from the hRET^ECD^-NRTN-GFRα2 (PDB 6Q2O) structure. C) Equivalent separation between the Cα E620 from the 2 molecules of hRET^ECD^ from the hRET^ECD^-GDF15-GFRAL (PDB 6Q2J) structure. Overall structure is represented as a cartoon and the Ca^2+^ ions are represented as spheres. RET is coloured cyan, teal and pale cyan in zRGα1a, hRET^ECD^-NRTN-GFRα2 and hRET^ECD^-GDF15-GFRAL structures respectively. GFRα1a, GFRα2 and GFRAL are coloured green, dark green and pale green, respectively. GDNF, NRTN and GDF15 are coloured orange, red and light pink, respectively. All images were rendered in PyMOL (Schrodinger, 2015).

The irregular CLD2-β1 (residues 153-160) is largely separated from the main CLD2 sheet and lies between CLD1-β1 and CLD2-β7, anchored largely through CLD2-β2 sidechains (such as R172 and R176) rather than mainchain interactions (Supplementary Figure 5). One end of CLD2-β1 is tethered through packing of two short α-helices from CLD2-β1 and CLD2-β2, while the other end is locked down by aromatic sidechains from residues amino-terminal to CLD1-β1. This configuration contributes to a substantial internal cavity between CLD1 and CLD2, with a surface volume of ∼510 Å^3^. We note that analysis of the published human CLD(1-2)(Kjær et al., 2010) (PDB 2×2U) also revealed a similar but smaller internal cavity of ∼324 Å^3^ (Supplementary Figure 5)(Abagyan et al., 1994; An et al., 2005; Fernandez-Recio et al., 2005). On the opposing side of the clamshell interface CLD1-β2 and CLD2-β2 contribute through both side and main chain interactions.

**Figure 5:**
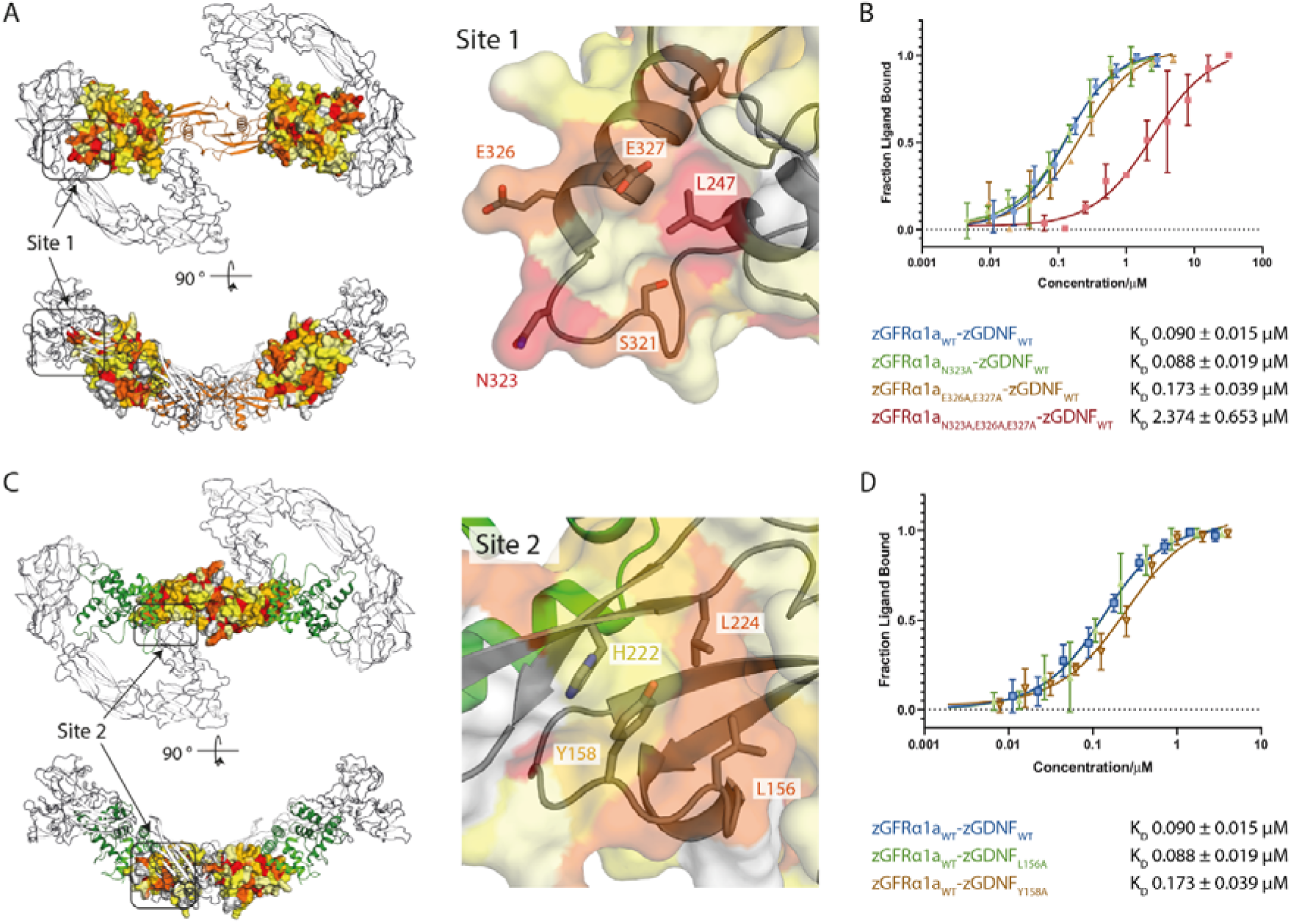
Mutational analysis of zGDNF and zGFRα1 site 1 and 2 interactions with zRET^ECD^. A) Heat map of the sequence conservation between hGFR paralogues, and zGFRα1a [zGFRα1a (Uniprot Q98TT9), hGFRα1 (Uniprot P56159), hGFRα2 (Uniprot O00451), hGFRα3 (Uniprot O60609), hGFRα4 (Uniprot Q9GZZ7), and hGFRAL (Uniprot Q6UXV0)] mapped onto the structure of zGFRα1a D2-D3 domains reported here. Residues are coloured by similarity (red highly similar to yellow through to white, least similar) as described in the methods section. Two orthogonal views are shown and are rendered by PyMOL (Schrodinger, 2015). Right panel, closeup of site 1 and conserved zGFRα1a residues B) Binding curves and K_D_ values obtained using microscale thermophoresis for zGFRα1a^D1-3^ and mutations assessed in complex with zGDNF^mat.^. C) Heat map of the sequence similarity between GDNF paralogues depicted as a surface representation, mapped onto zGDNF^138-235^. Right panel, closeup of site 2 contact between RET^CRD^ and zGDNF dimer. D) Microscale thermophoresis binding curves and K_D_ values for zGDNF and mutations L156A and Y158A probed in complex with wild type zGFRα1a binding to zRET^ECD^.

The limited size of the CLD(2-3) interface is typical of a calcium-dependent cadherin domain pair, with three calcium ions (Ca-1/Ca-2/Ca-3) proximal to the two domains (Figure 1C)(Shapiro and Weis, 2009). Ca-1 and Ca-2 lie in close proximity to one another (3.9 Å apart in chain A) and share three coordinating ligands, the side chains of E164, E218 (CLD2) and D253 (CLD3). Ca-1 is exposed to the solvent at the edge of CLD2, with the coordination sphere completed with D216 and two water molecules, one of which is coordinated by with N165 (Figure 1C). The Ca-2 coordination sphere includes D253, a mainchain carbonyl from E251 (CLD2), and D287 (CLD3), which is a ligand shared with Ca-3 (Figure 1C). Ca-3 is buried within CLD3 and located 6.9 Å away from Ca-2, the coordination shell is completed with the side chains of D252, D285, N299 and D363 and the mainchain carbonyl of N254 (Figure 1C).

CLD3 consists of 135 amino acids and is the largest RET CLD. It shows the greatest structural divergence of all CLDs (∼5Å rmsd) compared to the smaller canonical cadherin domains (Supplementary Figure 3) (Shapiro and Weis, 2009). Additional elements within CLD3 include a loop insertion between β2-β3 adjacent to the calcium-binding site, an α-helix between β3 and β4, and a much longer pair of antiparallel β-strands β4 and β5. Unusually, CLD3 lacks any disulfide bonds and its CLD4 interface is offset at one side of the domain giving a pronounced curvature to the entire CLD(1-4) module. CLD3 has five potential glycosylation sites (NetNGlyc prediction server, http://www.cbs.dtu.dk/services/NetNGlyc/); two were removed by site-directed mutagenesis in zCLD(1-4)^red.sug^ and three are visible in the electron density (Supplementary Figure 1). These features collectively ensure CLD3 plays a crucial role in the stability and curvature of the zCLD(1-4) module.

The CLD(3-4) interface diverges substantially from classical cadherins and has previously confounded efforts to predict the precise CLD(3-4) domain boundaries (Anders et al., 2001). It lacks calcium ions and has a predominantly hydrophobic character, with peripheral hydrophilic interface contacts (Figure 1D). Hydrophobic contacts include CLD3 sidechains F270 and V349 that pack against CLD4 F418 and I421 sidechains and are tethered by V384 from a rigid connecting linker with sequence P383-V384-P385. An exception to the hydrophobic character of the interface is the buried R272 sidechain from the CLD3-β1-β2-loop (Figure 1D). The aliphatic portion of R272 packs against V349, V384 and I421, while its guanidinium head engages mainchain carbonyls on the CLD3-β5-β6-loop and the CLD3-CLD4 linker (Figure 1D). This residue is equivalent to R287 in humans, a known site of mutation in a severe form of Hirschsprung’s disease (R287Q), highlighting the crucial nature of this residue for folding (Attie et al., 1995; Pelet et al., 1998).

Differences in the CLD interface size indicate flexibility between CLD2 and CLD3 but rigidity between CLD3 and CLD4. This is supported by superpositions of the two independent molecules of zCLD(1-4)^red.sug^ demonstrating plasticity in the tapered CLD(2-3) interface (Figure 1E, Supplementary Figure 2). Superimposing chain B onto chain A, aligning through CLD(1-2) revealed the rigid CLD(3-4) module pivots about the CLD(2-3) calcium binding site interface with a variation of 12.3° which leads to a difference of 18.4Å at the furthest point from the CLD(2-3) interface (Figure 1E). Subtle angular differences proximal to the calcium ions, propagating down the module lead to a tightening of the C-shaped structure between chain A and chain B (Supplementary Figure 2).

### Cryo-EM structure of the ternary zebrafish GDNF-GFRα1a-RET^ECD^ complex

A reconstituted complex was assembled consisting of the zRET^ECD^ (residues 1-627), a C-terminal truncated zGFRα1a (zGFRα1a^D1-3^) covering residues 1-353, and an N-terminal truncated zGDNF, residues 135-235, (zGDNF^mat.^), defined hereafter as zRGα1a from <u>R</u>ET-<u>G</u>DNF-GFR <u>α1a</u> (Figure 2A, Supplementary Figure 6). The zRGα1a complex homogeneity and stability was substantially improved by crosslinking the sample with glutaraldehyde using the GraFix technique (Kastner et al., 2008). An initial cryo-EM dataset (dataset 1) collected on the reconstituted zRGα1a yielded a 3D cryo-EM map that confirmed a 2:2:2 stoichiometry, consistent with SEC-MALLS data (Supplementary Figure 6) and similar to recently published human RET complexes (Bigalke et al., 2019; Li et al., 2019). The map displayed substantial anisotropic resolution due to particle orientation bias on the gridS. To overcome this, a second dataset was collected with a sample grid tilted at an angle of 30° (dataset 2) (see Supplementary Figure 7). The combined particles from both datasets were used to generate an initial 3D volume with C2 symmetry applied in CryoSPARC-2. Additional processing with symmetry expansion in RELION-3 (Kimanius et al., 2016; Scheres, 2012; Zivanov et al., 2018), improved the anisotropy and resolution of the map by addressing flexibility at the 2-fold symmetry axis, to produce a map with a nominal resolution of 3.5Å (Figure 2C, Supplementary Figure 8). Subsequent analysis of this final map with 3DFSC indicated that there were a limited number of particles contributing to the Z-direction of the 3D reconstruction which resulted in the resolution in that direction being limited to ∼10Å (Supplementary Figure 8)(Tan et al., 2017).

**Figure 6:**
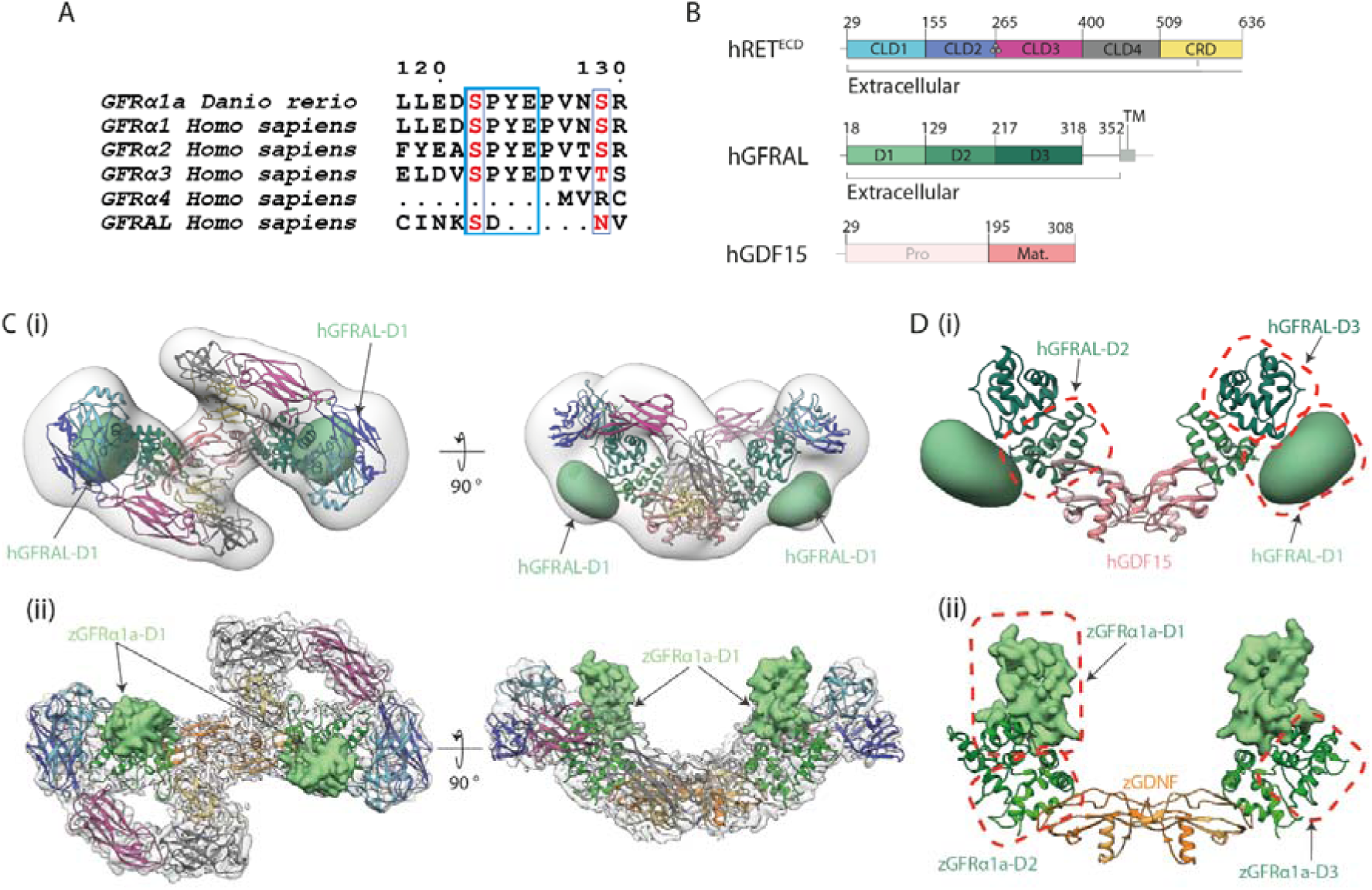
Divergent GFRα1/GFRAL co-receptor D1 domain positions within the RET^ECD^ ternary complex. A) The D1-D2 domain linker motif (SPYE), highlighted in cyan is conserved between zGFRα1a, GFRα1, GFRα2, GFRα3. It is missing from the shorter GFRα4 and the divergent GFRAL. B) Schematic diagram of human RET^ECD^, GFRAL and GDF15 construct boundaries used and individual domains annotated as per Figure 1. C) (i) Negative stain EM envelope of a reconstituted hRET^ECD^_2-_hGDF15_2_-hGFRAL_2_ (hR15AL) complex docked with hR15AL (PDB:6Q2J) revealing additional map potential indicated by a green Gaussian volume (generated from a D1 domain homology model). (ii) Cryo-EM map of zRGα1a (light grey) superposed with the final model (coloured as per Figure 2) with GFRα1a^D1^ shown (light green Gaussian volume at 5 Å^2^). D) Comparison of co-receptor D1 domain position and interfaces (i) GFRAL^D1^ makes different contacts to domains D2-D3 (green), GFRAL^D1^ shown as a 30Å^2^ Gaussian volume (light green), GDF15 salmon. (ii) zGFRα1a^D1^ contacts and coloured as per Figure 2. zGFRα1a^D1^ represented as a 5 Å^2^ Gaussian volume (light green), Images rendered in Chimera (Pettersen et al., 2004)

**Figure 7:**
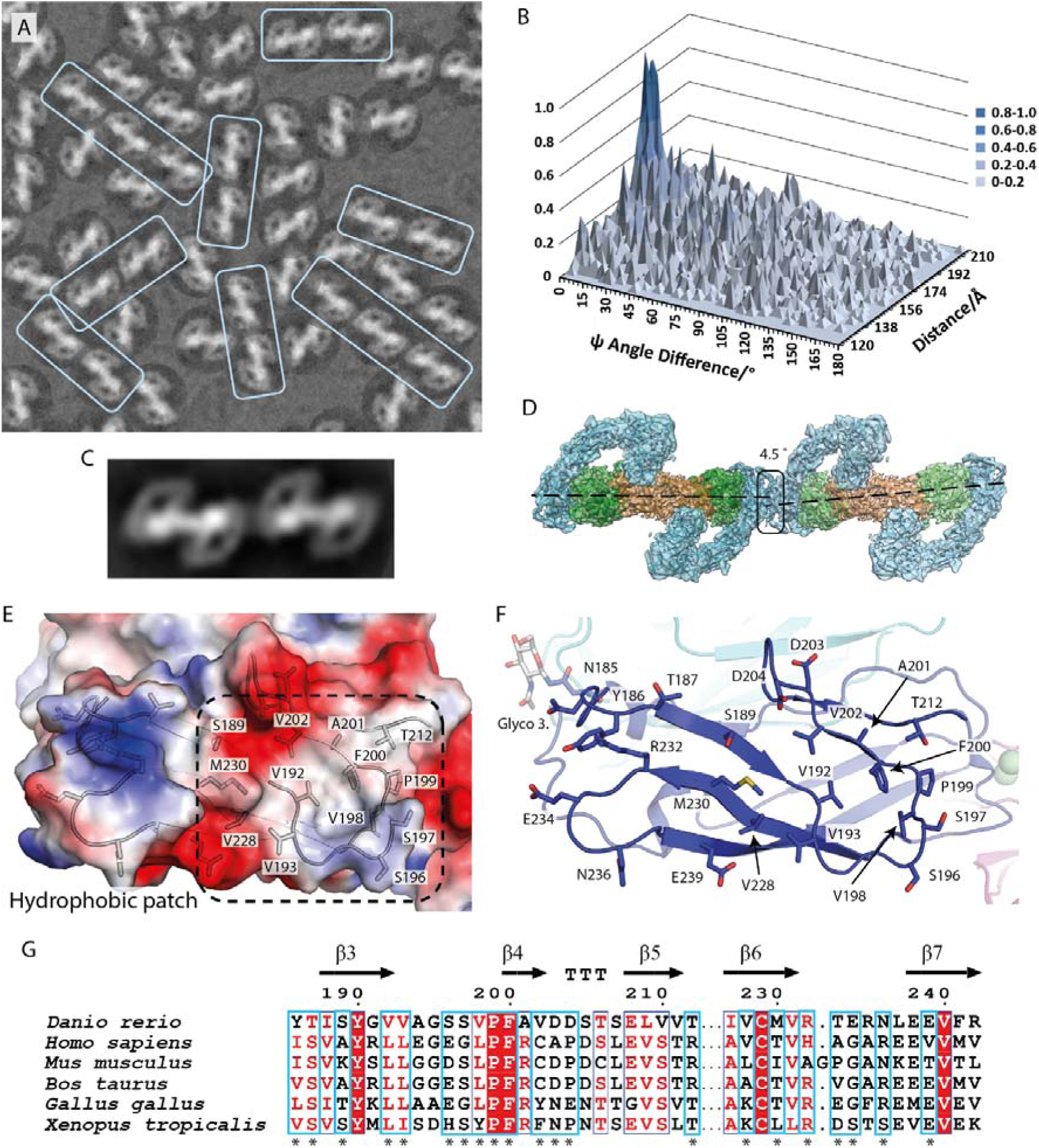
Evidence for linear arrays of zRGα1a particles on cryo-EM grids. A) Close-up of a representative micrograph for un-crosslinked zRGα1a, in which a dominant 2D class average projecting down the molecular dyad is fitted into picked particles from the micrograph using RELION-2.1 (Zivanov et al., 2018). The particle orientation bias is evident from the linear particle arrays highlighted within boxes. B) Statistical distribution of the difference between the angle psi (Δψ) between two particles and their separation distance. Here the angle ψ is defined for each particle as the angle of rotation of each particle required to align it onto the 2D class average. C) 2D class average from automated particle picking contains two adjacent zRGα1a complexes consistent with panel A). D) zRGα1a-zRGα1a interface highlighted with a black box and rendered by Chimera.(Pettersen et al., 2004). The angle and separation between each complex is based on the peak coordinates from panel B) with a Δψ angle of 4.5 ° and separation of 181 Å (with a frequency cut off at 0.5). Both particles are assumed to be at the same Z height. E) An electrostatic potential surface representation of one half of the predicted homotypic zRET^CLD2^ interface, generated in PyMOL (Schrodinger, 2015), indicating a hydrophobic patch central to the interface. A cartoon and stick representation of surface residues are shown (transparent trace). F) Close-up of a cartoon representation for one half of the CLD2-CLD2 interface, highlighting residues at the interface as sticks, coloured as per Figure 1. G) Sequence alignment of representative RET sequences from three higher and three lower vertebrates as per Figure 3. The residues highlighted with an asterix and surrounded with cyan boxes map to the predicted CLD2-CLD2 interface.

The zRGα1a cryo-EM map resembles a figure-of-eight with a molecular two-fold centred at the crossover point (Figure 2B). To enable building of a full structure into the map, we determined a crystal structure of zGDNF^mat^-zGFRα1a^151-353^ lacking domain D1 (referred to hereafter as zGFRα1a ^ΔD1^) at 2.2Å (see Materials and Methods and Supplementary Figure 9). We then fitted crystal structures for zRET CLD(1-4) and zGDNF^mat^-zGFRα1a^ΔD1^ into the symmetry-expanded map (Figure 2C) together with homology models for the zRET^CRD^ and zGFRα1a^D1^. An initial model for zRET^CRD^ was generated from the hRET^ECD^-hGFRα2-NRTN structure (Li et al., 2019) and for zGFRα1a^D1^ from the hGFRα2-NRTN (Sandmark et al., 2018) structure by substituting zebrafish sequences followed by model optimisation using Swiss-Model (Schwede et al., 2003) and Modeller (Webb and Sali, 2016), respectively. The initial structure was refined against the symmetry-expanded map and rebuilt, before placing it into the C2 averaged map for further refinement in PHENIX (Adams et al., 2010) (Supplementary Table 2, Supplementary Figure 10). The final near complete structural model has a cross correlation of 0.63 against this map. The highest resolution features of the map are at the centre of the complex, close to GDNF and zRET^CLD4-CRD^ (Supplementary Figure 8). For example, the intermolecular disulfide bridge which covalently links two GDNF protomers is clearly visible within the volume (Figure 2B). *N*-acetylglucosamine (GlcNAcβ1-Asn) glycan rings linked to asparagine sites were also evident in the map. Density was also evident for zGFRα1a^D1^, sandwiched between zGFRα1a^D3^ and zRET^CLD1^, at a similar position to GFRα2^D1^ (Bigalke et al., 2019; Li et al., 2019; Sandmark et al., 2018) (Supplementary Figure 11).

The final structure shows zGDNF at the core of the complex flanked by two zGFRα1a^D1-3^ co-receptors, both of which are further enveloped by two “G”-shaped RET^ECD^ molecules (Figure 2D). The spur of the RET^ECD^ “G”-shape is formed by the CRD domain making contacts with both GDNF and zGFRα1a, as first predicted from lower-resolution negative stain EM analysis (Goodman et al., 2014) as well as other structures (Bigalke et al., 2019; Li et al., 2019). There are two major interfaces between zRET^ECD^ and its ligand-co-receptor at opposing ends of zRET^ECD^, each is well defined in the cryo-EM map with sidechain level information (Figure 2D). The dominant interaction is between zCLD(1-3) and GFRα1^D3^ (defined hereafter as the site 1), with a key second site between zCRD and a concave surface presented by the GDNF dimer and a loop from GFRα1 (defined hereafter as site 2) (Figure 2D). Site 2 shows a close equivalence to the “low” affinity TGFβ receptor I binding site for TGF-β (Groppe et al., 2008; Kirsch et al., 2000) and also used by other TGF-β superfamily ligands (Hinck et al., 2016).

Site 1 on zRET involves elements from the CLD(1-2) clamshell structure and the CLD(2-3) calcium-binding region (Figure 2D). Both regions engage the zGFRα1 domain D3 (zGFRα1^D3^) close to helix α4, its preceding loop and helix α1 of domain D2. Together these elements form a wedge-shape surface to access the calcium-binding region of zRET^CLD(2-3)^. This interface covers a total area of 846 Å^2^ and comprises both hydrophilic and electrostatic interactions as calculated by PDBePISA (Krissinel and Henrick, 2007). The isolated CLD2-β1 strand bridges between the CLD1-CLD2 interface, running antiparallel to the zGFRα1^D3^ helix α4. Hydrophilic sidechains from helix α4 interact with CLD2-β1 mainchain as well as two proximal strands; CLD1-β1 and CLD1-β7 (Figure 2D). The sidechain from R330 of zGFRα1^D3^ helix α4, lies close to the mainchain carbonyl of I157 from CLD2-β1 and the sidechain of E326 is positioned near the sidechains of N247 and Y249 (hydroxyl). The loop preceding helix α4 of zGFRα1a^D3^ is anchored between the CLD3-β2-β3-loop and the CLD3-β4-β5-loop; mainchain-mainchain interactions form between P290 from the CLD3-β2-β3-loop and S321 of zGFRα1a^D1-3^ (Figure 2D). The mainchain of N323 from the loop preceding α4 of zGFRα1a^D1-3^ appears to interact with the guanidinium head of R333 from CLD3-β4, and the sidechain of N323 interacts with the mainchain of D250 at the calcium-binding site (Figure 2D).

Site 2 interaction involves the zRET^CRD^ and a concave “saddle” shaped surface formed by both protomers of the zGDNF^mat.^ dimer and a loop from zGFRα1a^D2^ (Figure 2D). This is in agreement with our previous assignment of this site as a “shared” site (Goodman et al., 2014) The interface is mainly hydrophobic in character and has a surface area of 598 Å^2^. The surface contains three main elements; a ß-turn from zGFRα1^D2^ centred on R180, residues 156-159 (LGYR) and residues 222-224 (HTL) from the fingers of one GDNF protomer (GDNF1) and residues 176-179 (DATN) with the “heel” helix of the second protomer (GDNF2). These residues engage G588 and Y589 from the CRD-β3-β4-loop (Figure 2D) and make Van der Waal’s contacts the I546 sidechain from CRD-β1-β2-loop (Figure 2D). A hydrophobic interaction between I586 from the CRD-β3-β4-loop and the T179 from the loop preceding the zGDNF2 “heel” (Figure 2D). The remaining contacts are mainly hydrophilic in nature between the heel of GDNF2 and the CRD. From the heel of zGDNF2; N180^GDNF^ interfaces with the amide of G587, and K182 of GDNF2 interacts with E613. This contact is consistent with the absence of a crosslink in the XL-MS data (Supplementary Figure 14). The zRET^CRD^ ß5-ß6 beta-turn is 2aa shorter than hRET^CRD^ allowing it to engage amino-terminal residues 138-140 of zGDNF2 with a likely salt bridge between E607 and R140. Also H222 from zGDNF1 is likely to contact E590, (equivalent to E595 in human RET, a known Hirschprung’s (HSCR) mutation site (So et al., 2011).

Two further contacts with zRET are indicated but are less well defined in the map. A limited interface between zRET^CLD1^ and GFRα1^D1^ is evident but this region shows lower resolution features than the rest of the cryo-EM map (Supplementary Figure 11). Nevertheless, the map allows zRET^CLD1^ and GFRα1^D1^ domains to be placed and the interaction is very similar to that observed for the NRTN-GFRα2^D2^ structure (Li et al., 2019). Second, residues immediately after the CRD from residue 615 to 627 are poorly ordered. This acidic stretch includes 12 residues likely to pass beneath the highly basic GDNF ligand (pI of 9.3 for mature zGDNF) before entering the plasma membrane. The final residue in RET^ECD^ observed is P617 which is separated by a distance of 40.9 Å from the dimer equivalent residue. A lower map contour shows density for these residues beneath the GDNF molecular 2-fold axis (view shown in Figure 2D) consistent with density seen for NRTN-GFRα2-RET (Bigalke et al., 2019).

### Clade-specific features influence ligand binding affinity

Comparison of site 1 of zRET in both the crystal and cryo-EM structure reveals differences in the conformation of residues 288-298 from a CLD3 loop (Figure 3A). In the absence of ligand, this loop packs against CLD3 core (loop “down” position) interacting with the β4 strand. In the presence of ligand this loop forms a central part of the interface with zGFRα1a^D3^ and is repositioned upwards (loop-”up”) towards the calcium ions and engages L247 of helix α1 of zGFRα1a^D2^ (Figure 3A). No equivalent interaction is observed for the human RET CLD3 structure (Figure 3B). The cryo-EM map clearly shows zGFRα1a^D3^ sidechain contacts with Y292 and how this residue shifts substantially relative its unliganded position (Figure 3C). This movement of 19.2Å (hydroxyl-hydroxyl) or 7.6Å (Cα-Cα) also results in main chain amides from P290 and V291 of CLD3-β2-β3-loop lying close to the mainchain carbonyl of S320 from zGFRα1a^D3^, forming a pseudo-β-sheet interaction (Figure 3A).

In view of the critical role of this loop in the zRET co-receptor recognition, it is surprising that loop CLD3-β2-β3 contains an “indel” of two extra amino acids Y292 and P293, unique to lower vertebrates (Figure 3D, Supplementary Figure 16). The equivalent shorter loop in human RET adopts a helical turn connecting the two β-strands (Figure 3C) (Li et al., 2019). To probe the contribution of the CLD3-β2-β3-loop to zGDNF-zGFRα1a binding, we truncated the residues P290-Q296 to AAG and assessed its ligand binding properties by microscale thermophoresis (MST). Surprisingly loop truncation improved binding affinity for ligand-co-receptor by 5-fold compared to wild type zRET^ECD^, with a dissociation constant (K_D_) of 18 nM (± 5 nM) compared to 90 nM (± 15 nM) for wild type zRET (Figure 3E). This gain-of-function increase in affinity implies either that higher vertebrates RET^ECD^ have a higher affinity for ligand than their lower vertebrate counterparts or that the loop contributes to an auto-inhibitory function in lower vertebrates. Taken together, our structural results show an unexpected conformational change in a clade-specific loop forming part of the high affinity ligand binding site proximal to the CLD(2-3) calcium sites.

Comparisons of interfaces within ternary RET complexes either between species (human and zebrafish GDNF-GFRα1) or paralogues (Neurturin-GFRα2 and GDF15-GFRAL) reveal considerable variation in contacts at site 1 and nearly identical contacts at site 2. This translates into a substantial variation in the size of these interfaces (Supplementary Table 3). One contributing factor to these variations is the additional contacts seen between helix α1 of zGFRα1^D2^ and residues 288-298 of zRET. Another example is GFRAL makes multiple additional contacts through residues 247-266, centered on the disulfide C252-C258, to engage residues flanking the beta-hairpin at Y76/R77 and R144/Y146 on CLD1 ß7-strand. Both elements are unique to higher vertebrate RET and contribute to the ligand-free RET dimer interface (Kjær et al., 2010; Li et al., 2019).

Comparison of all available liganded RET^ECD^ structures at site 2 consistently show a spacing of 44.2-47.0Å between each pair of CRD C-termini (measured at residue E613/620 in zRET/hRET) within a RET dimer (Figure 4A-C). This suggest a stringent requirement for CRD spacing to couple the transmembrane and intracellular modules. We note this distance is defined by the geometric length of a GFL ligand dimer and the position of the CRD relative to the dyad-axis of GDNF, presumed to sit above the RET transmembrane region.

### Structure-function analysis of zRET-GDNF-GFRα1a interaction sites

To probe the importance of each of the zRET interaction sites on ligand-complex assembly, mutations in the GDNF co-receptor at site 1 or GDNF at site 2 were selected. Our rationale was that given the composite binding site on RET, it would be easier to define the contribution of residues in the ligand or co-receptor. To guide mutant selection, surface residue heat maps, based on residue type and percentage similarity, were analysed for GFRα co-receptor subunit using alignments of homologues and paralogues from higher and lower vertebrates (Figure 5A and Supplementary Figure 17). The loop-helix α4 element of zGFRα1a^D3^ contributes residues N323, E326 and E327 to the RET-co-receptor interface and are present in most GFR sequences. These residues were mutated to alanine, both individually and as a triple-mutant. Using solution-based microscale thermophoresis (MST), affinity measurements of zGDNF^mat.^ _WT_ -zGFR1α1a^D1-3^ _N323A_ and zGDNF^mat.^ _WT_ -zGFR1α1a^D1-3^_E326A,E327A_ complexes binding to fluorescently-labelled zRET^ECD^ indicated only a modest impact, with a 2-fold decrease in affinity of E326A-E327A, corresponding to a K_d_ of 0.17 µM ± 0.039 µM vs 0.090 µM ± 0.015 µM for zGDNF ^mat.^ _WT_ -zGFR1α1a^D1-3^ _WT_ (Figure 5B). However, when combined as a triple mutation, zGDNF_wt_-zGFR1α1a^D1-3^ _N323A,E326A,E327A_, a 25-fold reduction in affinity was observed, (K_d_ of 2.35 µM ± 0.653 µM) (Figure 5B).

To probe the contribution of site 2 interface residues (Figure 5C and Supplementary Figure 18), residues L156, Y158, L224 and E220/H222 of zGDNF^mat.^ were selected for mutation to alanine and prepared using insect cells co-expressed with wild type zGFRα1a^D1-3^. The L224A and E220A/H222A mutations adversely affected the expression of zGDNF^mat.^ and could not be evaluated. MST was used to test the affinity of zGDNF^mat.^ _L156A_ -zGFRα1a^D1-3^ and zGDNF^mat.^ _WT_ -zGFRα1a^D1-3^ _WT_ towards zRET^ECD^. A 2-fold decrease in affinity observed for zGDNF^mat.^_Y158A_ towards zRET^ECD^, whereas no substantial loss in affinity was observed for zGDNF^mat.^_L156A_ (Figure 5D). We interpret the minimal effect of these mutations to zGDNF^mat.^ on zRET^ECD^ binding is indicative of a low affinity interaction site relative the zCLD(1-3)-zGFRα1a^D3^ site 1. Taken together, the data for zRET loop deletion and targeted zGFR1α and zGDNF mutations point to Site 1 being the dominant high affinity binding site despite both sites being required for ternary complex assembly.

### Different D1 domain orientation between GDNF and GDF15 co-receptor complexes

In the zRGα1a cryo-EM structure, the GFRα1^D1^ domain packs against GFRα1^D3^ using a linker with a conserved SPYE motif that is retained in all co-receptor sequences except GFRAL (Figure 6A). We therefore hypothesised that GFRAL^D1^ may require different contacts with RET through a distinctive D1-D2 linker sequence. To explore this possibility, a ternary complex was assembled comprising the <u>hR</u>ET^ECD^, hGDF<u>15</u>^mat.^ (hGDF15^195-380^) and hGFR<u>AL</u>^D1-3^ (hGFRAL^18-318^) (referred to hereafter as hR15AL) (Figure 6B) and cross-linked using GraFix to stabilise the complex (Supplementary Figure 19). A low-resolution negative stain envelope was produced with a total of 6519 particles with C2 symmetry averaging applied (Figure 6C, Supplementary Figure 19). While the overall shape of the envelope is similar to that of the zRGα1a map with a winged figure-of-eight appearance, it was evident that the wings are at a more acute angle to one another than in the zRGα1a cryo-EM map corresponding to a more “upright” hR15AL complex than the zRGα1a complex (Figure 6C).

Docking the recently published hRET^ECD^GDF15^mat.^GFRAL^129-318^ cryo-EM structure (PDB 6Q2J)(Li et al., 2019) into the low-resolution envelope corroborated initial observations of a more acute angle of both hRET copies compared to zGDNF-GFRα1a. It also revealed substantial density not accounted for by the fitted model, located beneath CLD(1-2) and flanking domain 2 and 3 (D2 and D3) of GFRAL (Figure 6C). The lack of domain 1 (D1) in the fitted model indicates that the area of unoccupied density is most likely the location of GFRAL^D1^ (Figure 5C). Such a position is in marked contrast to zGFRα1a^D1^ in zRGα1a (Figure 6D). This indicates a substantial plasticity in GFRAL as the most divergent GFR family member, explaining its lack of sequence conservation within the D1-D2 linker. It also emphasises further the ability of RET^ECD^ to accommodate a variety of ligand-co-receptors geometries from the flatter ARTN-GFRα3 to the upright GDF15-GFRAL complex, as shown by Li and coworkers (Li et al., 2019). Further studies are required to map in detail the additional interactions provided by GFRAL^D1^ to bind RET. We conclude that plasticity is not only evident within RET^ECD^ in accepting different GFL ligand-co-receptor geometries but also points to different roles for domain D1 between paralogues.

### Linear arrays of RET^ECD^-GDNF-GFRα1a observed on cryo-EM grids

Cryo-EM micrographs of the non-crosslinked sample of zRGα1a revealed significant orientation bias of the zRGα1a particles, with a single predominant orientation observed (Figure 7A). Upon closer inspection using RELION particle reposition (Zivanov et al., 2018) a dominant interaction between zRGα1a particles was observed throughout the grids resulting in linear arrays of complexes (Figure 7A). We analysed 3756 randomly picked particles from 14 micrographs. Using an interparticle distance of 214.2 Å (170 pixels) from the centroid of one particle to the centroid of neighbouring particles (x, y coordinates from the star file) 4132 particle pairs were defined. A 3D surface distribution plot of the difference in psi angles (Δψ) for pairs of particles against the distance between their centroids was calculated (Figure 7B), the psi (ψ) angles are generated in RELION 2D classification (Kimanius et al., 2016; Zivanov et al., 2018). An error of 3 ° exists within the plot due to the angular sampling value used during 2D classification. The 3D plot revealed that particles at a distance of 181 ± 3 Å from one another have an average ψ angle difference of 4.5 ± 2.3 °, using a minimal frequency of ψ angle difference to average distance of the more than 0.5 (Figure 7B). The recurrent and repetitive nature of this end-to-end contact for neighbouring particle pairs was further captured in a 2D class average, which used 1194 particle pairs (2388 individual particles) (Figure 7C).

Using the information gathered from the particle pair analysis, two copies of the zRGα1a complex structure were aligned with an inter-particle distance ∼180 Å apart and an angle of 4.5 ° between the two copies (Figure 7D). Observations of both the single particle as well as the 2D class averages generated for a pair of zRGα1a complexes show that the two wings of the figure-of-eight structure do not appear to be symmetrical, with a slightly more acute angle appearing between zGFRα1a and zGDNF on the sides in contact with one another in the neighbouring particles (Figure 7A and 7C). The inter-particle interaction site observed on cryo-EM grids lies on a predominantly hydrophobic surface of CLD2, comprising V192, V193, V198, P199, F200, V202 and M230 (Figure 7D, 7E and 7F). This hydrophobic patch is conserved between lower and higher vertebrates and is flanked by both basic (R232) and acidic clusters (D203, D204 and E239) that reciprocally neutralise each other across the zRGα1a-zRGα1a interface (Figure 7G). We note a highly conserved glycosylation site at N185 of CLD2 (found in both higher and lower vertebrates) is situated on the periphery of the multimerisation interface (Supplementary Figure 20). In a linear array context, this glycan could potentially interact with calcium ion Ca-1 (near CLD(2-3) junction) of an adjacent ternary complex to complete its coordination shell in *trans*, displacing the loosely-bound waters found in the zCLD(1-4)^red.sug^ structure. Further analyses are required to demonstrate a functional role for this multimeric interaction for full-length RET in a cellular context. Nevertheless, the high sequence conservation within the interface points to an important role beyond ligand-co-receptor interaction.

## Discussion

Here we establish principles for understanding the assembly of RET ligand-co-receptor complexes. We rationalise how RET can accept of a range of activating GFL-co-receptor binary complexes through conformational adaptations between RET and co-receptor. By using crystallography and cryo-EM we define the architecture and ligand-recognition properties of zebrafish RET^ECD^ and compare this to the human RET^ECD^. Our results provide four main insights; (1) there is conformational flexibility at the CLD(2-3) interface of RET^ECD^ that contribute to optimised adaptations at the co-receptor binding site (2) there are conformational differences between unliganded and liganded RET centred on a clade-specific RET loop (3) a strict spatial separation of RET^ECD^ C-termini within the ternary complex is imposed by each CRD interaction with GFL dimer (4) differences in co-receptor engagement and putative higher-order multimers of ligand-bound RET suggest divergent interactions at each level of receptor engagement.

Previous insights into GFL-co-receptor recognition from negative stain and cryo-electron microscopy have revealed two main contact sites in RET (Bigalke et al., 2019; Goodman et al., 2014; Li et al., 2019). These structures explained why an intact calcium-loaded RET^ECD^ is required for GDNF-GFRα1 binding as the GFRα1^D3^ loop-helix α4/GFRα1^D2^ helix α1 wedge targets the calcium-dependent CLD(2-3) hinge while the GDNF dimer targets the CRD. The GFRα1 wedge may act as a sensor for calcium bound to RET implicating calcium not only in promoting RET folding but also proper recognition by co-receptor for signalling (Nozaki et al., 1998). The RET^CRD^ interaction with both protomers of a GDNF dimer is directly equivalent to the binding site of “low affinity” TGF-β/BMP family of type 1 receptors for TGF-β (“knuckles” and “thumb”) (Hinck et al., 2016). Whereas the TGF-β “fingers” engage the “high affinity” TGF-β receptor, equivalent to GFRα co-receptors binding to GFL “fingers”.

Several studies identified a role for site 1 contacts close to N323 in RET ternary complex formation (Goodman et al. 2014; Bigalke et al. 2019; Li et al. 2019). The strikingly distinctive contacts made by different GFRα homologues at site 1 suggest conformational adaptions enable the recognition of multiple GFR co-receptors and different GFR_2_GFL_2_ geometries. Our findings suggest engagement of ligand-co-receptor through the calcium-dependent CLD(2-3) hinge promotes a remodelling of the lower-vertebrate-specific loop and may precede site 2 RET^CRD^ engagement. This could involve either a pre-assembled RET-GFRα complex or presentation of GFRα after dimerization by GFL, prior to RET^CRD^ interaction. We show here from substitution of zGDNF residues in site 2 (L156A and Y158A) that these contacts do not appear to play a dominant role in ternary complex assembly. This contrasts with a study showing mutation of Y119 to E in Neurturin (equivalent to Y158 of zGDNF) disrupted ternary complex formation and signalling (Bigalke et al. 2019). Given the analogous RET^CRD^ contacts at site 2 for each GFL dimer are proximal to the RET transmembrane segment, suggests an organising role for signal transduction in addition to ligand recognition.

The D1 domain is missing from structures for GDNF-GFRα1 and GDF15-GFRAL, but had been observed for NRTN-GFRα2 alone or bound to hRET^ECD^ (Bigalke et al., 2019; Li et al., 2019). We were able to place the GFRα1 domain D1 adjacent to zRET^CLD1^, consistent with previous negative stain EM models (Goodman et al., 2014). As previously shown the D1 proximity to RET^CLD1^ is not essential for ternary complex formation. We present evidence from a low-resolution map density consistent with a quite different contact position for the GFRAL D1 domain adjacent to GFRAL D2 and D3, on the outside/ of RET, underneath the “wings” rather than within the hRET^ECD^ ternary complex. This explains the absence of the otherwise conserved SPYE motif common to GFRα1/2/3 motifs at the D1 and D3 interface. This position for the GFRAL D1 domain arises from a more upright position for GFRAL observed than GDNF-GFRα1 complexes (Li et al., 2019). While the functional significance of this difference is yet to be understood, it could impact on the assembly of higher order multimers such as those observed for zRGα1a.

We and others have provided structural evidence for RET dimers in the absence of ligand-co-receptor through a CLD1-2 dimer interface involving R77 and R144 sidechains (Kjær et al., 2010; Li et al., 2019). Here we describe linear arrays of zGDNF-zGFRα1-zRET^ECD^ complexes on cryo-EM grids mediated by a hydrophobic patch on an exposed part of CLD2 in the ternary complex. We also observe “stacks” of these linear arrays similar to a dimer of dimers interface reported for hRET^ECD^ for a hNRTN-hGFRα2-hRET^ECD^ ternary complex (data not shown; Li et al., 2019). This interface was reported to influence the rate of receptor endocytosis. These findings suggest that a signalling-competent RET^ECD^ conformation is likely to involve higher order multimers consistent with findings for other RTKs such as EphR (Seiradake et al., 2010) EGFR (Needham et al., 2016) and DDR1 (Corcoran et al., 2019) RTKs. Therefore, a crucial aspect of receptor activation beyond the positioning of the RET transmembrane regions within a dimeric assembly is their arrangement within higher order clusters.

In summary, this study reveals several under-appreciated aspects of GFL-co-receptor binding to RET including receptor flexibility, clade-specific adaptations and conformational changes. All these features reveal a substantial tolerance within RET to accommodate different GFL-co-receptors using a flexible arm. It also suggests a key requirement for coupling ligand-binding to RET activation is a strict spatial separation between CRD C-termini within RET dimers imposed by the geometric dimensions of each GDNF family ligand. The next challenge will be to visualise such arrangements of a full-length RET multimer in a membrane context and to use this knowledge in the design of both antagonist and agonist biologicals that with therapeutic utility.

## Material and Methods

### Zebrafish RET CLD(1-4) expression and purification

Zebrafish RET^1-504^ (zCLD(1-4)^red.sug.^) was designed with glycosylation site mutations N259Q, N308Q, N390Q and N433Q to aid in crystallisation. This construct was cloned into a pBacPAK-LL-vector together with a 3C-cleavable C-terminal Protein A tag. A recombinant baculovirus was prepared using the FlashBAC system (2B Scientific). For protein production, SF21 cells were grown to a cell density of 1×10^6^ and incubated with recombinant virus for 112 hours at 27 °C. The media was harvested and incubated with IgG sepharose (Sigma), with 1 ml of resin slurry to 1 l of media, whilst rolling at 4 °C for 18 hrs. The resin was recovered and washed with 5 column volumes (c.v.) of 20 mM Tris (pH 7.5), 200 mM NaCl, 1 mM CaCl_2_ then incubated with 1:50 (w/w) PreScission Protease (GE Healthcare) for 18 hrs at 4 °C. The eluted zCLD(1-4)^red.sug.^ was further purified using a SuperDex 200 (GE Healthcare).

### zCLD(1-4)^red.sug.^ crystallisation and X-ray data collection

The purified zCLD(1-4)^red.sug.^ was concentrated to 12 mg/ml. Vapour diffusion drops were set up with 2 µl of protein and 2 µl of precipitant; 50 mM MES (pH 6.2), 31.5 % PEG MME 350 (v/v), against 90 µl of precipitant. After 24 hrs of equilibration seeding was performed using Crystal probe (Hampton Scientific). Crystals grew over 14 days at which point they were harvested and flash frozen in liquid nitrogen.

### zCLD(1-4)^red.sug.^ x-ray data processing and structure determination

Data from these crystals was collected at the Diamond Light Source, initially on beamline I04 and finally on beamline I03. The data was processed with XIA2 utilising DIALS (Winter et al., 2018), before further processing through STARANISO (Tickle et al., 2018) for anisotropy correction to give a 2.08 Å dataset (cut to 2.20 Å for refinement owing to low completeness in the outer shells). Crystals belonged to the triclinic space group P1 with cell dimensions a=51.2 Å, b=70.5 Å, c=105.4 Å, a=105.4°,b=100.9°,c=100.3°. Molecular replacement was used as implemented in PHASER (McCoy et al., 2007) to initially locate two copies of CLD1-2 (PDB code 2XU). The positions of the two associated copies of CLD4 were then determined, utilising an ensemble of the following seven models (superposed by secondary structure matching in COOT): 1L3W (resid A 6-99)(Boggon et al., 2002), 1NCI (resid A 6-99)(Shapiro et al., 1995), 1OP4 (resid A 40-123)(Koch et al., 2004), 4ZPL (resid A 206-314)(Rubinstein et al., 2015), 4ZPM (resid B 207-317)(Rubinstein et al., 2015), 4ZPO (resid A 205-311)(Rubinstein et al., 2015) and 4ZPS (resid A 205-313)(Rubinstein et al., 2015). Initial refinement with PHENIX.REFINE was followed by automated model building with PHENIX.AUTOBUILD (Terwilliger et al., 2007) which completed most of the two polypeptide chains present. Cycles of manual model building with COOT and refinement with PHENIX.REFINE (Afonine et al., 2012) followed. Insect cell glycosylation sites were modelled and checked using PRIVATEER (Agirre et al., 2015), with additional libraries, describing the linkages between monomers generated, and used initially in refinement to maintain a reasonable geometry.

### zGDNF^mat.^-zGFRα1^D1-3^ expression and purification

Baculoviruses for zebrafish GFRα1a^1-353^ (zGFRα1a^D1-3^) and zebrafish GDNF^135-235^ (zGDNF^mat.^) were produced using the pBacPAK-LL-zGFRα1a^D1-3^-3C-ProteinA construct and the pBacPAK-LL-melittin-zGDNF^mat.^-3C-ProteinA respectively and FlashBac viral DNA (2B Scientific) using standard protocols (2B Scientific). Recombinant baculoviruses producing either zGDNF^mat.^ or zGFRα1^D1-3^ with a 3C cleavable protein A tag were expressed using SF21 insect cells. Briefly, 6 x 500 ml flasks of SF21 cells grown to a cell density of 1 x 10^6^ in SFIII media, were infected with 10 ml of the zGDNF_mat._ baculovirus stock and 2 ml of the zGFRα1^D1-3^ baculovirus stock for 86 hrs. Cells were pelleted at 3500 xg and the media containing the secreted 2:2 zGFRα1a^D1-3^-zGDNF^mat.^ complex was pooled. A 1 ml slurry of IgG sepharose resin (GE Healthcare) was added to 1 l of media and incubated at 4 °C for 18 hrs. The resin was recovered and washed with 5 column volumes of 20 mM Tris (pH 7.0), 150 mM NaCl and 1 mM CaCl_2_, resuspended 2 column volumes of the same buffer and incubated with GST-3C (20 µl at 8 mg/ml) for 16 hours. zGDNF^mat^ -zGFRα1^D1-3^ was further purified using size exclusion chromatography using a Superdex 200 (16/600) (GE Healthcare) in 20 mM Tris (pH 7.0), 100 mM NaCl and 1 mM CaCl_2_.

### zGDNF^mat.^-zGFRα1^D1-3^ crystallisation and structure determination

Purified zGDNF^mat.^-zGFRα1^D1-3^ was concentrated to 2.5 mg/ml. 100 nl of protein was dispensed with 100 nl of precipitant in the sitting well trays (MRC-2 drop trays) which comprised 100 mM Tris (pH 8.0), 5 % (w/v) PEG 20,000, 3.7 % (v/v) Methyl cyanide and 100 mM NaCl. A volume of 90 µl of precipitant solution was dispensed into the well, the trays were incubated at 22 °C. Crystals of zGDNF^mat.^-zGFRα1^D1-3^ formed after 30 days. Crystals were harvested after 55 days and frozen in liquid N_2_ with 30% ethylene glycol used as a cryo-protectant. Data was collected on I04 at Diamond using PILATUS 6M Prosport+ detector. The X-ray diffraction data collected was reduced and integrated using DIALS (Clabbers et al., 2018; Waterman et al., 2016; Winter et al., 2018) at Diamond. The data was phased using PHASER (McCoy et al., 2007) molecular replacement in CCP4 (1994; Winn et al., 2011) using the human GDNF-GFRα1 (PDB 3FUB)(Rubinstein et al., 2015). Model refinement was performed using COOT (Emsley and Cowtan, 2004; Emsley et al., 2010) and PHENIX.REFINE (Adams et al., 2010; Afonine et al., 2012) against the dataset that was reduced and integrated using the STARANISO (Tickle et al., 2018) at a resolution of 2.2Å. Glycosylation sites were validated using PRIVATEER (Agirre et al., 2015).

### zRET^ECD^-zGDNF^mat.^-zGFRα1^D1-3^- (zRGα1a) complex expression and purification

A recombinant baculovirus was prepared to produce zRET^ECD^ (residues 1-626) using the pBacPAK-LL-zRET^ECD^-3C-Protein A construct and FlashBac viral DNA (2B Scientific) using standard protocols and as described above. To produce zRET^ECD^, SF21 insect cells grown using SFIII media in 6×500 ml flasks to a cell density of 1×10^6^ were then infected with 2 ml of the baculovirus that contained zRET^ECD^ for 86 hrs at 27 °C. Cells were pelleted at 3500 g and the media containing secreted zRET^ECD^ was pooled and 1 ml of IgG sepharose resin (GE Healthcare) was added to 1 l of media and incubated at 4 °C for 18 hrs. The resin was recovered and washed with 5 column volumes of 20 mM Tris (pH 7.0), 150 mM NaCl and 1 mM CaCl_2_, resuspended in 2 column volumes of the same buffer. Purified 2:2 zGFRα1a^D1-3^- zGDNF^mat.^ complex was then added directly. The sample was incubated for 45 min at 4 °C. The resin with the zRGα1a complex was then recovered and washed with 5 c.v. of 20 mM Tris (pH 7.0), 150 mM NaCl and 1 mM CaCl_2_ buffer, resuspended in 2 column volumes of buffer and incubated with GST-3C (20 µl at 8 mg/ml) for 18 hours at 4 °C. The eluted zRGα1a complex was further purified using size exclusion chromatography using a Superdex 200 (16/600) (GE Healthcare) in 20 mM HEPES (pH 7.0), 150 mM, NaCl and 1 mM CaCl_2_.

To prepare a cross-linked sample, 100 µl of purified zRGα1a (4 mg/ml) was applied on top of a 5-20 % (w/v) sucrose gradient which contained a 0-0.1 % (v/v) glutaraldehyde gradient, the gradient was buffered with 20 mM HEPES (pH 7.0), 150 mM NaCl and 1 mM CaCl_2_. Ultracentrifugation was performed at 33,000 r.p.m (SW55 rotor) for 16 hours at 4 °C. The sucrose gradient was fractionated in 125 µl fractions, the glutaraldehyde was quenched with 1 M Tris (pH 7.0), to a final concentration 100 mM. The fractions that contained cross-linked zRGα1a were pooled and further purified by Superdex200inc 10/300 (GE Healthcare) in a buffer of 20 mM Tris (pH 7.0), 150 mM NaCl and 1 mM CaCl_2_, in order to remove the sucrose from the crosslinked zRGα1a complex.

### zRGα1a cryo-electron microscopy sample preparation

To prepare cryo-EM grids, 1.2/1.3 300 mesh Cu Quantifoil™ grids 300 mesh grids were glow discharged using 45 mA for 30 s using a Quorum Emitech K100X. For the untilted dataset (Dataset 1), 4 µl of crosslinked zRGα1a sample, at 0.1 mg/ml, was applied to the grids, using a Vitrobot Mark IV (Thermo Fisher) with the parameters; 90 s wait time, 5 s blot time at 22 °C with 100 % humidity. The same glow discharge parameters were used for the grids for the tilted dataset (dataset 2), 4 ul was applied to the grid at 4 °C and a 20 s wait with 3 s blot time under 100 % humidity. For the non-crosslinked zRGα1a sample, the same glow discharge parameters were used for 1.2/1.3 300 mesh Cu Quantifoil™ grids 300 mesh grids. 4 µl of non-crosslinked zRGα1a at 0.1 mg/ml was applied to the grids with the same parameters as those used for the grids prepared for dataset 1, these grids were used for dataset 3.

### Cryo-EM data acquisition: Datasets 1 to 3

Frozen-hydrated grids of the crosslinked zRGα1a sample were imaged on a Titan Krios electron microscope (Thermo Fisher) operating at 300 kV at the Francis Crick Institute. Movies were captured on a BioQuantum K2 detector (Gatan) in counting mode at 1.08 Å/pixel and with an energy filter slit width of 20 eV. Dataset 1 was collected with a 0° tilt angle, a defocus range of 1.4-3.5 µm and comprised a total of 6105 movies. For dataset 2, 6375 movies were collected in total using a tilt angle of 30° and the same defocus range used for dataset 1. Movies from datasets 1 and 2 had an exposure of 1.62 e^-^/Å^2^ per frame for a total electron exposure of 48.6 e^-^/Å^2^. The dose rate was 6.4 e^-^/pixel/sec and exposure time was 9 seconds/movie. For dataset 3, frozen-hydrated grids of non-crosslinked zRGα1a were collected on a Talos Arctica microscope (Thermo Fisher) operating at 200 kV at the Francis Crick Institute. A total of 1705 movies were captured on a Falcon 3 detector in integrating mode at 1.26 Å/pix and a defocus range of 1.5-3.0 µm. Movies from dataset 3 had an exposure of 6.07 e^-^/Å^2^ per frame which led to a total exposure of 60.66 e^-^/Å^2^. All datasets were collected using EPU version 1.9.0 (Thermo Fisher).

### Cryo-EM data processing of crosslinked zRGα1a (dataset 1)

MotionCorr2 (Zheng et al., 2017) was used to correct for motion in the movie frames in Scipion 1.2 (de la Rosa-Trevín et al., 2016). The contrast transfer function was estimated using CTFfind4.1(Rohou and Grigorieff, 2015). 5855 micrographs were selected from dataset 1 and initial particle picking was performed with RELION-2.1 manual picking, 4899 particles were extracted with RELION-2.1 (Zivanov et al., 2018) particle extract function (de la Rosa-Trevín et al., 2016) with a box size of 340 and binned two-fold. 2D classification was performed using RELION 2D classification, with 20 initial classes. Six classes were used to pick a subset of 3000 micrographs using RELION-2.1 autopicking in Scipion 1.2, giving 638,000 particles with box size 340, binned 2 fold. These were classified using 2D classification in RELION-2.1. Twelve classes were selected for picking using Gautomatch [K. Zhang, MRC LMB (www.mrc-lmb.cam.ac.uk/kzhang/)] to pick 2,424,600 particles, which were extracted with a box size of 340 pixels and binned 2-fold using RELION-2.1 2D class averaging was performed in CryoSPARC-2(Punjani et al., 2017) leading to 1,156,517 particles which were extracted using RELION-2.1 (Kimanius et al., 2016; Scheres, 2012; Zivanov et al., 2018) with a box size of 320 pixels.

### Cryo-EM data processing of crosslinked zRGα1a (tilted dataset 2)

Dataset 2 was processed and corrected for motion correction and CTF estimation as described above. A total of 4848 micrographs were used to pick particles semi-automatically with Xmipp and 69,386 particles were extracted with a box size of 360 pixels using RELION-2.1 (Kimanius et al., 2016; Scheres, 2012) in Scipion1.2 (de la Rosa-Trevín et al., 2016). RELION-2.1 (Kimanius et al., 2016; Scheres, 2012; Zivanov et al., 2018) 2D classification was then performed, with subsequent picking using RELION automatic picking leading to 1,183,686 particles being extracted using RELION-2.1 (Kimanius et al., 2016; Scheres, 2012; Zivanov et al., 2018) with a box size of 340 binned 2-fold. Subsequent 2D classification in RELION-2.1 (Kimanius et al., 2016; Scheres, 2012; Zivanov et al., 2018) lead to 12 classes which were used by Gautomatch [K. Zhang, MRC LMB (www.mrc-lmb.cam.ac.uk/kzhang/)] to pick 1,393,023 particles. The particles were extracted with RELION-2.1 (Kimanius et al., 2016; Scheres, 2012; Zivanov et al., 2018) with a box size 320, 2 fold binned, were imported into CryoSPARC-2 (Punjani et al., 2017) and 2D classification generated 208,057 particles from 3175 micrographs. These particles were re-extracted with a box size of 320 and per-particle CTF estimation was performed using GCTF(Zhang, 2016).

### Combining and re-processing cryo-EM datasets 1 and 2 for crosslinked zRGα1a

Dataset 1 and 2 were combined and an initial 2D classification was performed in CryoSPARC-2 on the 1,364,574 particles(Afonine et al., 2018). Following this, 1,242,546 particles underwent two heterogeneous refinements using 5 classes with strict C2 symmetry applied in CryoSPARC-2 lead to a homogeneous refinement with 468,922 particles(Punjani et al., 2017). Once re-imported into Scipion1.2, RELION 2D class averaging was implemented to generate 364,158 and 22,358 particles from dataset 1 and dataset 2, resectively(Kimanius et al., 2016; Scheres, 2012). Particle polishing was performed in RELION-2.1 (Zivanov et al., 2018). Once imported into CryoSPARC-2, 2D class averaging removed any further particles, yielding 382,547 particles used for a homogeneous refinement followed by a non-uniform refinement with C2 symmetry applied. This final reconstruction gave a resolution of 3.3 Å as calculated using the ‘gold’ standard (FSC=0.143)(Kimanius et al., 2016; Scheres, 2012). Symmetry expansion was performed in RELION-2.1 and 3D-refinement with masking was performed with no symmetry applied (Kimanius et al., 2016; Scheres, 2012; Zivanov et al., 2018). Postprocessing in RELION-2.1 of the final symmetry expanded reconstruction with a resolution 3.5 Å (Supplementary Figure 6)(Kimanius et al., 2016; Scheres, 2012; Zivanov et al., 2018).

### Building the zRGα1a complex into the final cryo-EM map

To build a full ligand-co-receptor complex, the zGDNF^mat.^-zGFRα1^ΔD1^ crystal structure described here was used together with a homology model of domain D1 (zGFRα1^29-121^) generated by MODELLER from the GFRα2-neurturin crystal structure (PDB 5MR4) (Sandmark et al., 2018; Webb and Sali, 2016). For zRET, chain A of the CLD(1-4) module described here was used together with a CRD model generated with SwissPROT (Schwede et al., 2003) using the structure of hRET^ECD^ in complex with GFRα2-neurturin (PDB 6Q2O)(Li et al., 2019; Webb and Sali, 2016). The zGDNF-zGFRα1 and zRET^ECD^ structures were then docked into the symmetry expanded map using PHENIX (Adams et al., 2010). The model was refined against the sharpened map using PHENIX_REAL_SPACE_REFINE (Afonine et al., 2018) and manual model building and model refinement was done in COOT (Emsley and Cowtan, 2004; Emsley et al., 2010). The final symmetry expanded model was used to generate the 2:2:2 zRGα1a model, which was placed in the C2 averaged map using PHENIX (Adams et al., 2010) using PHENIX_REAL_SPACE_REFINE (Afonine et al., 2018). Glycosylation sites were validated using PRIVATEER (Agirre et al., 2015). Protein-protein interface areas were calculated using PDBePISA (Krissinel and Henrick, 2007).

### Cryo-EM data processing for a non-crosslinked zRGα1a sample (dataset 3)

MotionCorr2 (Zheng et al., 2017) was used to correct for motion in the movie frames in RELION-3 (Zivanov et al., 2018). The contrast transfer function was estimated using CTFfind4.1 (Rohou and Grigorieff, 2015). 384 micrographs were selected from and initial particle picking was performed with RELION-3 manual picking, 951 particles were extracted with RELION-3 (Zivanov et al., 2018) particles extract with a box size of 320 and binned 2 fold. 2D classification was performed using RELION 2D classification, with 10 initial classes(Kimanius et al., 2016; Scheres, 2012; Zivanov et al., 2018). One class, due to the orientation bias, was selected and used by RELION autopick to pick from a subset of 81 micrographs. This gave 19,715 particles picked and extracted with a box size of 320 pixels using RELION-3. These particles were sorted in RELION-3 and 15,519 were then were classified using RELION 2D classification. A total of 11070 particles were used from 81 micrographs to explore the linear particle arrays observed for the zRGα1a complex.

### Analysis of zRGα1a multimer formation on cryo-EM grids

Following 2D class averaging in RELION-3, the final 11070 particles were repositioned onto 81 micrographs collected from cryo-grids prepared from the non-crosslinked zRGα1a sample using RELION particle reposition. A Python script was written to extract the particle number, psi angle (ψ) and Cartesian coordinates of particle pairs from the 2D class average STAR file. Particle pairs were detected through analysing each single particle and locating surrounding particles within 214.2 Å (170 pixels), using their extracted Cartesian coordinates. A subset of 14 micrographs was used, where a total of 3756 individual particles lead to 4132 particle pairs. The distance between each particle pair was determined using the X and Y coordinates. The ψ angles were corrected to positive integers, and were permitted to be within the 180 ° range due to the C2 symmetry of the complex. The difference between the two positive ψ angles from the particle pairs (Δψ) was calculated as an absolute value. Distance between the particles and the Δψ between particle pairs was calculated and plotted on a 3D surface plot with the bins every 2 Å and every 2.6 °, respectively.

### Human RET^ECD^ expression and purification

A codon-optimised human RET^ECD^ (hRET^ECD^) cDNA encoding residues 1-635 followed by a TEV-cleavable Avi and C-tag was cloned into a pExpreS2.1 vector (ExpreS2ion Biotechnologies, Hørsholm, Denmark) with Zeocin resistance. A stable pool of S2 cells, secreting hRET^ECD^, was generated by transfecting 25 ml of S2 cells grown in Ex-Cell420 medium (Sigma) with 10 % (v/v) FBS at a density of 5×10^6^ cells/ml using 12.5 µg of DNA and 50 µl of ExpresS^2^-Insect TR (5×). Stably transfected cells were selected with 2 mg/ml Zeocin with repeated medium exchange. The culture was expanded to 1 litre in a 5L glass-flask and the supernatants collected after 7 days.

For purification, 1 ml of C-tag capture resin (ThermoFisher) was added to a cleared and filtered S2 supernatant and incubated for 18 hrs at 4 °C. The resin was pelleted and washed several times with PBS before eluting bound hRET^ECD^ by competition with PBS containing 200 µg/ml SEPEA peptide. At this point, the affinity and biotinylation tags were removed by digestion with TEV (a 1:10 ratio of TEV protease:RET). The purified hRET^ECD^ was further purified by size-exclusion using a Superdex200 10/300 with a 50 mM Tris (pH 7.5), 100 mM NaCl buffer.

### Human GDF15-GFRAL complex expression and purification

Both human GFRAL^21-352^ (referred to hereafter as hGFRAL^D1-3^) and hGDF15^198-308^ (referred to hereafter as hGDF15^mat^) were cloned into a pCEP vector with an N-terminal BM40 secretion sequence. The hGFRAL construct had a C-terminal 6 His tag. The constructs were co-transfected into Expi293 cells (Life Tech) using polyethylimine. The transfected cells were incubated in Freestyle media at 37 °C, 8 % CO_2_ with 125 rpm shaking. Conditioned media was harvested after 5 days, and Tris pH 8.0 and Imidazole added to a final concentration of 10 and 20 mM respectively. The media was incubated with Ni-NTA agarose beads whilst rolling at 4 °C for 2 hours. The beads were recovered and washed with 20 mM Tris (pH 7.4), 137 mM NaCl and the protein was eluted with 20 mM HEPES (pH 7.4), 137 mM NaCl and 500mM imidazole. The protein was concentrated to ∼5 mg/ml. This protein was further purified by Superdex 200 increase size exclusion chromatography in buffer 20 mM HEPES (pH 7.4), 137 mM NaCl to give a pure 2:2 GDF15-GFRAL complex.

### hRET^ECD^-hGDF15^mat.^-hGFRAL^D1-3^ (hR15AL) complex assembly and purification

An excess of purified hRET^ECD^ (300 µl, 1.1 mg/ml) was incubated with purified hGDF15-hGFRAL (300 µl, 0.75 mg/ml) for 1 hr whilst mixing at 4 °C in the presence of 10-fold excess heparin sulfate DP-10 (20 μM) (Iduron, UK). The hR15AL complex was further purified by size exclusion chromatography using a Superdex 200 increase in to 20 mM HEPES (pH 7.0), 150 mM NaCl and 1 mM CaCl_2_. For sample crosslinking, 100 µl of the hR15AL complex (0.75 mg/ml) was applied on top of a 5-20 % (w/v) sucrose gradient which contained a 0-0.1 % (v/v) glutaraldehyde gradient, the gradient was buffered with 20mM HEPES (pH 7.0), 150 mM NaCl and 1 mM CaCl_2_. Ultracentrifugation was performed at 33,000 rpm for 16 hours at 4 °C. The sucrose gradient was fractionated in 125 µl fractions, the glutaraldehyde was quenched with 1M Tris (pH 7.0), to a final concentration 100 mM. The fractions were assessed using SDS-PAGE and fractions that contained the complex were used for negative stain.

### hR15AL negative stain preparation, data acquisition and processing

Cu 200 mesh carbon coated grids were glow discharged under vacuum using 45 mA for 30 s. A sample of 4 µl of the crosslinked hR15AL undiluted from the GraFix column was applied to the charged grid and left for 30 s and the excess removed by blotting and placing the grid, sample side facing the solution, in 10 µl of 2 % (w/v) uranyl acetate solution in d.H_2_O, and blotting immediately twice, followed by placing the grid in the 3^rd^ 10 µl drop sample side facing down and leaving it in solution for 1 min, followed by a final blot until almost all the solution has been wicked off. The grid was then left to dry for 5 mins.

Micrographs were collected on a BMUltrascan 1000 2048×2048 CCD detector using a Tecnai Twin T12 (Thermo Fisher) at 120kV with a defocus range of 1-1.5 µm and with a 1 s exposure time. A total of 299 micrographs were collected and particles were picked using Xmipp (De la Rosa-Trevín et al., 2013) semi-automated picking, in Scipion1.2 (de la Rosa-Trevín et al., 2016). This gave 27,551 particles were extracted with RELION-2.0 particle extraction (Kimanius et al., 2016; Scheres, 2012). 2D class averaging was performed with RELION-2.0 (Kimanius et al., 2016; Scheres, 2012). The resulting 16,159 particles were used to generate an initial model using RELION 3D ab-initio model. 3D classifications with 5 classes were performed using RELION-2.0 3D classification(Kimanius et al., 2016; Scheres, 2012). 6519 particles were taken forward into the final reconstruction a resolution of 25.8 Å using RELION-2.0 3D refinement (Kimanius et al., 2016; Scheres, 2012). The data processing was done in Scipion1.2 (de la Rosa-Trevín et al., 2016).

### Microscale thermophoresis (MST) measurement of zRET^ECD^ binding affinity

MST measurements were performed at 25 °C in 20 mM HEPES (pH 7.0), 150 mM NaCl, 1 mM CaCl_2_ and 0.05 % (v/v) Tween-20 using a Nanotemper Monolith NT.115 (Nanotemper). To measure the affinity of zGFRα1^D1-3^-zGDNF^mat.^ towards zRET^ECD^; zRET^ECD^ was labelled with NHS-RED 2^nd^ generation dye (Amine Reactive) using the labelling kit (Nanotemper). A 1:1 serial dilution of unlabelled zGFRα1^D1-3^-zGDNF^mat.^ (WT and mutants) was performed. The samples were incubated with the labelled zRET^ECD^-NHS-RED (50 nM, fluorophore, 83.7 nM zRET^ECD^) for 10 mins at 22 °C. Hydrophobic treated capillaries were filled with the serially diluted samples (Nanotemper). The MST run was performed using a Monolith 1.115 with the LED power at 20% with a measurement time of 20 sec. To measure the affinity of zGFRα1^D1-3^-zGDNF^mat.^ towards zRET^ECD^_P291-Q296;AAG_; zRET^ECD^ _P291-Q296;AAG_ was labelled with NHS-RED 2^nd^ generation dye (Amine Reactive) using the labelling kit (Nanotemper), and the procedure was carried out as above with zRET^ECD^ _P291-Q296;AAG_ -NHS-RED (50 nM, fluorophore, 80.7 nM zRET^ECD^).

### Surface conservation analysis and heatmaps for different GFL-GFR ligand-coreceptor pairs

The sequence for the globular domains of zGFRα1a (Uniprot Q98TT9) was aligned to hGFRα1 (Uniprot P56159), hGFRα2 (Uniprot O00451), hGFRα3 (Uniprot O60609), hGFRα4 (Uniprot Q9GZZ7), and hGFRAL (Uniprot Q6UXV0), using Clustal Omega.(Sievers et al., 2011) The sequence of the mature zGDNF (Uniprot Q98TU0) was aligned to hGDNF (Uniprot P39905), hNRTN (Uniprot Q99748), hARTN (Uniprot Q5T4W7), hPSPN (Uniprot O60542), and hGDF15 (Uniprot Q99988) using Clustal Omega.(Sievers et al., 2011) Using these alignments, residues were categorised based on residues type and a heat map generated and values mapped onto a surface representation on the zGFRα1a^D2-D3^. D1 was excluded from the analysis due to the major differences between each of the co-receptors; which is missing hGFRα4 and is located in a completely different position in hGFRAL. Each of the categories for residue type are as follows; the aromatic residues (F, W, and Y), the aliphatic residues (A, I, L, and V), residues containing an alcohol functional group (S and T), positively charged residues (R and K), negatively charged residues (D and E), and residues containing an amide bond in the side chain (N and Q), and C, G, H and M were counted individually. The sequence similarity was numbered from 0-1, 0 indicating no similarity at all and 1 indicating the residue type was identical between the GFR or GFL family members respectively. The values per residue in the sequence were used as B-factors for the structure and were represented as a surface colour coded with the highest residue similarity in red (1) through yellow (0.5) to white (0).

## Acknowledgements

We thank members of the McDonald laboratory for helpful discussions and comments on the manuscript. We thank Simone Kunzelmann for help with MST experiments. We thank Raffaella Carzaniga and Lucy Collinson for EM training and support. We thank Alessandro Costa and Tom Miller for their advice regarding the particle replacement analysis. We also wish to acknowledge the helpful advice and support of Peter Rosenthal on all aspects of cryo-electron microscopy. We thank Carlos Ibanez for critical reading. We would like to thank Diamond Light Source for beamtime (proposals mx13775) and the staff of beamlines I03 and I04 for their assistance with crystal testing and data collection. N.Q.M. acknowledges that this work was supported by the Francis Crick Institute, which receives its core funding from Cancer Research UK (FC001115), the UK Medical Research Council (FC001115) and the Wellcome Trust (FC001115).

## Author contributions

S.E.A. prepared the zRGα1a complex and carried out EM data processing, crystallised the GDNF-GFRα1a complex and performed the MST assays. A.P. phased the zCLD(1-4) X-ray crystal structure and built the structural model. P.P.K. optimised the CLD(1-4) crystals and obtained and cryo-cooled the final crystals. K.M.G. performed initial CLD(1-4) expression and crystallisation experiments and Agata.N. performed CLD(1-4) crystallisation optimisation experiments. Andrea.N. collected the zRGα1a Krios datasets and wrote the python script to detect particle pairs. Annabelle.B and S.K. prepared the hRET^ECD^. D.C.B. expressed and purified the hRET^ECD^-hGDF15-hGFRAL complex. C.P.E assisted in cryo-EM sample optimisation, EM data processing, and model refinement. S.E.A. crosslinked the hR15AL complex and collected and processed the EM data. Aaron.B performed the XL-MS zRGα1a experiments. F.M.H. expressed some of the zGFRα1 mutants used for the MST analysis. P.B.M. collected the native non-cross-linked zRGα1a dataset on the Talos Arctica microscope at the Francis Crick Institute.

## Data availability

The coordinates for the zRET-CLD(1-4), zGDNF-zGFRα1a and zRGα1a are available in the PDB with the primary accession code XXXX, 7AB8 and XXXX, respectively. The zRGα1a C2 symmetry applied map, the zRGα1a symmetry expanded map and the hR15AL negative stain envelopes are available on the EMDB with accession codes XXXX, XXXX and XXXX, respectively.

## Author information

The authors declare no competing financial interests. Correspondence and requests for materials should be addressed to N.Q.M.

## Notes

**Competing interests:** The authors declare that no competing interests exist

### Competing Interest Statement

The authors have declared no competing interest.

