## Supplementary Information for "A two-site flexible clamp mechanism for RET-GDNF-GFRα1 assembly reveals both conformational adaptation and strict geometric spacing"

**Supplementary Methods**

*XL-MS analysis of the zRET$^{ECD}$-GFRα1-GDNF complex:*

All chemicals were purchased from Sigma at the highest purity unless otherwise stated. A total of 200 ng protein in 20 mM HEPES (pH 7.5), 150 mM NaCl, 1 mM $CaCl_2$ was cross-linked using 1 mM disuccinimidyl sulfoxide (DSSO)(Kao et al., 2011) (Thermo Fisher Scientific) with mild shaking for 15 min at 37 °C. The reaction was quenched using a final concentration of 5% hydroxylamine for a further 15 min at 37 °C. The sample was subsequently alkylated, reduced and proteolysed. To do this, the sample was dried to completion using vacuum centrifugation and resolubilised with sonication into 8 M urea. Cysteine reduction was carried out using 2.5 mM TCEP for 30 min at 37 °C and alkylated in the dark using 5 mM iodoacetamide at room temperature for 30 min. The urea was diluted to 1 M using 50 mM triethylammonium bicarbonate and proteins were proteolysed using trypsin (Pierce) at 1:50 w/w trypsin:protein overnight at 37 °C. The solution was acidified to pH 2-3 using trifluoroacetic acid (TFA) and desalted using in house built STAGE tips made using Empore SPE C18 disks (3M, 66883-U). The eluent was then dried to completion.

*Liquid Chromatography Mass Spectrometry*

Peptides were reconstituted in 0.1 % TFA (v/v) and chromatographically resolved using an Ultimate 3000 RSLCnano (Dionex) HPLC. Peptides were first loaded onto an Acclaim PepMap 100 C18, 3 µm particle size, 100 Å pore size, 20 mm x 75 µm ID (Thermo Scientific, 164535) trap column using a loading buffer (2 % acetonitrile (MeCN) (v/v) and 0.05 % TFA in 97.95 % H2O) with a flow rate of 7 µL/min. Chromatographic separation was achieved using an EASY-Spray column, PepMap C18, 2 µm particles, 100 Å pore size, 500 mm x 75 µm ID (Thermo Scientific, ES803). The gradient utilised a flow of 0.275 µL/min, starting at 98 % mobile A (0.1% formic acid, 5 % dimethyl sulfoxide (DMSO) in $H_2O$) and 2 % mobile B (0.1 % formic acid, 75 % MeCN, 5% DMSO and 19.9 % H2O). After 3 min mobile B was increased to 8 % over 3 min, increased to 25 % over 69 min, to 45 % over 35 min, further increased to 90% in 17 min and held for 5 min. Finally, mobile B was reduced back to 5 % over 3 min for the rest of the acquisition.

MS1 data were acquired in real time over 150 minutes using an Orbitrap Fusion Lumos Tribrid mass spectrometer in positive, top speed mode with a cycle time of 5 s. The chromatogram (MS1) was captured using 60,000 resolution, a scan range of 375-1500 with a 50 ms maximum injection time, and 4e5 AGC target. MS2 dynamic exclusion with repeat count 2, exclusion duration of 30 s, 20 ppm tolerance window was used, along with isotope exclusion, a minimum intensity exclusion of 2e4, charge state inclusion of 3-8 ions and peptide mono isotopic precursor selection. Precursors within a 1.2 *m/z* isolation window were then fragmented using 25 % normalised collusion-induced dissociation (CID), 100 ms maximum injection time and 5 e4 AGC target. Scans were recorded using 30,000 resolution in centroid mode starting 120 *m/z*. MS3 spectra containing peaks with a mass difference of 31.9721 Da were further fragmented with a 43 % normalised higher collision induced dissociation, using a 2 *m/z* isolation window, 150 ms maximum injection time and 2e4 AGC target. 4 scans were recorded using an ion trap detection in rapid mode starting at 120 *m/z*.

*Data analysis.*

Data processing was carried out using Proteome Discoverer Version 2.3 (ThermoFisher Scientific) with the XlinkX node(2017; Liu et al., 2015). The acquisition strategy was set to MS2_MS3 mode. The database comprised solely of the specific $zRET^{ECD}$, $zGFR\alpha1a^{D1\text{-}3}$ and $GDNF^{mat.}$ sequences. Trypsin was selected as the proteolytic enzyme allowing up to two missed cleavages with a minimal peptide length of five residues. Masses considered were in the range of 0.3-10 kDa. The precursor mass tolerance, FTMS fragment mass tolerance, and ITMS Fragment Mass Tolerance were set to 10 ppm, 20 ppm and 0.5 Da respectively. A static carbamidomethyl (+57.021 Da) modification was utilised for cysteine residues, with an additional dynamic modification for oxidation (+15.995 Da) on methionine residues. The False Discovery Rate (FDR) threshold was set to 0.05 with percolator as the strategy. The list of reported cross-linked spectral matches were manually examined and cross-links with spectra that did not contain acceptable b and y ion coverage were excluded. The reduced list was exported to crosslinkviewer.org(Combe et al., 2015) in order to graphically view the cross-links.

| | zCLD(1-4)$^{red.sug.}$ | zGDNF$^{mat.}$-GFRα1a$^{ΔD1}$ |
|---|---|---|
| Wavelength (Å) | 0.9787 | 0.9795 |
| Resolution range * | 65.96 - 2.20 (2.28 - 2.20) | 50.76 - 2.2 (2.28 - 2.2) |
| Space group | P 1 | P 21 21 2 |
| Unit cell dimensions | | |
| *a*, *b*, *c* (Å) | 51.17 70.50 105.44 | 125.07 55.54 70.96 |
| α, β, γ (°) | 105 101 100 | 90 90 90 |
| Total reflections | 229073 (22789) | 51646 (5054) |
| Unique reflections | 67550 (4539) | 25823 (1368) |
| Multiplicity | 3.4 (3.4) | 2.0 (2.0) |
| Completeness (%) | 91.28 (66.36) | 91.93 (53.94) |
| Mean I/sigma(I) | 7.09 (1.92) | 14.30 (3.15) |
| Wilson B-factor (Å$^2$) | 28.88 | 20.87 |
| R-merge | 0.073 (0.68) | 0.056 (0.37) |
| R-meas | 0.087 (0.81) | 0.079 (0.53) |
| R-pim | 0.046 (0.43) | 0.056 (0.37) |
| CC1/2 | 0.996 (0.70) | 0.997 (0.80) |
| CC | 0.999 (0.91) | 0.999 (0.94) |
| Resolution used for refinement | 65.96 - 2.20 | 50.76-2.20 |
| Reflections used in refinement | 62771 (4522) | 23743 (1363) |
| Reflections used for R-free | 3098 (255) | 1152 (57) |
| R-work | 0.232 (0.316) | 0.199 (0.247) |
| R-free | 0.277 (0.383) | 0.230 (0.248) |
| CC (work) | 0.895 (0.615) | 0.888 (0.723) |
| CC (free) | 0.884 (0.553) | 0.888 (0.800) |
| Number of non-hydrogen atoms | 7997 | 2736 |
| macromolecules | 7289 | 2434 |
| ligands | 539 | 83 |
| solvent | 169 | 219 |
| Protein residues | 980 | 309 |
| RMS (bonds Å) | 0.009 | 0.006 |
| RMS (angles °) | 1.08 | 0.74 |
| Ramachandran favoured (%) | 96.6 | 97.03 |
| Ramachandran allowed (%) | 3.0 | 2.97 |
| Ramachandran outliers (%) | 0 | 0.0 |
| Rotamer outliers (%) | 0 | 0.0 |
| Clashscore | 14.56 | 4.03 |
| Average B-factor (Å$^2$) | 41.31 | 30.19 |
| macromolecules | 39.53 | 28.99 |
| ligands | 67.62 | 55.09 |
| solvent | 33.82 | 34.06 |
| Number of TLS groups | 8 | 1 |
| PDB code | XXXX | 7AB8 |

*Values in parentheses are for highest-resolution shell.

**Supplementary Table 1: Crystallography statistics**

| | zRGα1a C2 map | zRGα1a Symmetry Expanded Map | zR15AL Negative Stain Map |
|---|---|---|---|
| EMDB ID | XXXX | XXXX | XXXX |
| PDB ID | XXXX | | |
| Magnification | 46,296 | 46,296 | 40,719 |
| Voltage (kV) | 300 | 300 | 120 |
| Electron exposure ($e^-$/Å$^2$) | 48.6 | 48.6 | |
| Defocus Range (μm) | 1.4-3.5 | 1.4-3.5 | 1.0-1.5 |
| Pixel Size (Å) | 1.08 | 1.08 | 3.44 |
| Symmetry imposed | C2 | C1 | C2 |
| Initial particle images | 2,424,600 (dataset 1), 1,393,023 (dataset 2) | | 27,551 |
| Final particle images | 382,547 (360,189 dataset 1 and 22,358 dataset 2) | 382,547 (360,189 dataset 1 and 22,358 dataset 2) | 6,519 |
| Map resolution (Å) | 3.3 | 3.5 | 26 |
| FSC threshold | 0.143 | 0.143 | 0.143 |
| Map resolution range (Å) | 12 - 3.3 | 11 - 3.5 | |
| *Refinement* | | | |
| Initial model used (PDB ID) | XXXX, 7AB8 | | |
| Model resolution (Å) | 4.2 | | |
| FSC threshold | 0.5 | | |
| Model resolution range (Å) | | | |
| Map sharpening *B* factor (Å$^2$) | -75 | | |
| Model composition | | | |
| Non-hydrogen atoms | 16020 | | |
| Protein residues | 1996 | | |
| Ligands | 8 | | |
| N-glycans | 16 | | |
| *B*-factors (Å$^2$) | | | |
| Protein | 122.4 | | |
| Ligands | 111.6 | | |
| R.m.s. deviations | | | |
| Bond lengths (Å) | 0.004 (0) | | |
| Bond angles (°) | 0.646 (6) | | |
| Validation | | | |
| MolProbity score | 1.85 | | |
| Clashscore | 9.45 | | |
| Poor rotamer (%) | 0.89 | | |
| Ramachandran plot | | | |
| Favoured (%) | 94.94 | | |
| Allowed (%) | 5.06 | | |
| Disallowed (%) | 0.0 | | |

**Supplementary Table 2: EM data acquisition and processing statistics**

| | **LIGAND** | | | | **CO-RECEPTOR** | | | | **RET** | |
|---|---|---|---|---|---|---|---|---|---|---|
| | zGDNF | hGDNF | hNRTN | hGDF15 | zGFRα1 | hGFRα1 | hGFRα2 | hGFRAL | zRET | hRET |
| **LIGAND** | | | | | | | | | | |
| zGDNF | 1530Å | | | | 866Å | | | | 347Å | |
| zGDNF* | | | | | | | | | 251Å | |
| hGDNF | | 1003Å | | | | 961Å | | | | 187Å |
| hGDNF* | | | | | | | | | | 148Å |
| hNRTN | | | 2006Å | | | | 843Å | | | 296Å |
| hNRTN* | | | | | | | | | | 218Å |
| hGDF15 | | | | 1398Å | | | | 577Å | | 407Å |
| hGDF15* | | | | | | | | | | 369Å |
| **CO-RECEPTOR** | | | | | | | | | | |
| zGFRa1 | | | | | | | | | 846Å | |
| hGFRa1 | | | | | | | | | | 872Å |
| hGFRa2 | | | | | | | | | | 960Å |
| hGFRAL | | | | | | | | | | 1094Å |

Interface sizes (averaged over both protomers) calculated by PDBePISA

* Contact surface to the second protomer of GFL dimer

**Supplementary Table 3: Major interface sizes for ternary complexes of hRET/zRET, GFRa1/a2/GFRAL and GDNF/NRTN/GDF15**

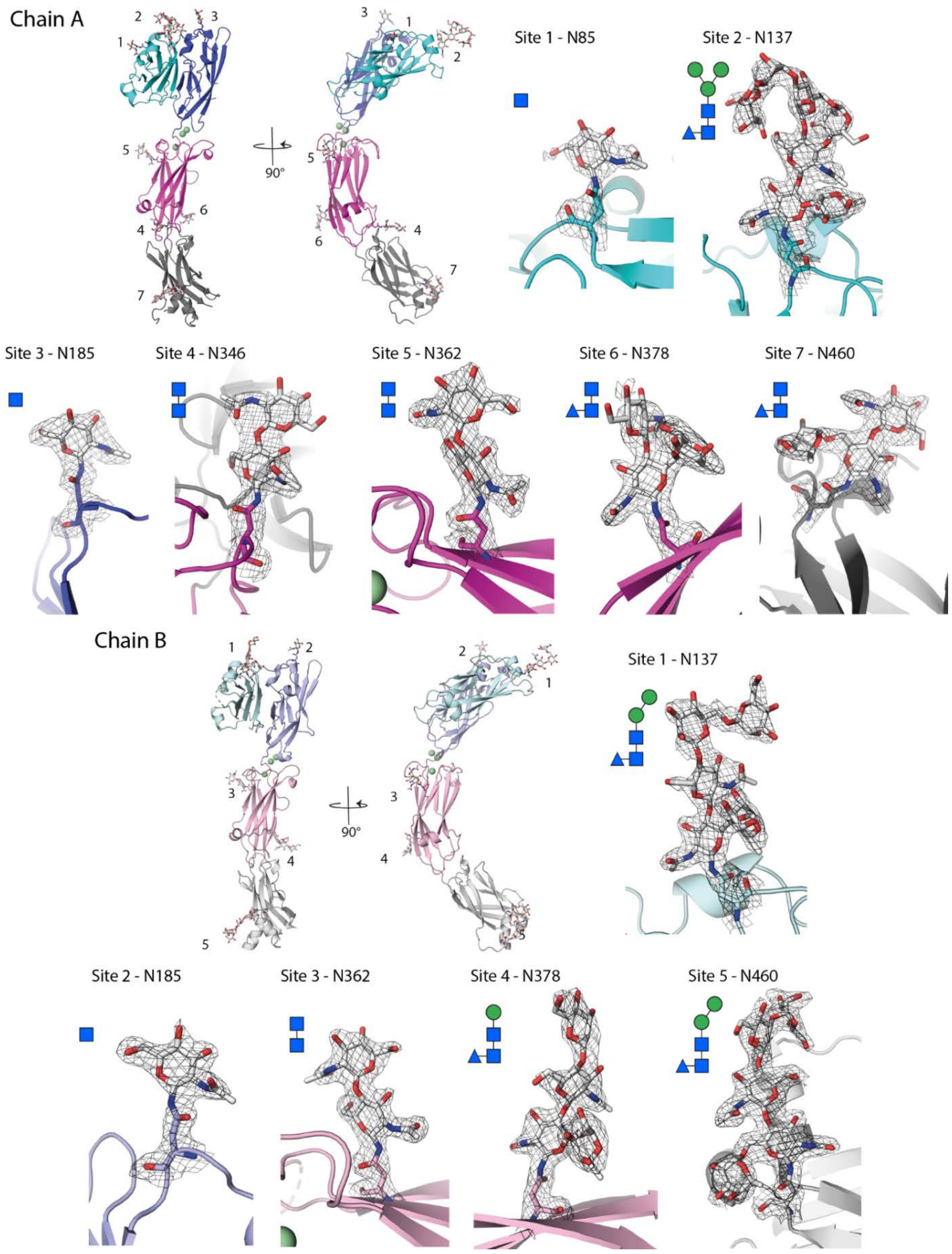

**Supplementary Figure 1: An abundance of N-linked glycans in the zRET$^{CLD1-4}$ module.** Location of N-linked sites within the zRET$^{CLD1-4}$ module and final electron density contoured at 1.0 σ corresponding to each N-linked glycosylation site are shown. Each site is numbered with the overall structure represented in cartoon and coloured according to each domain; CLD1 [cyan (chain A)/pale cyan (chain B)], CLD2 [blue (chain A)/pale blue (chain B)], CLD3 [magenta (chain A)/pink (chain B)], and CLD4 [grey (chain A)/light grey (chain B)]. The refined glycans are drawn in symbol form according to accepted convention, with *N*-acetylglucosamine, α-fucose and mannose represented as a blue square, blue triangle and a green circle, respectively. All images were rendered in PyMOL (Schrodinger, 2015).

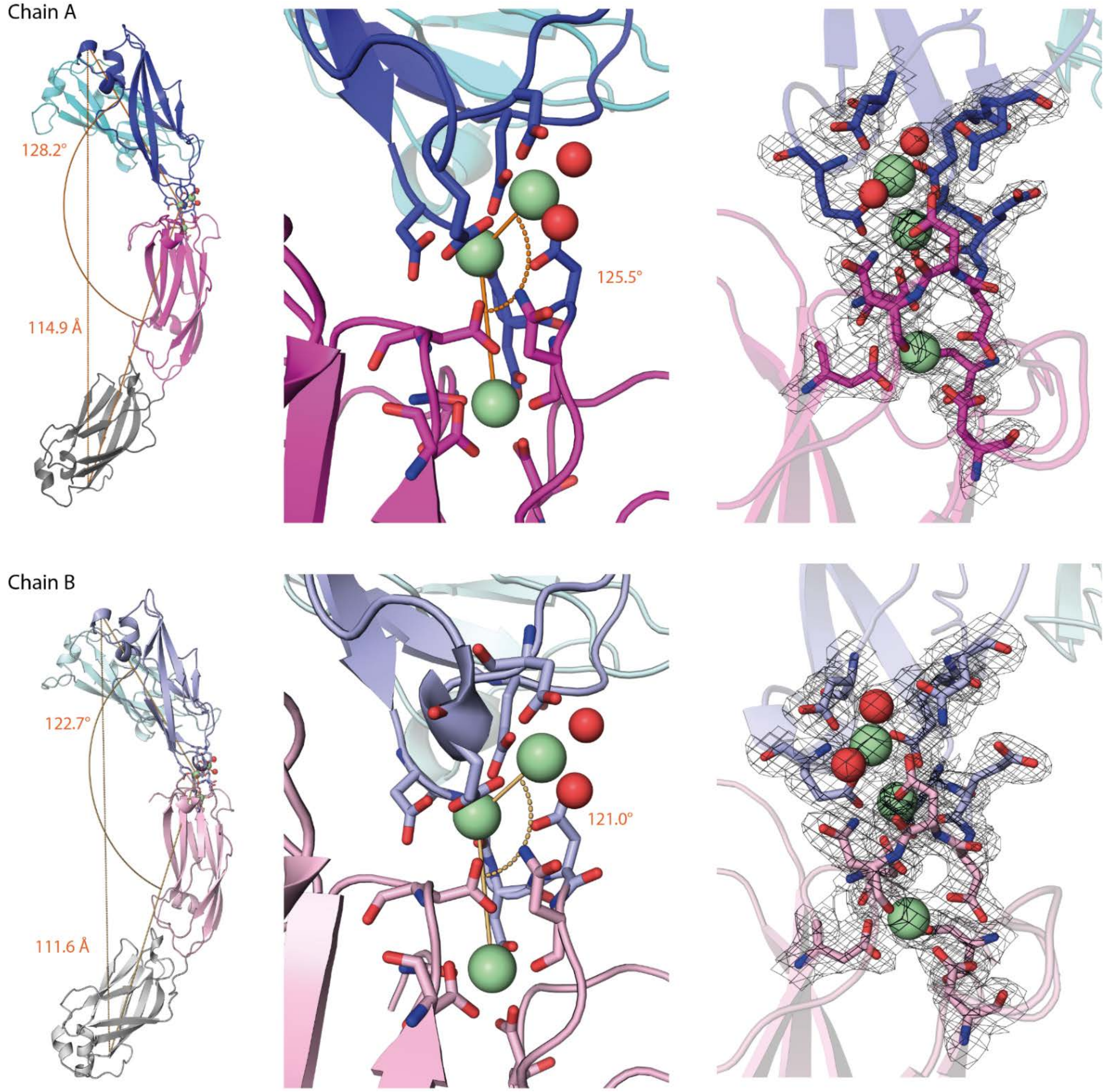

**Supplementary Figure 2: Flexibility at the calcium site of zRET$^{CLD1-4}$ propagates along the module.** Two molecules of zRET$^{CLD1-4}$ in the asymmetric unit. The overall structure of chain A and chain B is represented as a cartoon, with the CLD1 in cyan, CLD2 in blue, CLD3 in magenta and CLD4 in grey for chain A and in chain B; CLD1 in pale cyan, CLD2 in pale blue, CLD3 in light pink and CLD4 in light grey. Close up views of the calcium-binding site superposed onto the final electron density map calculated using m2Fo-DFc coefficients and contoured at 1.0 σ. The differences in the calcium ion positions in chain A and B give larger differences over the module. All images rendered in PyMOL (Schrodinger, 2015).

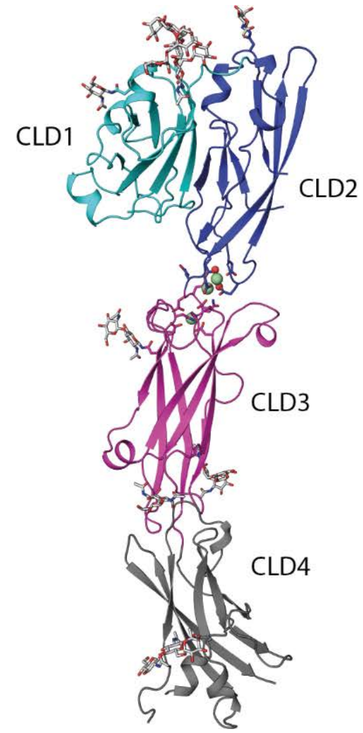

| | E-Cadherin D1 | E-Cadherin D2 | N-Cadherin D1 |
|---|---|---|---|
| CLD1 | 1.635 Å (15.19%) | 1.893 Å (14.89%) | 1.557 Å (13.92%) |
| CLD2 | 1.743 Å (23.17%) | 1.900 Å (18.29%) | 1.997 Å (23.81%) |
| CLD3 | 5.116 Å (10.00%) | 4.851 Å (15.38%) | 5.119 Å (6.12%) |
| CLD4 | 1.933 Å (11.69%) | 1.839 Å (18.39%) | 2.105 Å (9.41%) |

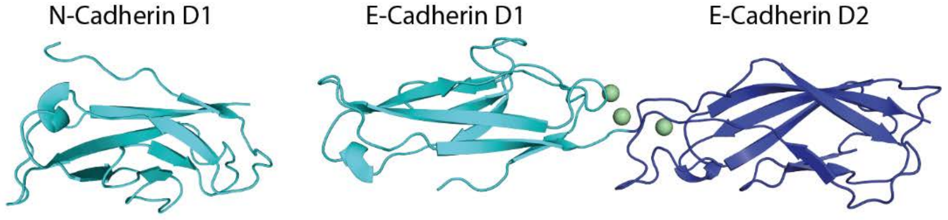

**Supplementary Figure 3: Structural comparisons of individual zRET cadherin-like domains and classical cadherins.** A table showing the calculated RMSD values (and sequence identity) between each of the cadherin like domains from the structure of zCLD1-4$^{red.sug.}$ and the structures of the classical cadherins; E-cadherin domains 1 and 2 (PDB 1EDH)(Nagar et al., 1996) a N-cadherin domain 1 (PDB 1NCI)(Shapiro et al., 1995). The structures of zCLD1-4 and N-cadherin D1 and E-cadherin D1 and D2 are represented as cartoon with zCLD1 in cyan, zCLD2 in blue, zCLD3 in magenta, zCLD4 in grey, N-cadherin D1 in cyan, E-cadherin D1 in cyan and E-cadherin D2 in blue. Chain A of each structure was used to calculate the RMSD and sequence identity in COOT(Emsley and Cowtan, 2004) using SSM superpose function. All the images were rendered in PyMOL(Schrodinger, 2015).

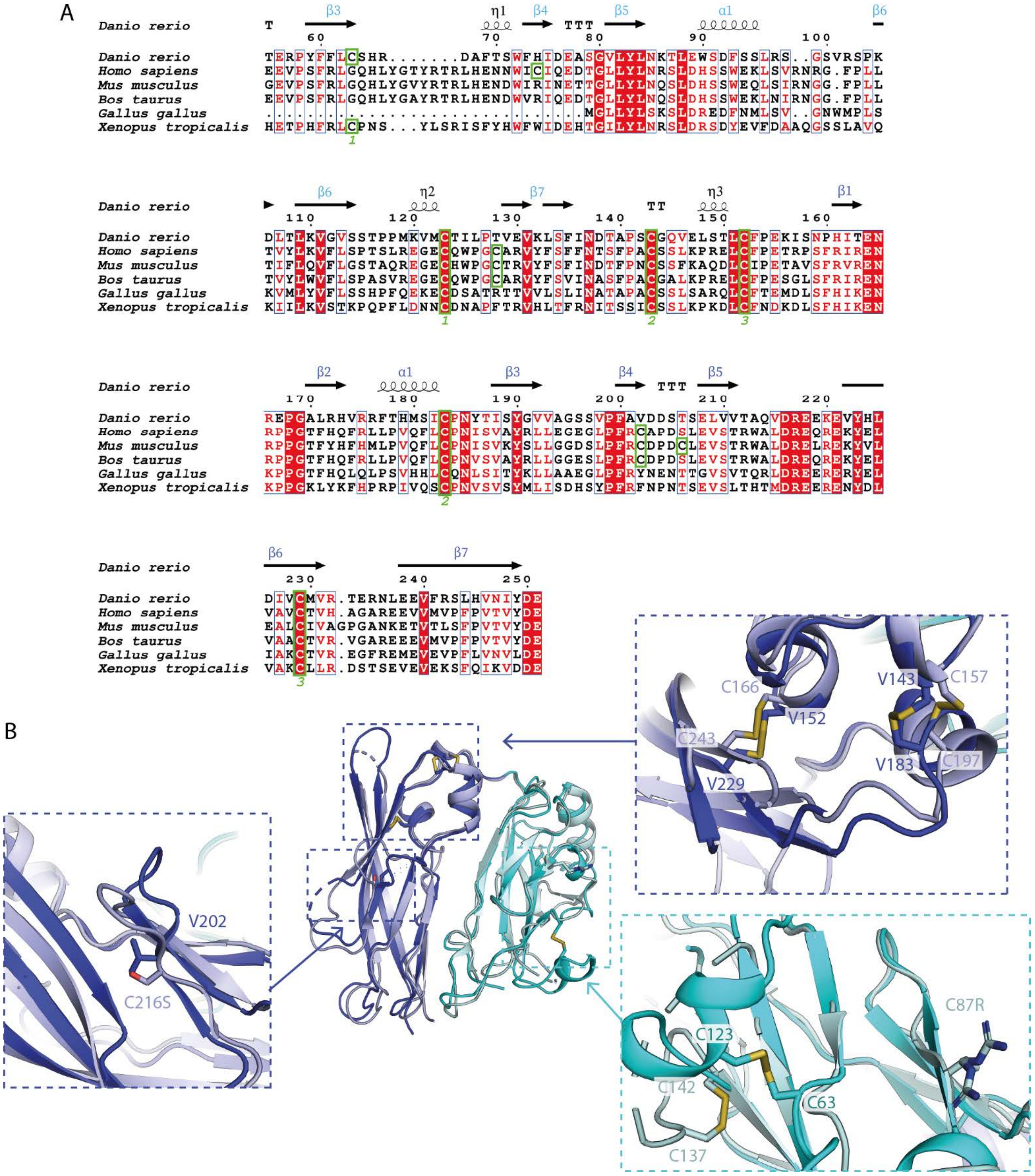

**Supplementary Figure 4: Structural divergence within zRET$^{CLD1\text{-}2}$ is driven by a disulfide swap between higher and lower vertebrates.** A) Sequence alignment of the C-terminal portion of CLD1 and CLD2 in higher and lower vertebrates; *Danio rerio* (Uniprot P04749)*, Homo sapiens* (Uniprot P04749)*, Mus musculus* (Uniprot P04749), *Bos taurus* (Uniprot P04749), *Gallus gallus* (Uniprot P04749), and *Xenopus tropicalus* (Uniprot P04749). Cysteines within the sequence are highlighted, demonstrating the disulfide bond shuffling. B) The structures of zCLD1-2 (in cyan and blue) aligned with the hCLD12 structure (in pale cyan and pale blue, PDB 2X2U), represented as a cartoon, with close-ups of each of the disulfides located within the structure, highlighted as sticks. The two unpaired cysteines unique to hRET were mutated to and arginine (C87R) and a serine (C216S) to aid structure determination of hRET$^{CLD1\text{-}2}$ (Kjær et al., 2010). Images were rendered in PyMOL (Schrodinger, 2015).

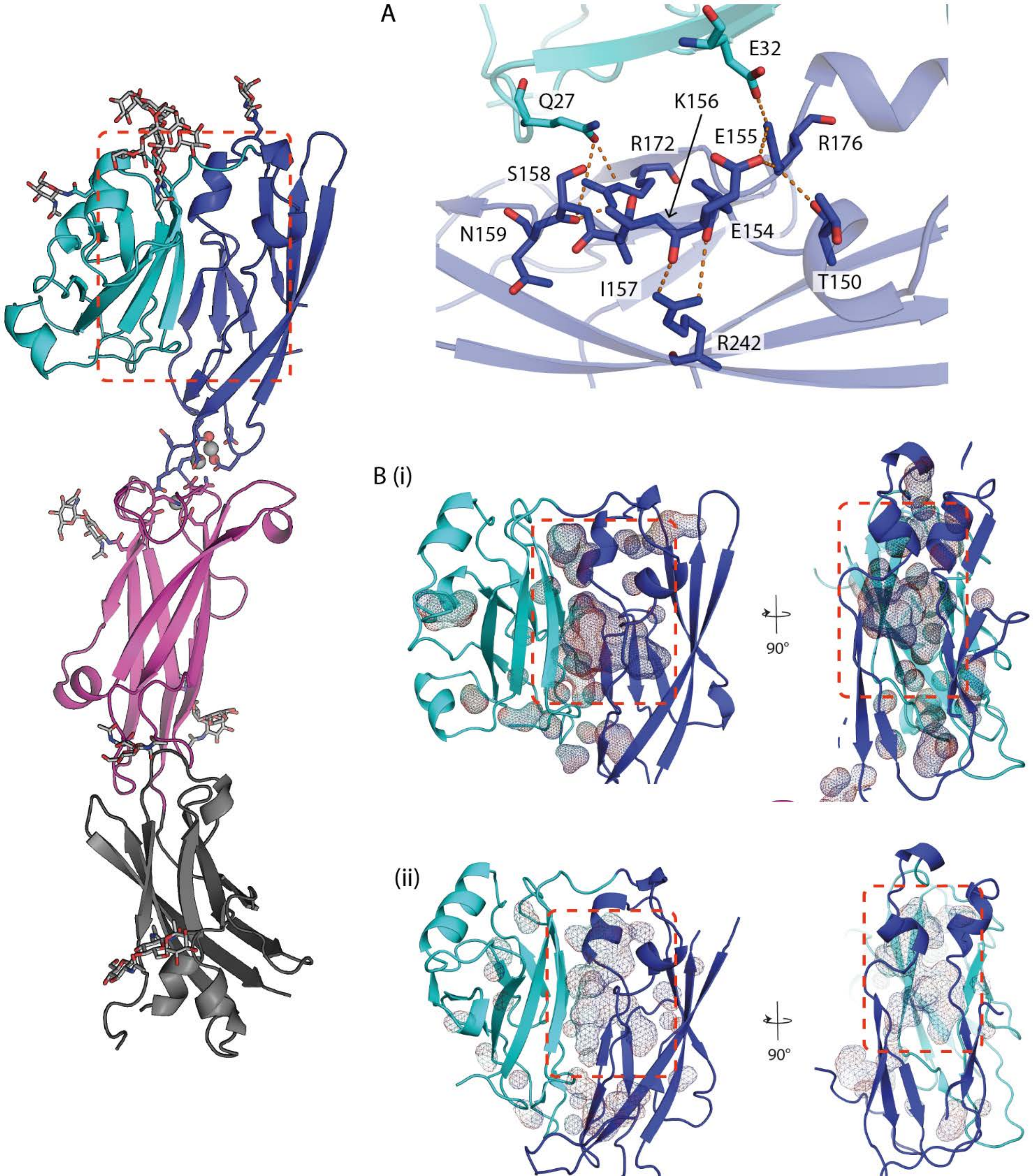

**Supplementary Figure 5: zRET$^{CLD1-2}$ clamshell interface and an internal cavity adjacent to CLD2-ß1 strand.** The overall structure of zCLD1-4 represented as a cartoon and CLD1 in cyan, CLD2 in blue, CLD3 in magenta and CLD4 in grey. A) The residues incorporated into the CLD1-2 clamshell interface; with CLD2-β1 stabilised with R242 (CLD2-β6), R172 and R176 (CLD2-β2) the latter of which also interact with Q27 and E32 (CLD1-β1), residues are highlighted as sticks. B Orthogonal views of the cavity (shown in mesh) within CLD1-2 for both zebrafish (i) and human (ii) (PDB 2X2U)(Kjær et al., 2010). Rendered in PyMOL (Schrodinger, 2015).

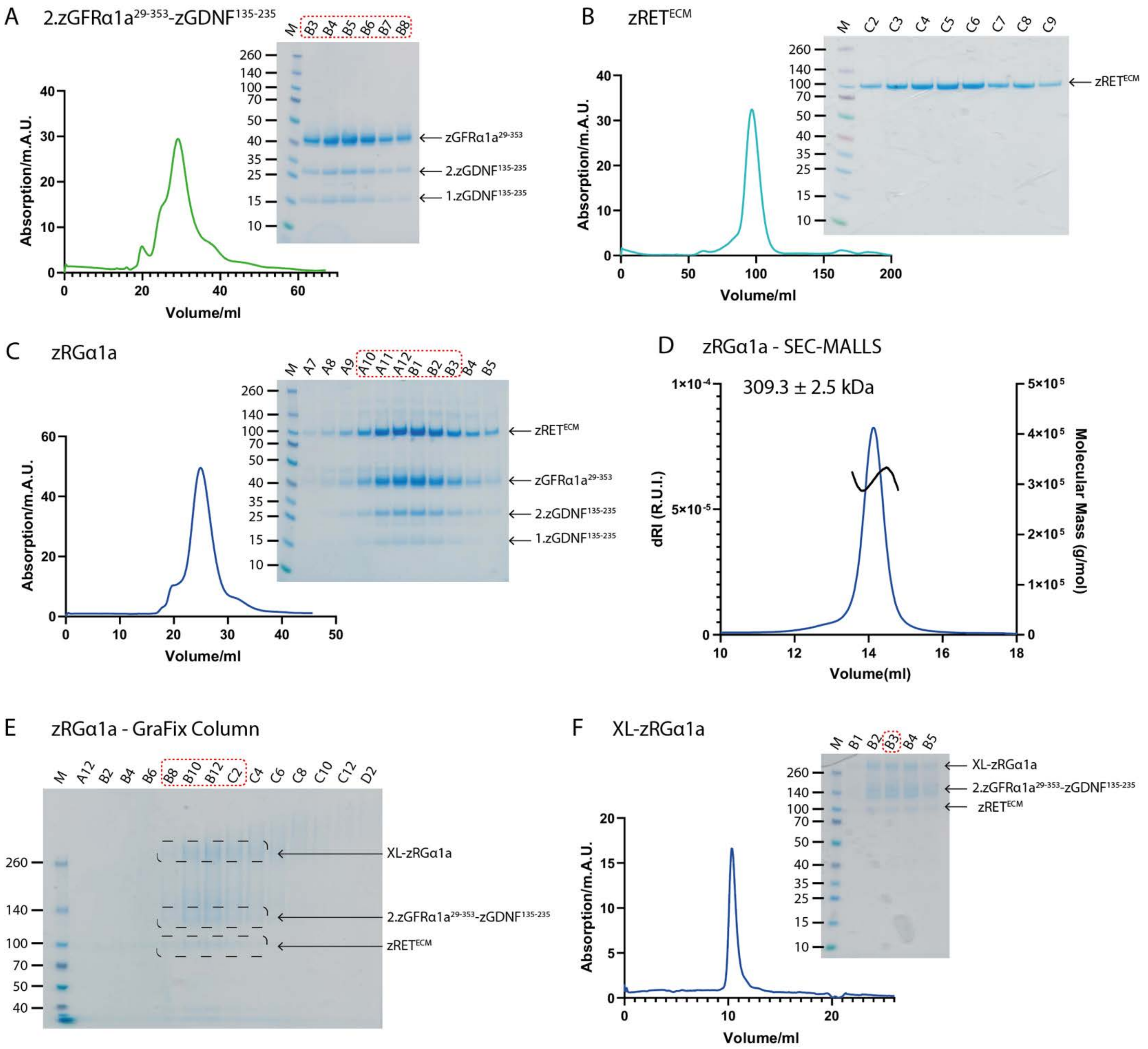

**Supplementary Figure 6: Purification of individual zRGα1a components and crosslinking of the zRGα1a ternary complex.** Size exclusion profiles of A) 2.zGFRα1a$^{29\text{-}353}$-zGDNF$^{135\text{-}235}$, B) zRET$^{ECD}$ and C) zRGα1a .D) SEC-MALLS trace of the purified zRGα1a. E) SDS-PAGE of fractionated GraFix stabilised zRGα1a sample. F) Size exclusion profile of the crosslinked zRGα1a sample.

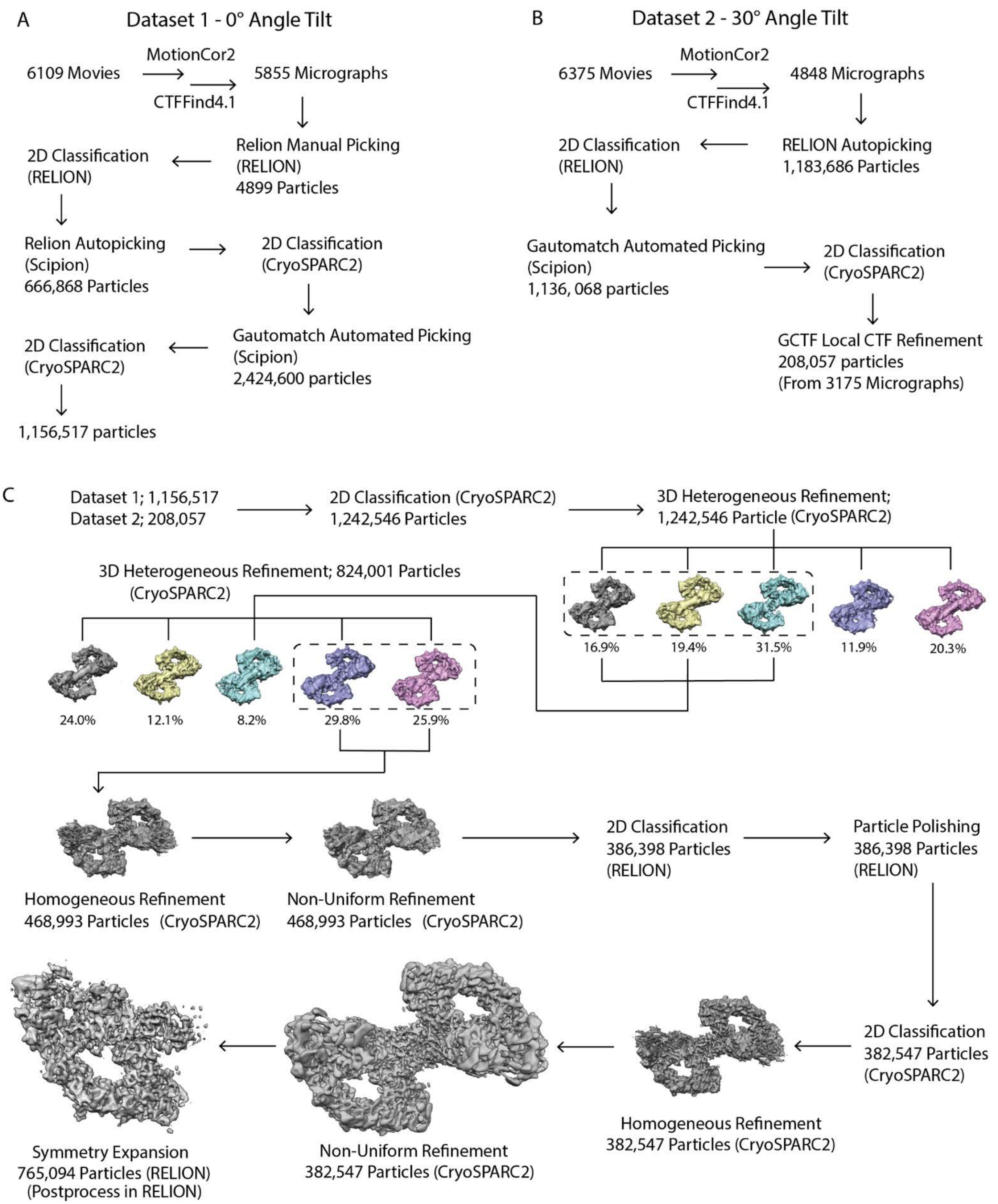

**Supplementary Figure 7: zRGα1a cryo-EM data processing workflow.** A) Non-tilted particles dataset 1 and B) tilted particles dataset 2 were processed independently. C) Combined particles from both datasets 1 and 2 and workflow. Software packages used; CryoSPARC2(Punjani et al., 2017), CTFFind4.1(Rohou and Grigorieff, 2015), Gautomatch [K. Zhang, MRC LMB (www.mrc-lmb.cam.ac.uk/kzhang/)], GCTF(Zhang, 2016), MotionCor2(Zheng et al., 2017) RELION(Kimanius et al., 2016; Scheres, 2012; Zivanov et al., 2018), Scipion(de la Rosa-Trevín et al., 2016),

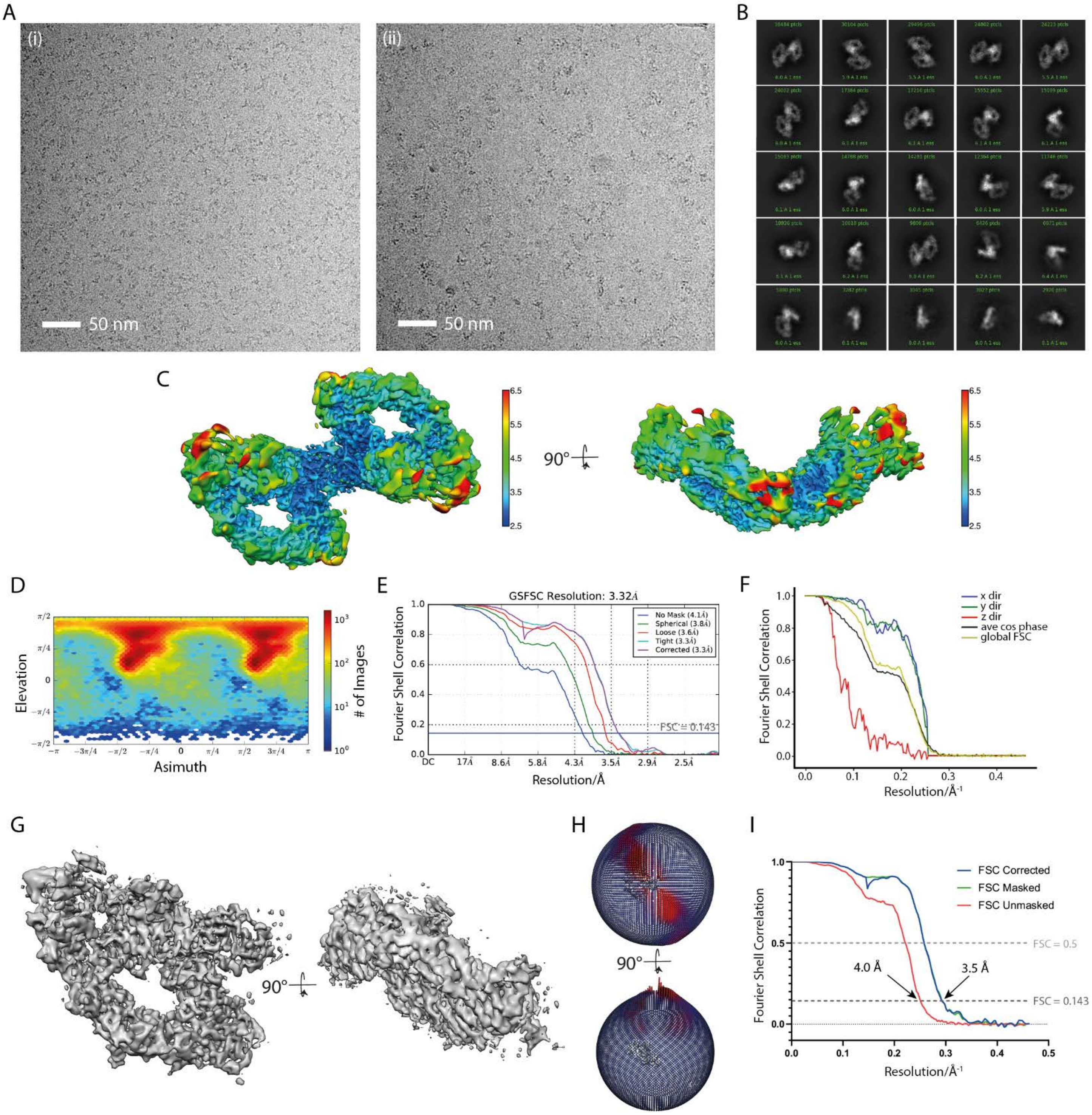

**Supplementary Figure 8: zRGα1a single particle cryo-EM reconstruction.** A) Example micrographs from the non-tilted dataset (i) and the dataset collected with a tilt angle of 30° (ii). B) Twenty-five 2D class averages from all the particles in the final reconstruction. C) The C2 averaged cryo-EM map is coloured by local resolution, generated by locRes option in CryoSPARC2 (Punjani et al., 2017) using blue for 2.5 Å resolution areas, green for 4.5 Å and red for 6.5 Å resolution. A total of 382,574 particles were generated using CryoSPARC2 (Punjani et al., 2017) non-uniform refinement and postprocessed in RELION(Kimanius et al., 2016; Scheres, 2012; Zivanov et al., 2018). Two orthogonal views of the zRGα1a complex are shown. D) Angular distribution of the particles in the C2 averaged map. E) Fourier shell correlation curve of the C2 averaged map. F) A 3DFSC(Tan et al., 2017) plot generated from the C2 averaged map. G) Two views orthogonal of the symmetry expanded map, generated using particles expansion in RELION (Kimanius et al., 2016; Scheres, 2012; Zivanov et al., 2018). H) Two orthogonal views of the angular distribution from the symmetry expanded map. I) The Fourier shell correlation curves from the symmetry expanded map, which shows an overall resolution of 3.5 Å. Images of the maps were rendered using Chimera (Pettersen et al., 2004).

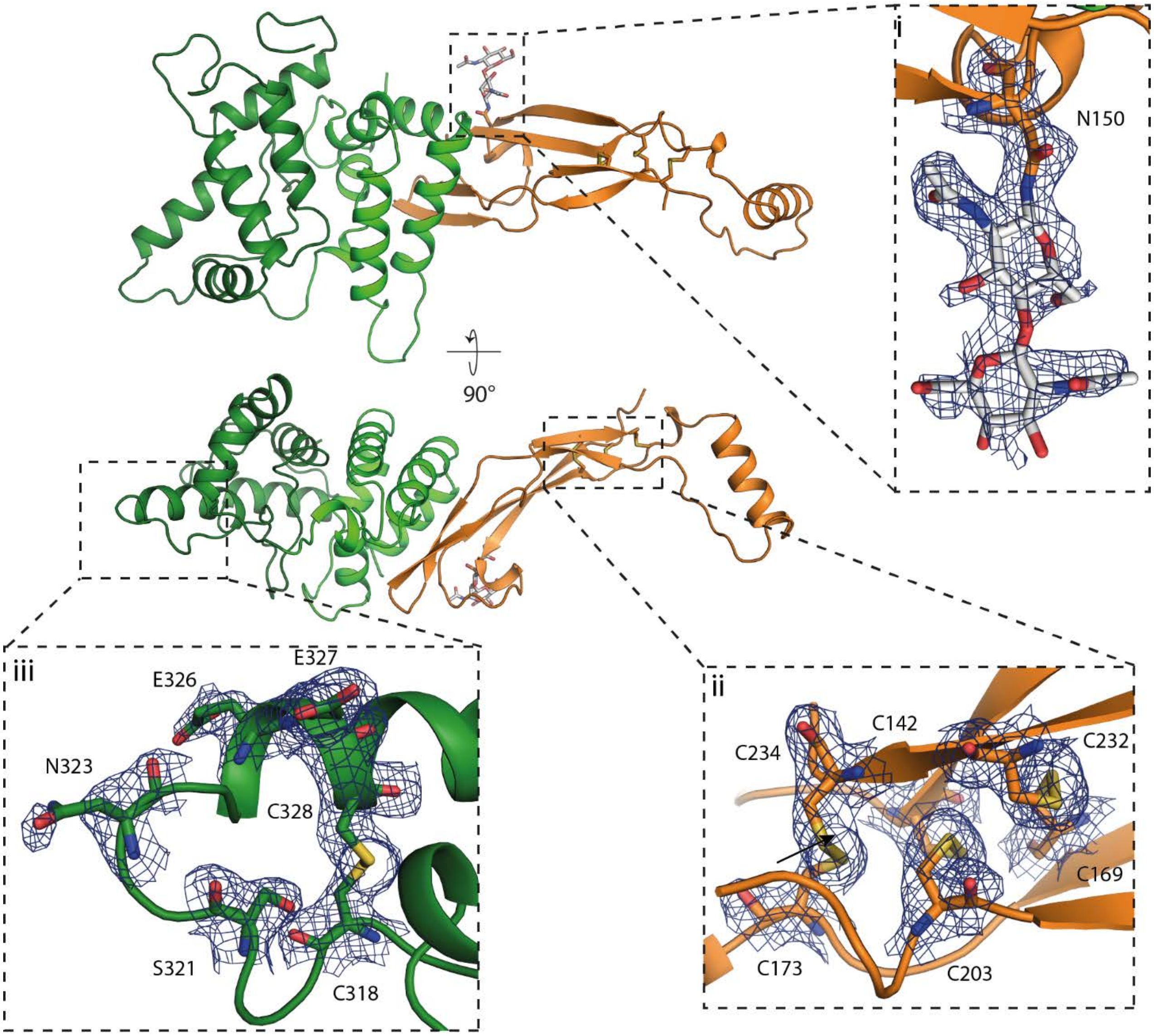

**Supplementary Figure 9: Structure determination and map quality for zGDNF$^{138\text{-}235}$zGFRα1a$^{151\text{-}353}$.** Final 2.2 Å structure of the contents of the asymmetric unit with a single copy of zGDNF$^{138\text{-}235}$zGFRα1a$^{151\text{-}353}$. The insets reveal the final electron density calculated for different areas of the structure using m2Fo-DFc coefficients and contoured at 1.0 σ. The glycosylation site located on N150 of GDNF (i), the binding site of zGFRα1a that interacts with RET CLD(2-3) calcium site (ii), and the disulfide bond network in zGDNF that makes it part of the cystine knot family (iii). All images were rendered in PyMOL (Schrodinger, 2015).

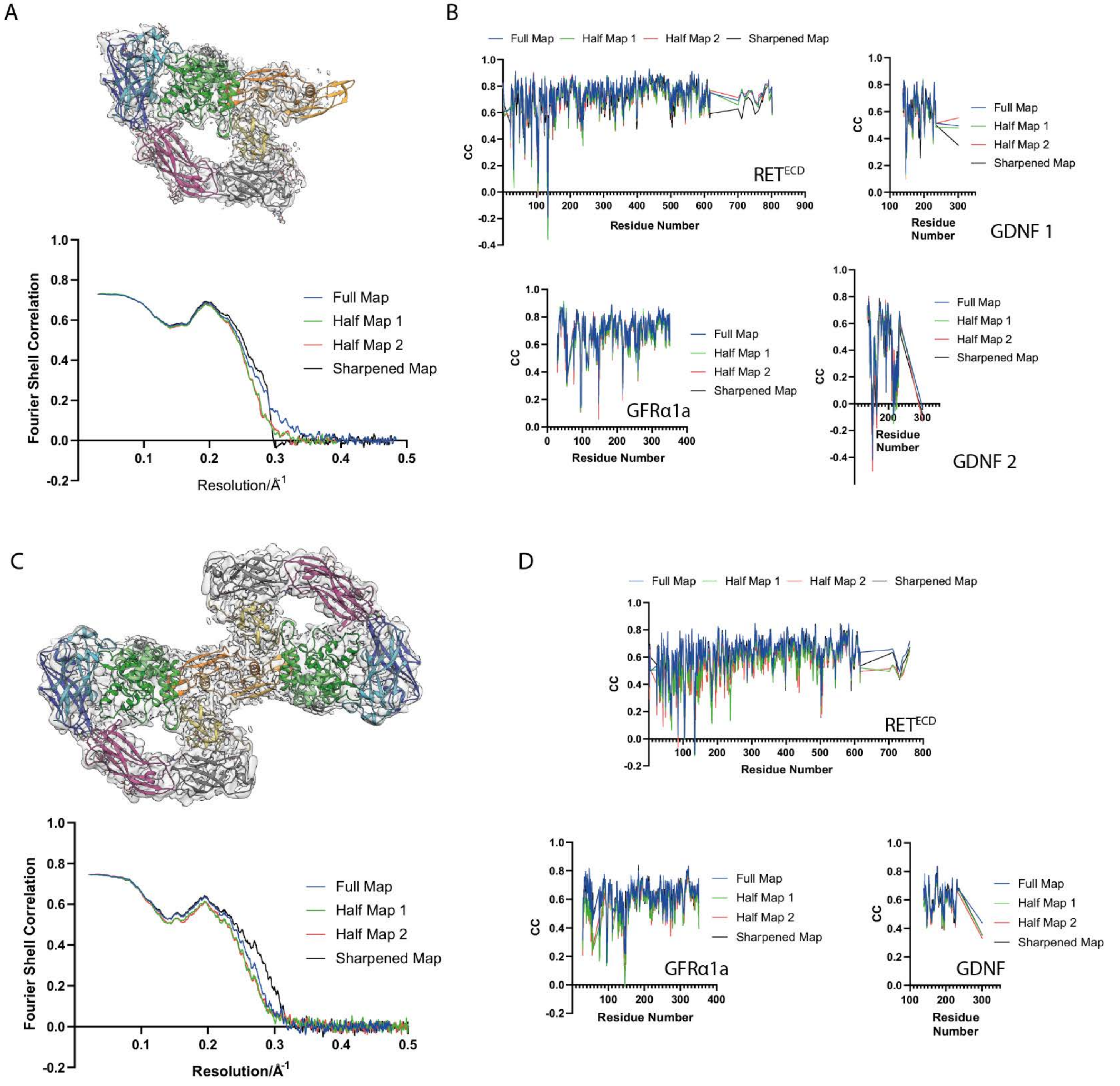

**Supplementary Figure 10: Correlation and validation of the symmetry expanded and C2 averaged zRGα1a reconstruction.** A) Map-to-model Fourier shell correlation curve of the symmetry expanded map and the zRGα1a structure. B) Cross correlation between each residue in the model and the symmetry expanded map. C) Map-to-model Fourier shell correlation between the C2 averaged cryo-EM map and the zRGα1a model. D) Cross correlation between each residue in the model and the C2 averaged map. All correlation statistics were provided with the use of the full maps and two half maps in each case using Phenix cryo-EM model validation tools(Afonine et al., 2018).

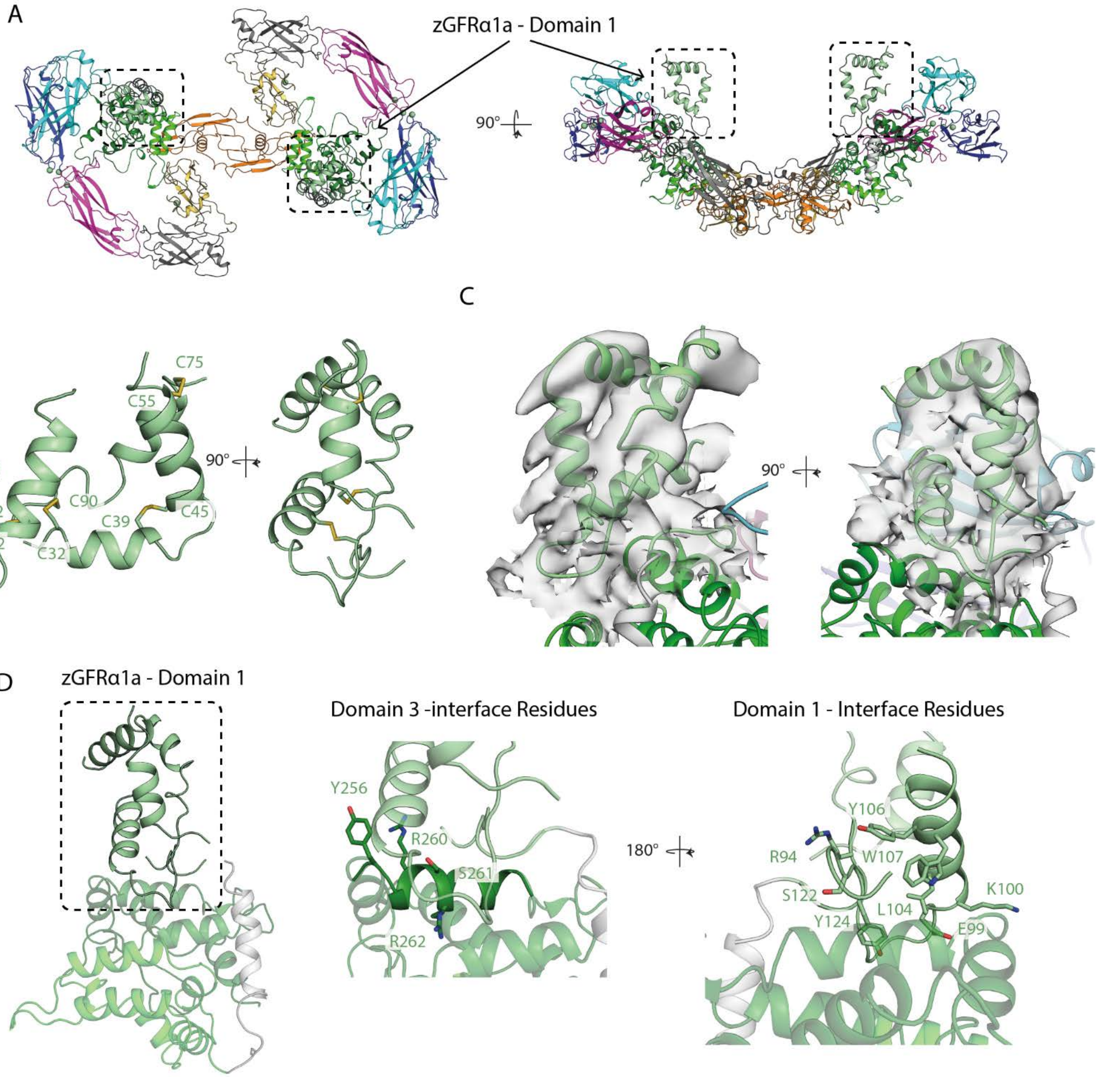

**Supplementary Figure 11: Fitting of the GFRα1a$^{D1}$ into the final zRGα1a structure** A) Two views of the C2 averaged model with zGFRα1a Domain 1 (zGFRα1a$^{D1}$) positioning highlighted. The final structure is a cartoon with the domains within the structure individually coloured; CLD1 in cyan, CLD2 in blue, CLD3 in magenta, CLD4 in grey, CRD in yellow, zGFRα1a$^{D1}$ in pale green, zGFRα1a$^{D2}$ in green, zGFRα1a$^{D3}$ in dark green, and zGDNF in orange. B) Helical structure of zGFRα1a$^{D1}$ with the disulfide interconnectivity to stabilise the structure. C) zGFRα1a$^{D1}$ with the segmented symmetry expanded map surrounding the model, with an inset showing the map and model close to disulfide bond between C39-C45. Images rendered in Chimera(Pettersen et al., 2004). D) A cartoon representation of zGFRα1a subunit with zGFRα1a$^{D1}$ domain are highlighted, two insets show the interface. Images in parts A, B and C were rendered in PyMOL (Schrodinger, 2015).

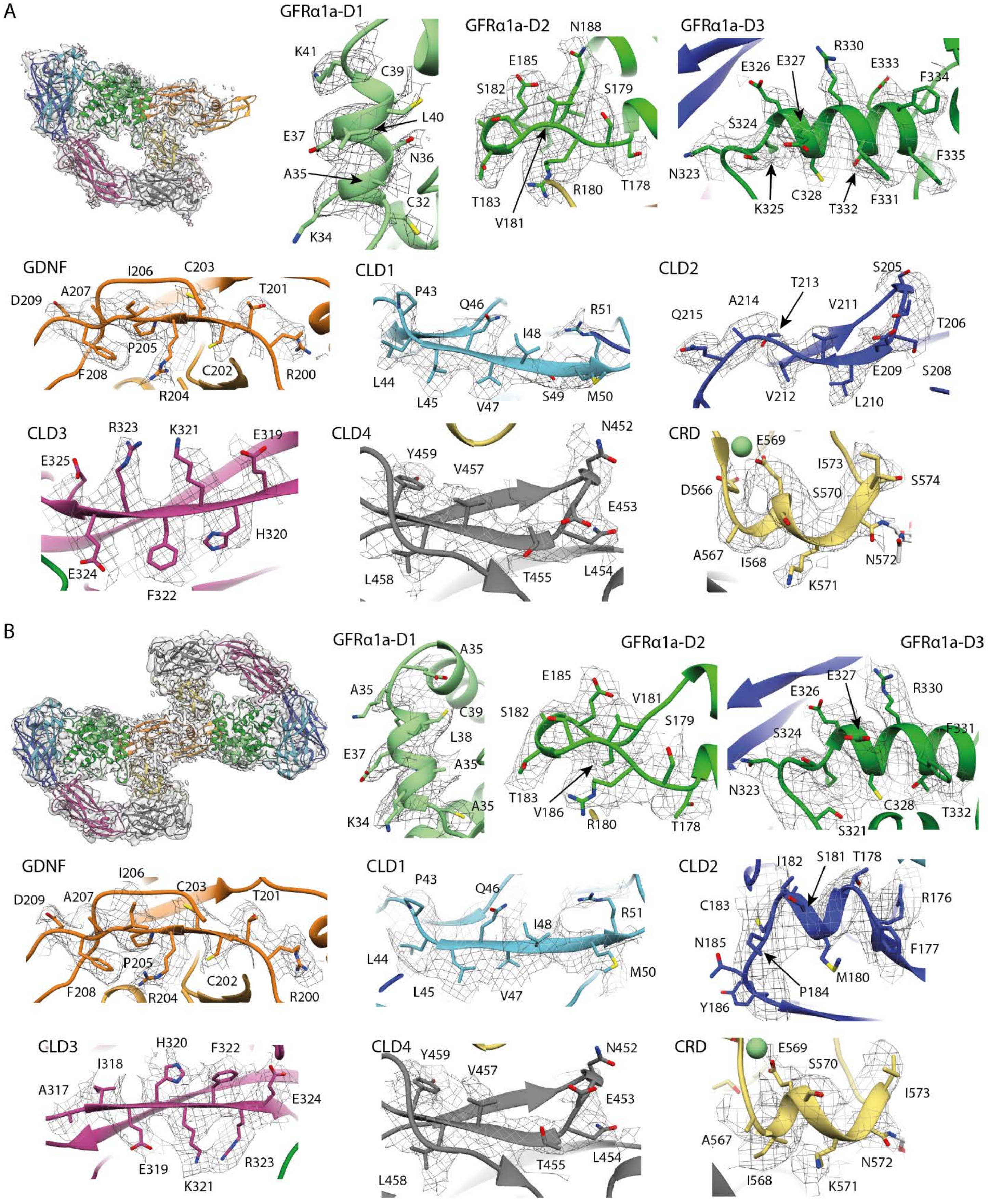

**Supplementary Figure 12: Quality of different regions of the final zRGα1a cryo-EM map superposed with the final model.** A) The symmetry expanded model and map with regions of each domain highlighted. B) The C2 averaged map and model with sections of each domain highlighted. The residues are represented as sticks and the mainchain as a ribbon, the maps are shown as a mesh. All images were rendered in Chimera (Pettersen et al., 2004).

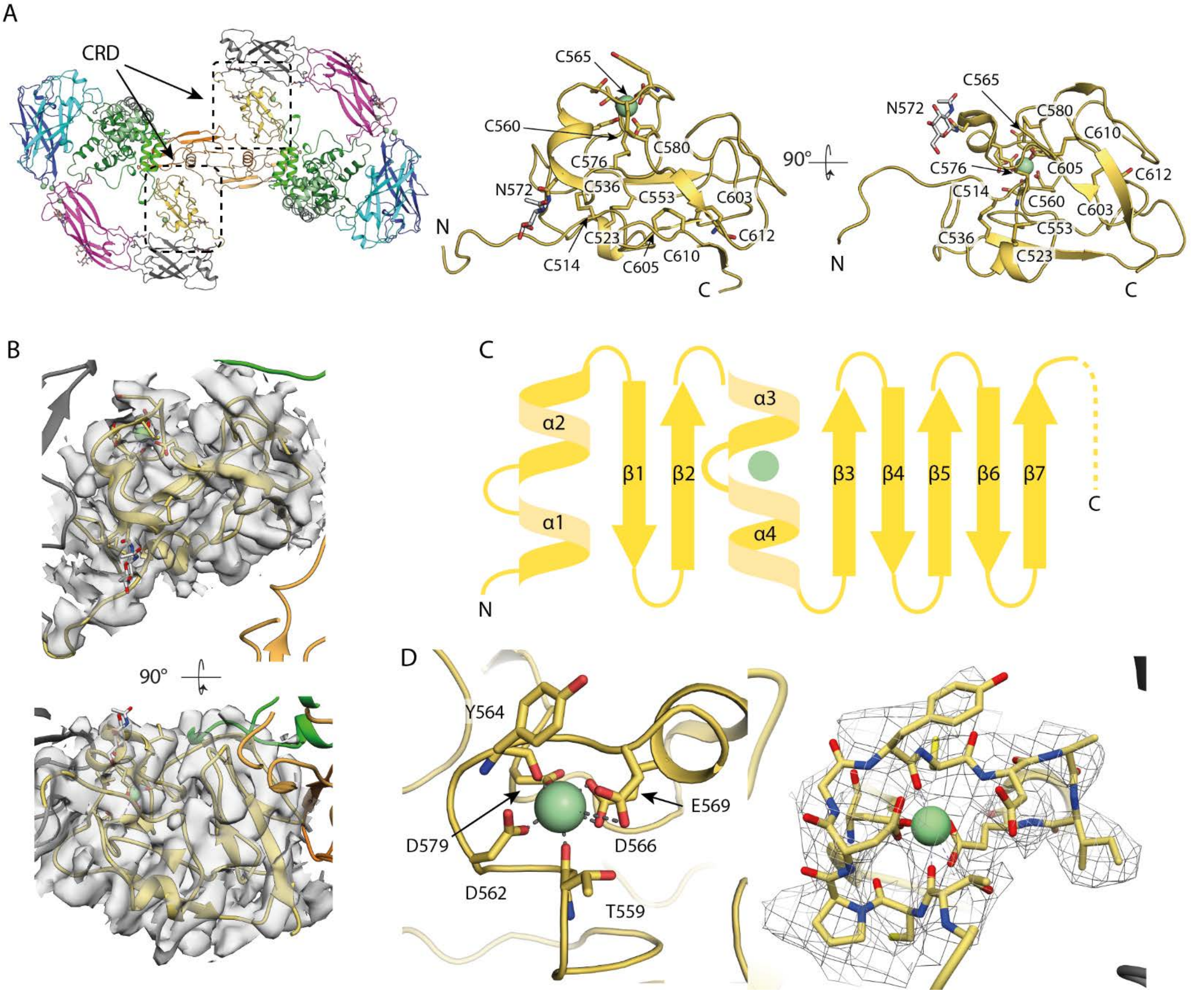

**Supplementary Figure 13: Building the zRET$^{CRD}$ structure.** A) Location of the cysteine rich domain (CRD) in the zRGα1a structure. Two close-up views of the CRD highlighting all cysteines and an N-linked glycosylated asparagine represented as sticks, with the domain shown as a cartoon in yellow and the calcium represented as a green sphere. B) Two close-up views of the model in the symmetry expanded map. C) A topology map of the CRD with the calcium ion represented as a green sphere, the final ten C-terminal residues missing from the structure are represented as a dashed line. D) A close-up of the calcium-binding site superposed with the C2 averaged map, with a calcium represented as a green sphere and the coordinating residues are shown as sticks. Superposition of final zRET$^{CRD}$ structure with hRET$^{CRD}$ gave an RMSD of 2.8Å over 104 C-alpha atoms. The connecting loop between zRET$^{CLD4}$ and zRET$^{CRD}$ (residues 501 to 512) adopts a quite different trajectory from that seen in hRET despite very similar positions of each domain in zRET and hRET. Images rendered in PyMOL (Schrodinger, 2015) and Chimera (Pettersen et al., 2004).

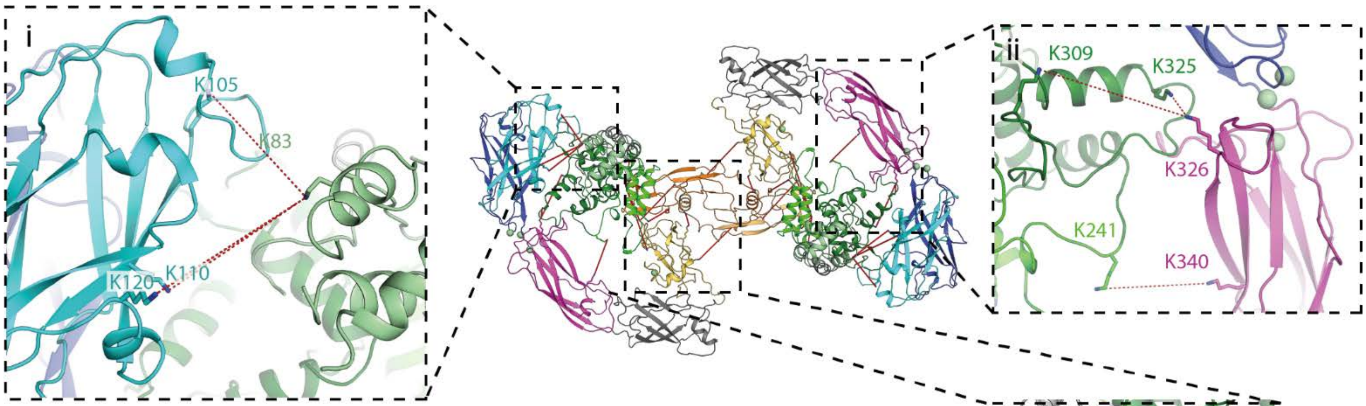

| Score | Protein1 | PepPos1 | PepSeq1 | LinkPos1 | Protein2 | PepPos2 | PepSeq2 | LinkPos2 | |
|---|---|---|---|---|---|---|---|---|---|
| 212.03 | GDNF | 192 | KLDKDTPSR | 1 | RETECD | 544 | DKGISKR | 2 | |
| 182.35 | GFRa1 | 309 | SPKISVSPFCDCSSSGNSKEECDR | 3 | RETECD | 326 | FREEKAIFGNVR | 5 | |
| 169.05 | RETECD | 609 | AGHGTCYCFEGKCFCER | 12 | GDNF | 226 | KHSAK | 1 | |
| 163.62 | GFRa1 | 325 | ISVSPFCDCSSSGNSKEECDRFTEFFTDNACLR | 16 | RETECD | 326 | FREEKAIFGNVR | 5 | |
| 155.38 | GDNF | 161 | TKEELIFR | 2 | RETECD | 544 | DKGISKR | 2 | |
| 147.34 | RETECD | 517 | TDEPRFPACAEKR | 12 | GDNF | 136 | VKGQGR | 2 | * |
| 140.5 | RETECD | 597 | DLYGIKAGHGTCYCFEGK | 6 | GDNF | 161 | TKEELIFR | 2 | |
| 127.49 | GDNF | 195 | KLDKDTPSR | 4 | RETECD | 511 | FPACAEKR | 7 | |
| 121.09 | GFRa1 | 158 | ENNCLNAAKACNLNDTCKK | 9 | GDNF | 161 | TKEELIFR | 2 | |
| 118.13 | GFRa1 | 83 | LVLDALKQSPLYNCR | 7 | RETECD | 105 | SPKDLTLK | 3 | |
| 118.11 | GFRa1 | 149 | LAPIYSGEPALAKENNCLNAAK | 13 | GDNF | 161 | TKEELIFR | 2 | |
| 112.24 | GDNF | 225 | TCCRPIAFDDDISFLDDSLEYHTLKK | 25 | GFRa1 | 191 | RKCHK | 2 | |
| 111.66 | RETECD | 120 | VGVSSTPPMKVMCTILPTVEVK | 10 | GFRa1 | 83 | LVLDALKQSPLYNCR | 7 | |
| 99.02 | GFRa1 | 48 | ANELCLKEPGCSSKYR | 14 | GDNF | 231 | KCACV | 1 | # |
| 92.69 | GFRa1 | 241 | RQTIVPACSYEDKERPNCLTLQVSCK | 13 | RETECD | 340 | GTVHEYKLK | 7 | |
| 82.45 | GFRa1 | 83 | LVLDALKQSPLYNCR | 7 | RETECD | 110 | DLTLKVGVSSTPPMK | 5 | |
| 79.74 | GFRa1 | 34 | LDCVKANELCLK | 5 | GDNF | 136 | VKGQGR | 2 | * |

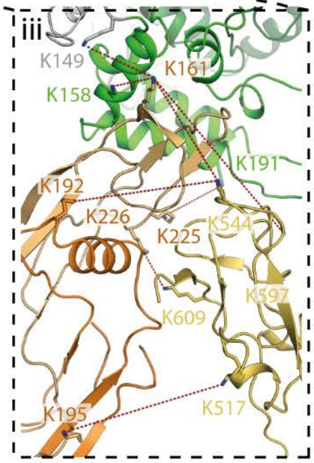

**Supplementary Figure 14: Mapping intermolecular crosslinks in the zRGα1a complex by XL-MS (Cross-Linking Mass Spectrometry).** The overall C2 zRGα1a model represented as a cartoon with the crosslinks formed between lysines using disuccinimidyl sulfoxide (DSSO) represented as dashed-red lines between crosslinked lysines represented as sticks. The model is coloured by domain according to Supplementary Figure 1. i) Close-up of zGFRα1a domain 1 crosslinking to CLD1 of zRET$^{ECD}$. ii) Close-up of lysines from CLD3 on the zRET$^{ECD}$ crosslinked to lysines from zGFRα1a domains 2 and 3, the view is 180° relative to the y-axis. iii) Close up of CRD-zGDNF-zGFRα1a interface, with the intermolecular crosslinks between the lysines of CRD from zRET$^{ECD}$ with zGDNF and zGFRα1a with zGDNF. All images are rendered in PyMOL(Schrodinger, 2015). The table inset shows all the intermolecular crosslinks with each peptide highlighted according to each protein; zRET$^{ECD}$ in cyan, zGDNF in orange and zGFRα1a in green. The crosslinked peptides highlighted with * do not have any structural model therefore are not shown in the model. The crosslinked peptide highlighted with # represents domain 1 of zGFRα1a cross-linked to zGDNF which has a distance of ~59 Å. This may indicate that zGFRα1a$^{D1}$ is quite mobile consistent with the poorer quality of the map for this domain.

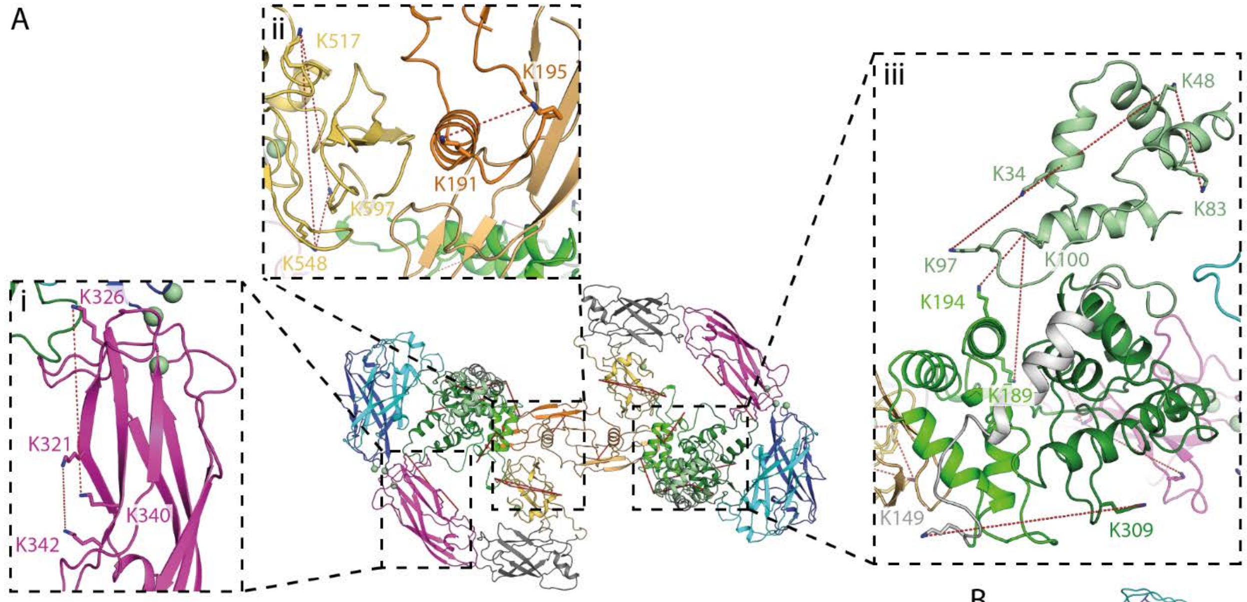

| Score | Protein1 | PepPos1 | PepSeq1 | LinkPos1 | Protein2 | PepPos2 | PepSeq2 | LinkPos2 | |
|---|---|---|---|---|---|---|---|---|---|
| 214.96 | GDNF | 195 | KLDKDTPSR | 4 | GDNF | 136 | VKGQGR | 2 | * |
| 171.33 | GFRa1 | 189 | VSTAEVCNKR | 9 | GFRa1 | 100 | EKNCLR | 2 | |
| 166.9 | GFRa1 | 71 | ESNFSMATGMEAKDECR | 13 | GFRa1 | 48 | EPGCSSKYR | 7 | * |
| 163.04 | GDNF | 161 | TKEELIFR | 2 | GDNF | 136 | VKGQGR | 2 | * |
| 159.98 | GFRa1 | 309 | SPKISVSPFCDCSSSGNSKEECDR | 3 | GFRa1 | 149 | LAPIYSGEPALAKENNCLNAAK | 13 | |
| 147.39 | RETECD | 326 | FREEKAIFGNVR | 5 | RETECD | 340 | GTVHEYKLK | 7 | |
| 144.03 | GDNF | 191 | ILNNLTHNKK | 9 | GDNF | 136 | VKGQGR | 2 | * |
| 133.6 | RETECD | 517 | FPACAEKR | 7 | RETECD | 548 | DKGISKR | 6 | |
| 119.14 | GDNF | 191 | YCSGPCHDAETNYDKILNNLTHNKK | 24 | GDNF | 195 | LDKDTPSR | 3 | |
| 118.75 | RETECD | 597 | DLYGIKAGHGTCYCFEGK | 6 | RETECD | 548 | DKGISKR | 6 | |
| 117.99 | RETECD | 342 | LKLSQNLSVTEQR | 2 | RETECD | 320 | NNFAIEHKFR | 8 | |
| 104 | GFRa1 | 34 | LDCVKANELCLK | 5 | GFRa1 | 97 | RGMKK | 4 | |
| 100.67 | GFRa1 | 194 | KCHKALR | 4 | GFRa1 | 100 | EKNCLR | 2 | |
| 99.02 | GFRa1 | 47 | ANELCLKEPGCSSKYR | 14 | GFRa1 | 83 | LVLDALKQSPLYNCR | 7 | |
| 98.46 | RETECD | 597 | DLYGIKAGHGTCYCFEGK | 6 | RETECD | 517 | FPACAEKR | 7 | |
| 88.56 | GFRa1 | 71 | ESNFSMATGMEAKDECR | 13 | GFRa1 | 83 | LVLDALKQSPLYNCR | 7 | * |
| 81.28 | GFRa1 | 34 | LDCVKANELCLK | 5 | GFRa1 | 47 | EPGCSSKYR | 7 | |
| 70.4 | RETECD | 340 | GTVHEYKLK | 7 | RETECD | 105 | SPKDLTLK | 3 | |

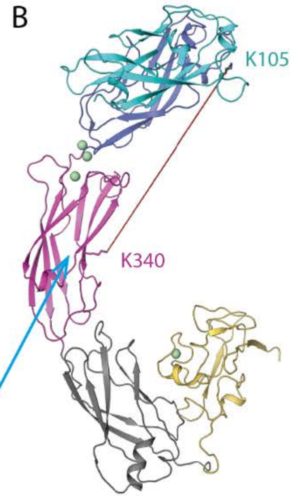

**Supplementary Figure 15: Intramolecular crosslinks within the zRGα1a complex.** A) The overall C2 zRGα1a model represented as a cartoon with the crosslinks formed between lysines using disyccinimidyl sulfoxide (DSSO) represented as red lines between crosslinked lysines represented as sticks. The model is coloured according to its domains; CLD1 in cyan, CLD2 in blue, CLD3 in magenta, CLD4 in grey, CRD in yellow, zGFRα1a$^{D1}$ in pale green, zGFRα1a$^{D2}$ in green, zGFRα1a$^{D3}$ in dark green and zGDNF in orange. i) Close-up of intramolecular crosslinks located in CLD3 from zRET$^{ECD}$. ii) Close-up of the intramolecular crosslinked lysines from CRD from zRET$^{ECD}$ and zGDNF. iii) The intramolecular crosslinks between the lysines located in zGFRα1a confirm the location of zGFRα1a$^{D1}$ adjacent to zGFRα1a$^{D3}$. B) The structure of zRET$^{ECD}$ alone with a lysine located in CLD1 crosslinked to a lysine located in CLD3 spanning a distance of 60 Å, this may indicate flexibility at the calcium-binding region may be even greater in solution than observed in the crysals. All images are rendered in PyMOL(Schrodinger, 2015). The table inset shows all the intramolecular crosslinks with each peptide highlighted according to each protein; zRET$^{ECD}$ in cyan, zGDNF in orange and zGFRα1a in green. The crosslinked peptides highlighted with *are not present in the model and are therefore not shown.

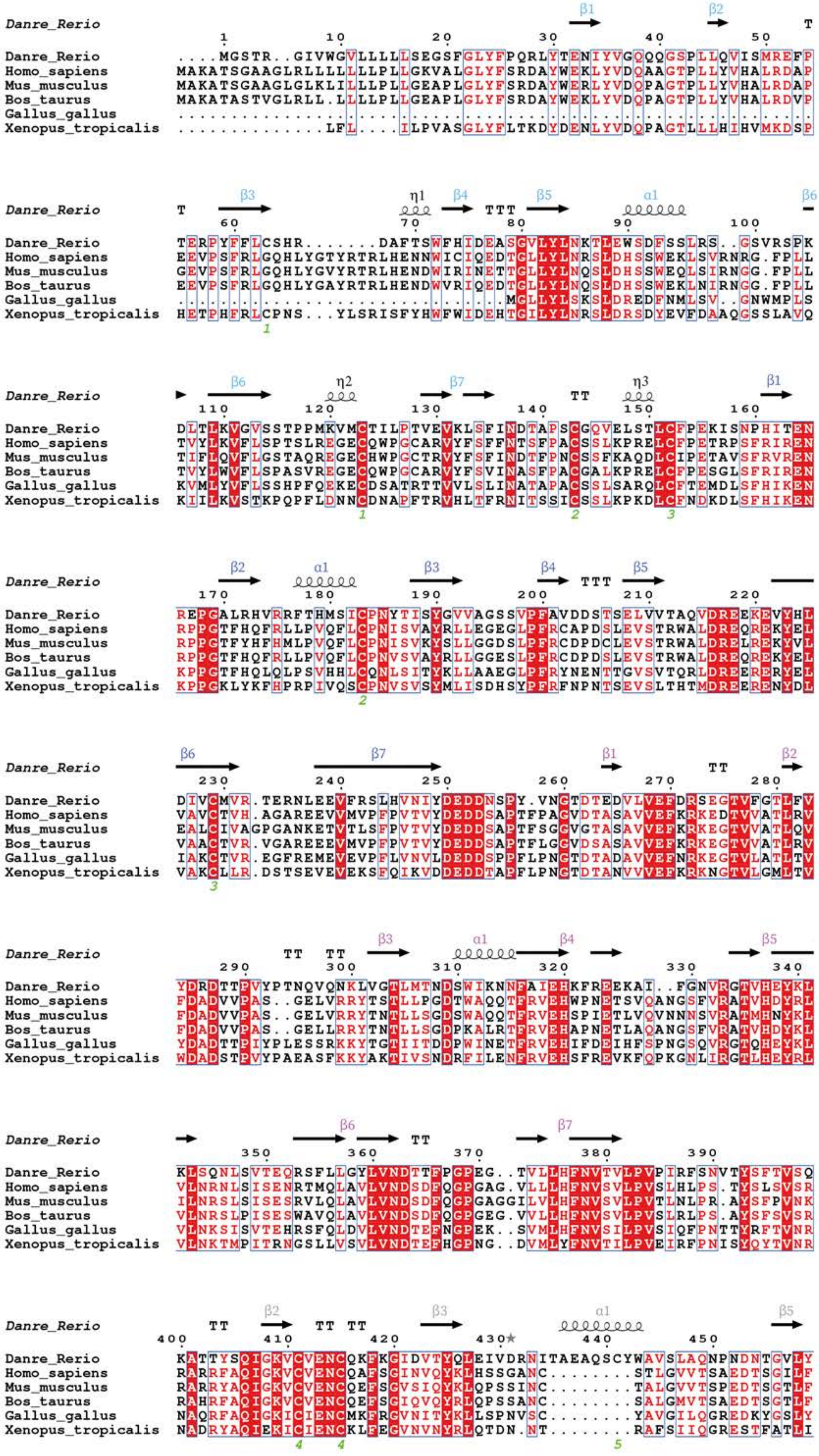
Danre_Rerio
Homo_sapiens
Mus_musculus
Bos_taurus
Gallus_gallus
Xenopus_tropicalis

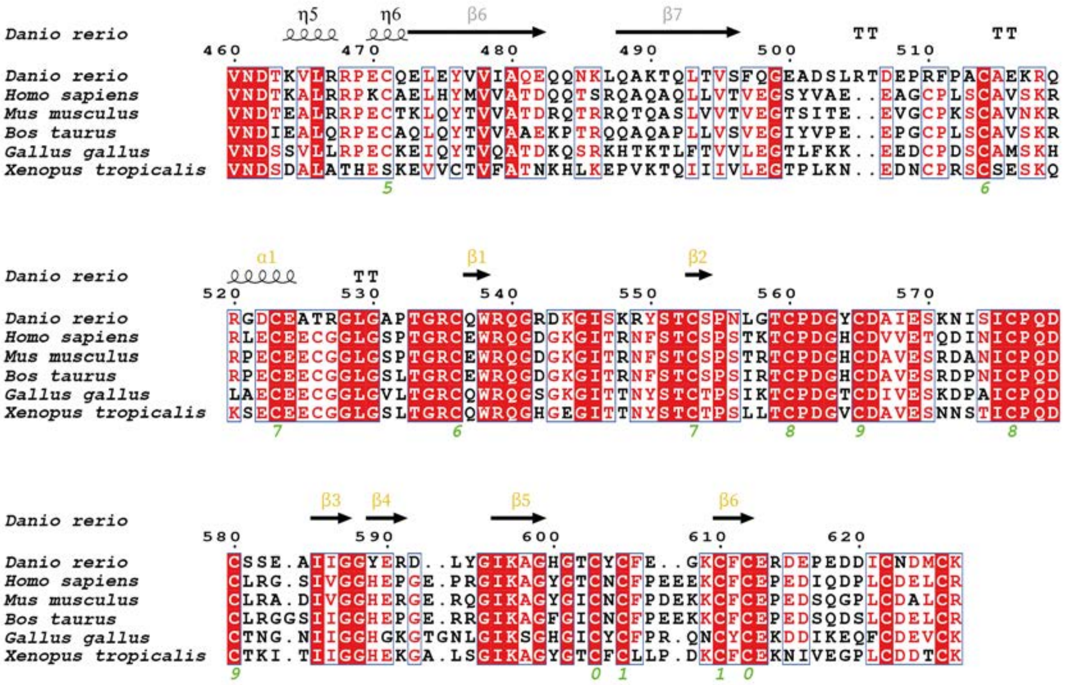

**Supplementary Figure 16: Sequence alignment of $zRET^{ECD}$ and selected RET orthologues.** A Sequence alignment of three higher vertebrate and three lower vertebrate species of RET. *Danio rerio* (Uniprot P04749), *Homo sapiens* (Uniprot P04749), *Mus Musculus* (Uniprot P04749), *Bos Taurus* (Uniprot P04749), *Gallus gallus* (Uniprot P04749), and *Xenopus tropicalis* (Uniprot P04749). The secondary structural elements are highlighted according to the domains; cyan for CLD1, blue for CLD2, magenta for CLD3, grey for CLD4 and yellow for the CRD. $zRET^{ECD}$ from the zRGα1a structure was used to identify the secondary structural elements. The structural alignments were performed using Clustal Omega(Sievers et al., 2011) and the representation was done using ESPript (http://espript.ibcp.fr)(Robert and Gouet, 2014).

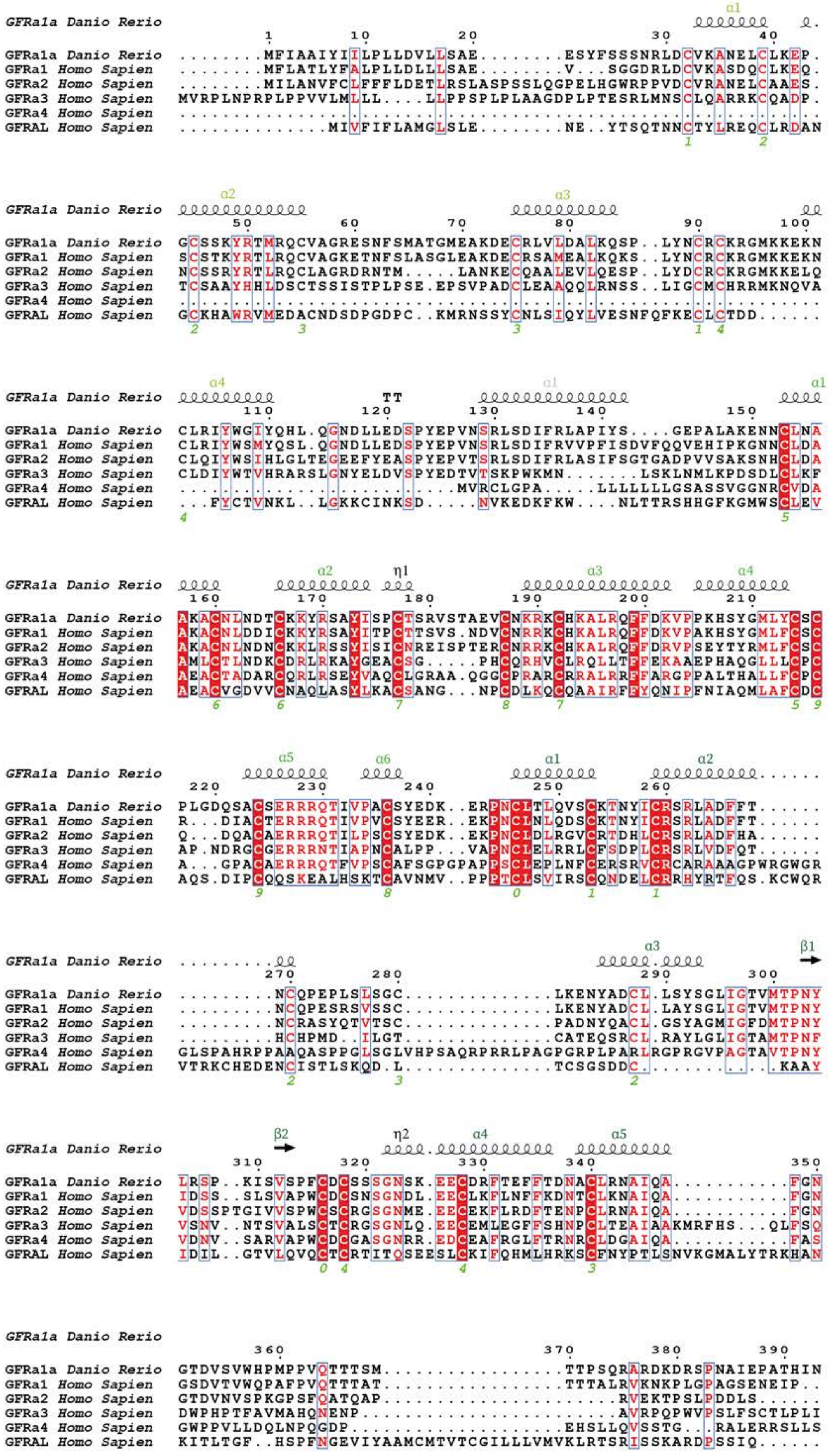

GFRa1a Danio Rerio ........MFIAAIYIILPLLDVLLSAE........ESYFSSSNRLDCVKANELCLKEP.
GFRa1 Homo Sapien ........MFLATLYFALPLLDLLLSAE........V...SGGDRLDCVKASDQCLKEQ.
GFRa2 Homo Sapien ........MILANVFCLFFFLDETLRSLASPSSLQGPELHGWRPPVDCVRANELCAAES.
GFRa3 Homo Sapien MVRPLNPRPLPPVVLMLLL....LLPPSPLPLAAGDPLPTESRLMNSCLQARRKCQADP.
GFRAL Homo Sapien ..............MIVFIFLAMGLSLE........NE..YTSQTNNCTYLREQCLRDAN
GFRa1a Danio Rerio GCSSKYRTMRQCVAGRESNFSMATGMEAKDECRLVLDALKQSP..LYNCRCKRGMKKEKN
GFRa1 Homo Sapien SCSTKYRTLRQCVAGKETNFSLASGLEAKDECRSAMEALKQKS..LYNCRCKRGMKKEKN
GFRa2 Homo Sapien NCSSRYRTLRQCLAGRDRNTM.....LANKECQAALEVLQESP..LYDCRCKRGMKKELQ
GFRa3 Homo Sapien TCSAAYHHLDSCTSSISTPLPSE.EPSVPADCLEAAQQLRNSS..LIGCMCHRRMKNQVA
GFRAL Homo Sapien GCKHAWRVMEDACNDSDPGDPC..KMRNSSYCNLSIQYLVESNFQFKECLCTDD......
GFRa1a Danio Rerio CLRIYWGIYQHL.QGNDLLEDSPYEPVNSRLSDIFRLAPIYS....GEPALAKENNCLNA
GFRa1 Homo Sapien CLRIYWSMYQSL.QGNDLLEDSPYEPVNSRLSDIFRVVPFISDVFQQVEHIPKGNNCLDA
GFRa2 Homo Sapien CLQIYWSIHLGLTEGEEFYEASPYEPVTSRLSDIFRLASIFSGTGADPVVSAKSNHCLDA
GFRa3 Homo Sapien CLDIYWTVHRARSLGNYELDVSPYEDTVTSKPWKMN.......LSKLNMLKPDSDLCLKF
GFRa4 Homo Sapien ...........................MVRCLGPA.....LLLLLLGSASSVGGNRCVDA
GFRAL Homo Sapien ...FYCTVNKL..LGKKCINKSD.....NVKEDKFKW....NLTTRSHHGFKGMWSCLEV
GFRa1a Danio Rerio AKACNLNDTCKKYRSAYISPCTSRVSTAEVCNKRKCHKALRQFFDKVPPKHSYGMLYCSC
GFRa1 Homo Sapien AKACNLDDICKKYRSAYITPCTTSVS.NDVCNRRKCHKALRQFFDKVPAKHSYGMLFCSC
GFRa2 Homo Sapien AKACNLNDNCKKLRSSYISICNREISPTERCNRRKCHKALRQFFDRVPSEYTYRMLFCSC
GFRa3 Homo Sapien AMLCTLNDKCDRLRKAYGEACSG.....PHCQRHVCLRQLLTFFEKAAEPHAQGLLLCPC
GFRa4 Homo Sapien AEACTADARCQRLRSEYVAQCLGRAA.QGGCPRARCRRALRRFFARGPPALTHALLFCPC
GFRAL Homo Sapien AEACVGDVVCNAQLASYLKACSANG...NPCDLKQCQAAIRFFYQNIPFNIAQMLAFCDC
GFRa1a Danio Rerio PLGDQSACSERRRQTIVPACSYEDK..ERPNCLTLQVSCKTNYICRSRLADFFT......
GFRa1 Homo Sapien R...DIACTERRRQTIVPVCSYEER..EKPNCLNLQDSCKTNYICRSRLADFFT......
GFRa2 Homo Sapien Q...DQACAERRRQTILPSCSYEDK..EKPNCLDLRGVCRTDHLCRSRLADFHA......
GFRa3 Homo Sapien AP.NDRGCGERRRNTIAPNCALPP...VAPNCLELRRLCFSDPLCRSRLVDFQT......
GFRa4 Homo Sapien A...GPACAERRRQTFVPSCAFSGPGPAPPSCLEPLNFCERSRVCRCARAAAGPWRGWGR
GFRAL Homo Sapien AQS.DIPCQQSKEALHSKTCAVNMV..PPPTCLSVIRSCQNDELCRRHYRTFQS.KCWQR
GFRa1a Danio Rerio .........NCQPEPLSLSGC.............LKENYADCL.LSYSGLIGTVMTPNY
GFRa1 Homo Sapien .........NCQPESRSVSSC.............LKENYADCL.LAYSGLIGTVMTPNY
GFRa2 Homo Sapien .........NCRASYQTVTSC.............PADNYQACL.GSYAGMIGFDMTPNY
GFRa3 Homo Sapien .........HCHPMD..ILGT.............CATEQSRCL.RAYLGLIGTAMTPNF
GFRa4 Homo Sapien GLSPAHRPPAAQASPPGLSGLVHPSAQRPRRLPAGPGRPLPARLRGPRGVPAGTAVTPNY
GFRAL Homo Sapien VTRKCHEDENCISTLSKQD.L.............TCSGSDDC.............KAAY
GFRa1a Danio Rerio LRSP..KISVSPFCDCSSSGNSK.EECDRFTEFFTDNACLRNAIQA...........FGN
GFRa1 Homo Sapien IDSS..SLSVAPWCDCSNSGNDL.EECLKFLNFFKDNTCLKNAIQA...........FGN
GFRa2 Homo Sapien VDSSPTGIVVSPWCSCRGSGNME.EECEKFLRDFTENPCLRNAIQA...........FGN
GFRa3 Homo Sapien VSNV..NTSVALSCTCRGSGNLQ.EECEMLEGFFSHNPCLTEAIAAKMRFHS...QLFSQ
GFRa4 Homo Sapien VDNV..SARVAPWCDCGASGNRR.EDCEAFRGLFTRNRCLDGAIQA...........FAS
GFRAL Homo Sapien IDIL..GTVLQVQCTCRTITQSEESLCKIFQHMLHRKSCFNYPTLSNVKGMALYTRKHAN
GFRa1a Danio Rerio GTDVSVWHPMPPVQTTTSM.................TTPSQRARDKDRSPNAIEPATHIN
GFRa1 Homo Sapien GSDVTVWQPAFPVQTTTAT.................TTTALRVKNKPLGPAGSENEIP..
GFRa2 Homo Sapien GTDVNVSPKGPSFQATQAP.......................RVEKTPSLPDDLS......
GFRa3 Homo Sapien DWPHPTFAVMAHQNENP.......................AVRPQPWVPSLFSCTLPLI
GFRa4 Homo Sapien GWPPVLLDQLNPQGDP...................EHSLLQVSSTG...RALERRSLLS
GFRAL Homo Sapien KITLTGF..HSPFNGEVIYAAMCMTVTCGILLLVMVKLRTSRISSKARDPSSIQ......

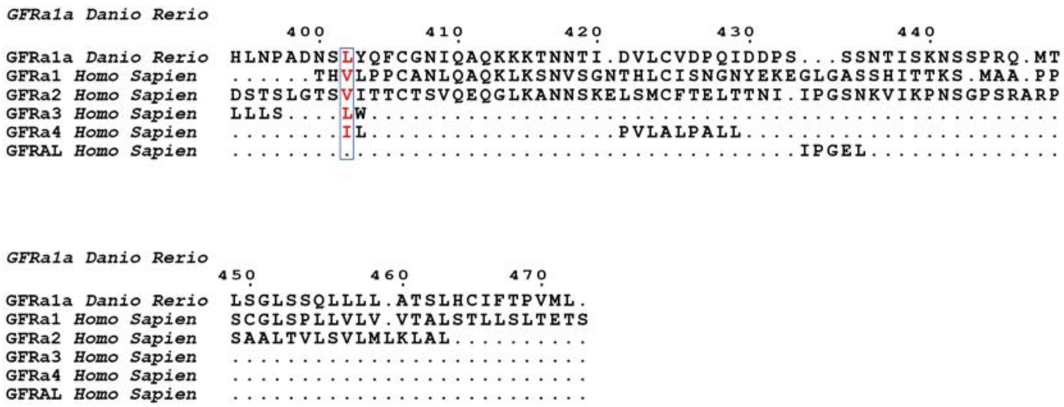

**Supplementary Figure 17: Sequence alignments for zGFRα1a and selected hGFR co-receptors.** A sequence alignment of zGFRα1a (Uniprot Q98TT9), hGFRα1 (Uniprot P56159), hGFRα2 (Uniprot O00451), hGFRα3 (Uniprot O60609), hGFRα4 (Uniprot Q9GZZ7), and hGFRAL (Uniprot Q6UXV0). The secondary structural elements are highlighted according to the domains; pale green for D1, green for D2 and dark green for D3. A linker α1 helix is shown between $zGFR\alpha1a^{D1}$ and $zGFR\alpha1a^{D2}$ in black. Secondary structural elements were defined from the zRGα1a structure. Structural alignments were performed using Clustal Omega(Sievers et al., 2011) and the representation was done using ESPript (http://espript.ibcp.fr)(Robert and Gouet, 2014).

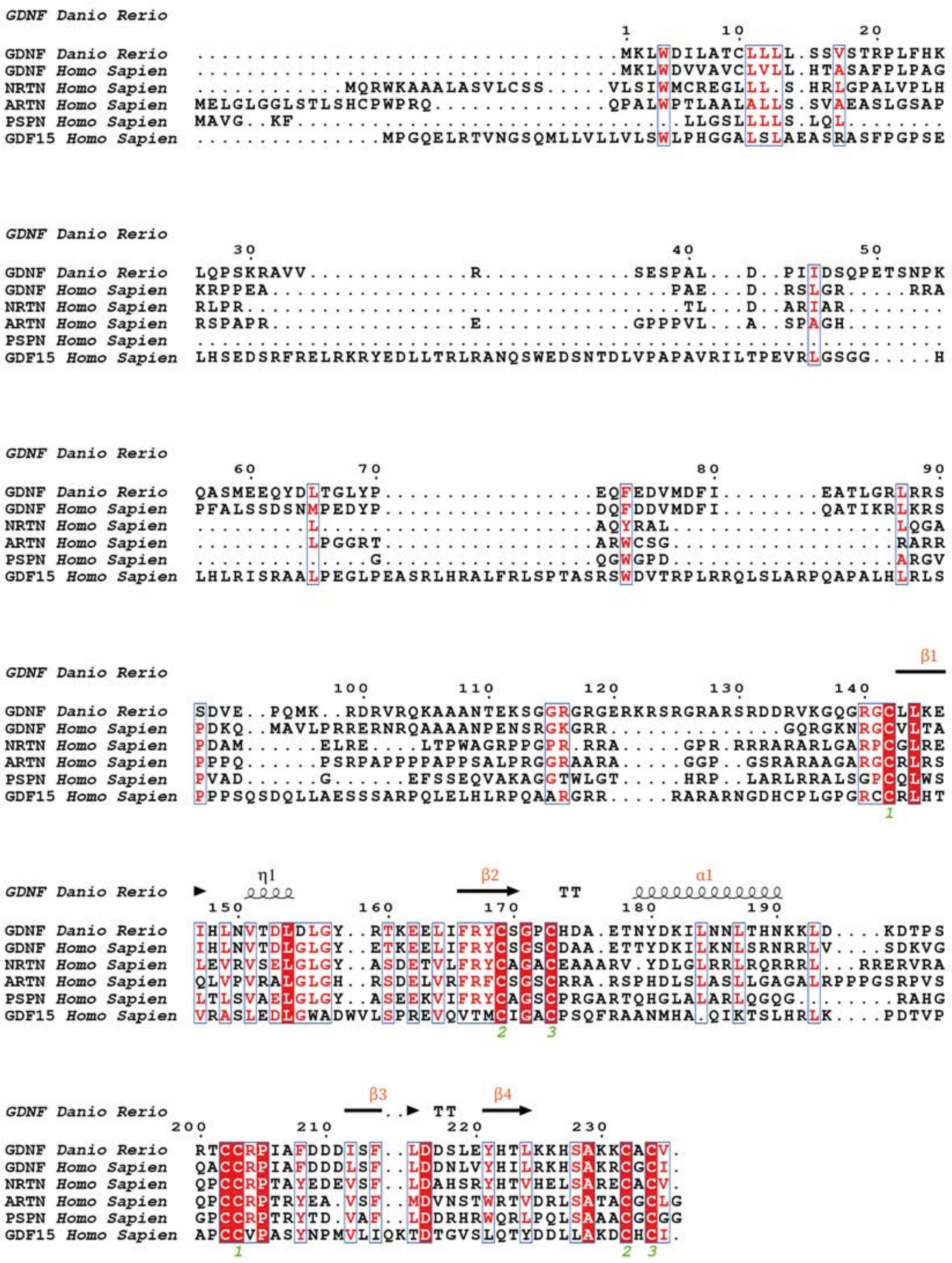

**Supplementary Figure 18: Sequence alignments for zGDNF and selected hGFLs.** A sequence alignment of zGDNF (Uniprot Q98TU0), hGDNF (Uniprot P39905), hNRTN (Uniprot Q99748), hARTN (Uniprot Q5T4W7), hPSPN (Uniprot O60542), and hGDF15 (Uniprot Q99988). The secondary structural details are highlighted in orange as per the colour used in the renderings of the structure. zGDNF from the zRGα1a structure was used to identify the secondary structural elements. The structural alignments were performed using Clustal Omega(Sievers et al., 2011) and the representation was done using ESPript (http://espript.ibcp.fr)(Robert and Gouet, 2014).

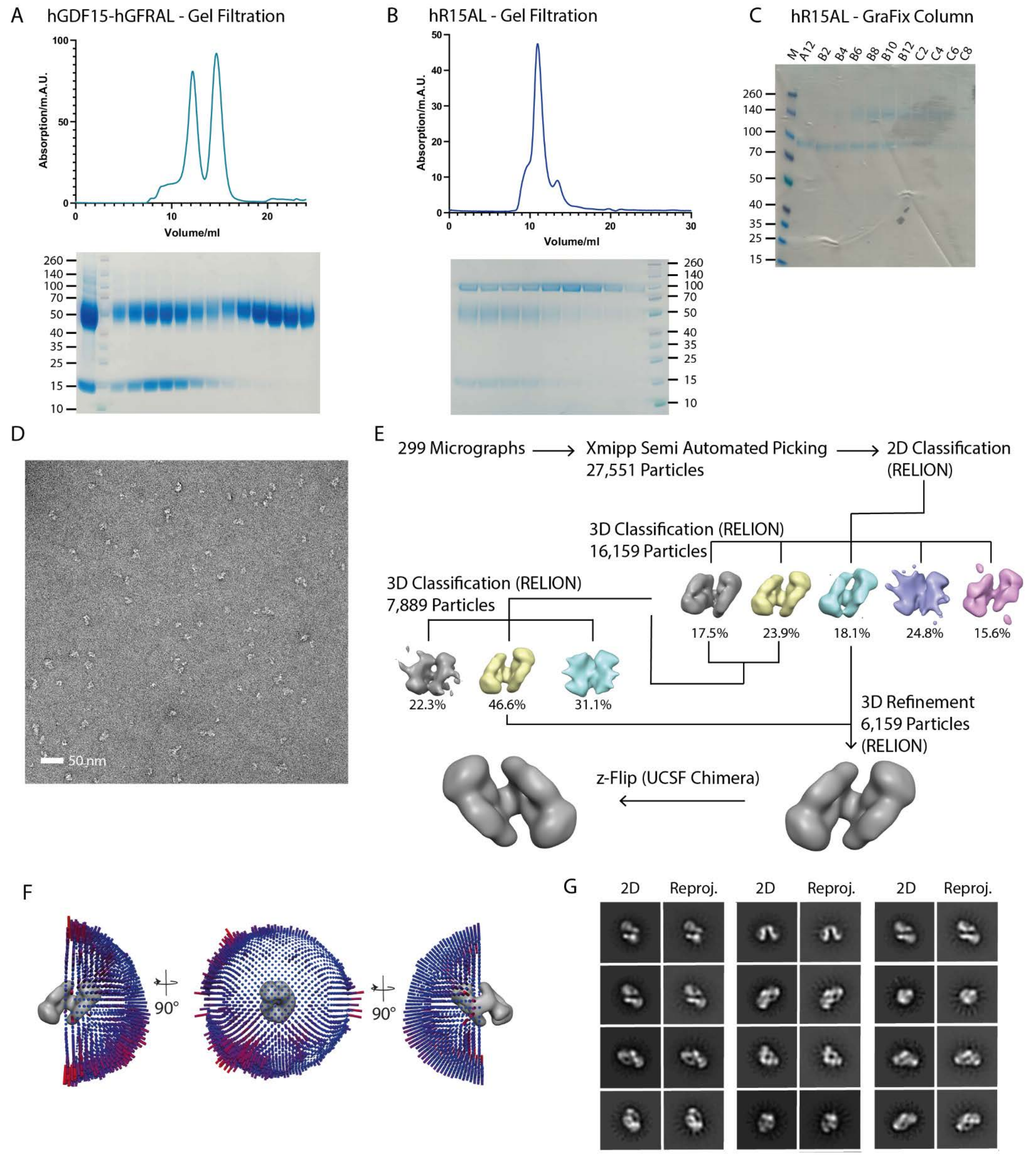

**Supplementary Figure 19: hR15AL sample preparation and EM data processing.** Size exclusion profiles and SDS-PAGE's of A) hGDF15-hGFRAL and B) hRET-hGDF15-hGFRAL (hR15AL). C) An SDS-PAGE of the fractions from the GraFix (Gradient Fixation) of hR15AL. D) A representative negative stain micrograph of the hR15AL-XL complex. E) The data processing route to achieve the final negative stain envelope of hR15AL. F) The particle distribution in the negative stain envelope with C2 symmetry applied. G) Projection matching, performed using Xmipp projection match(De la Rosa-Trevín et al., 2013), between the RELION (Kimanius et al., 2016; Scheres, 2012; Zivanov et al., 2018) 2D class averages from the particles that comprise the final reconstruction and different view of the 3D envelope.

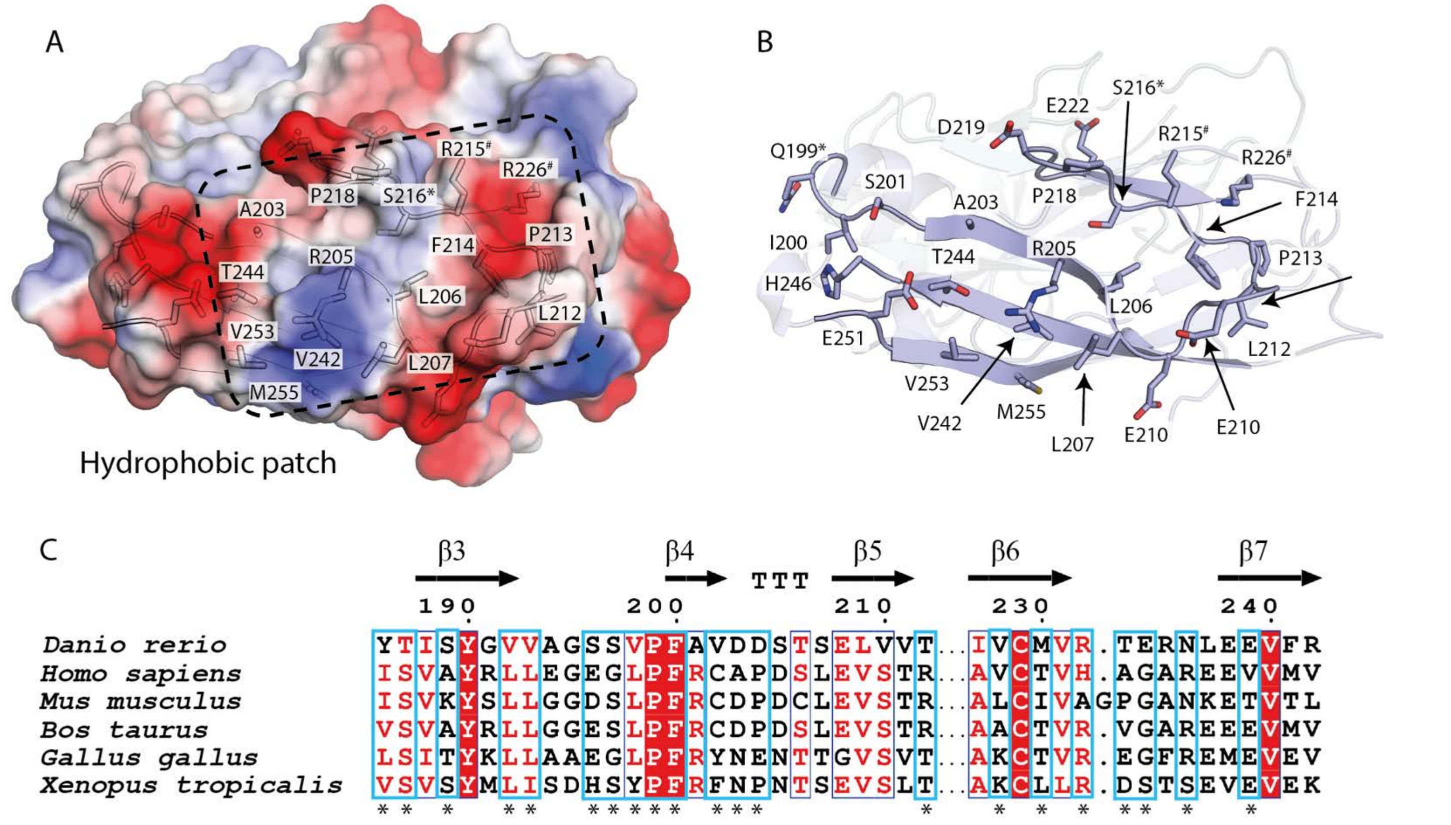

**Supplementary Figure 20: Equivalent multimer contact surface in hRET$^{CLD(1-2)}$ to zRET.** A) Electrostatic potential map of hCLD1-2 (PDB 2X2U)(Kjær et al., 2010) generated in PyMOL, with an overlay of the cartoon trace and key residues highlighted as sticks (Schrodinger, 2015). A hydrophobic area is highlighted, the majority of residues are hydrophobic, with the exception of R205, R215 and R226. # Residues truncated due to lack of electron density, * residue mutated C216S. B) Cartoon representation of the hCLD1-2 structure with key residues highlighted as sticks, CLD1 in pale cyan and CLD2 in pale blue. C) A portion of the Sequence alignment of three higher vertebrate and three lower vertebrate RET sequences that corresponds to the putative interaction site. The sequences are *Danio rerio* (Uniprot P04749)*, Homo sapiens* (Uniprot P04749)*, Mus musculus* (Uniprot P04749), *Bos taurus* (Uniprot P04749), *Gallus gallus* (Uniprot P04749), and *Xenopus tropicalus* (Uniprot P04749). The key residues highlighted in cyan boxes and an asterix. The sequence alignment was peformed using Clustal Omega (Sievers et al., 2011) and the representation was produced using ESPript (http://espript.ibcp.fr)(Robert and Gouet, 2014).

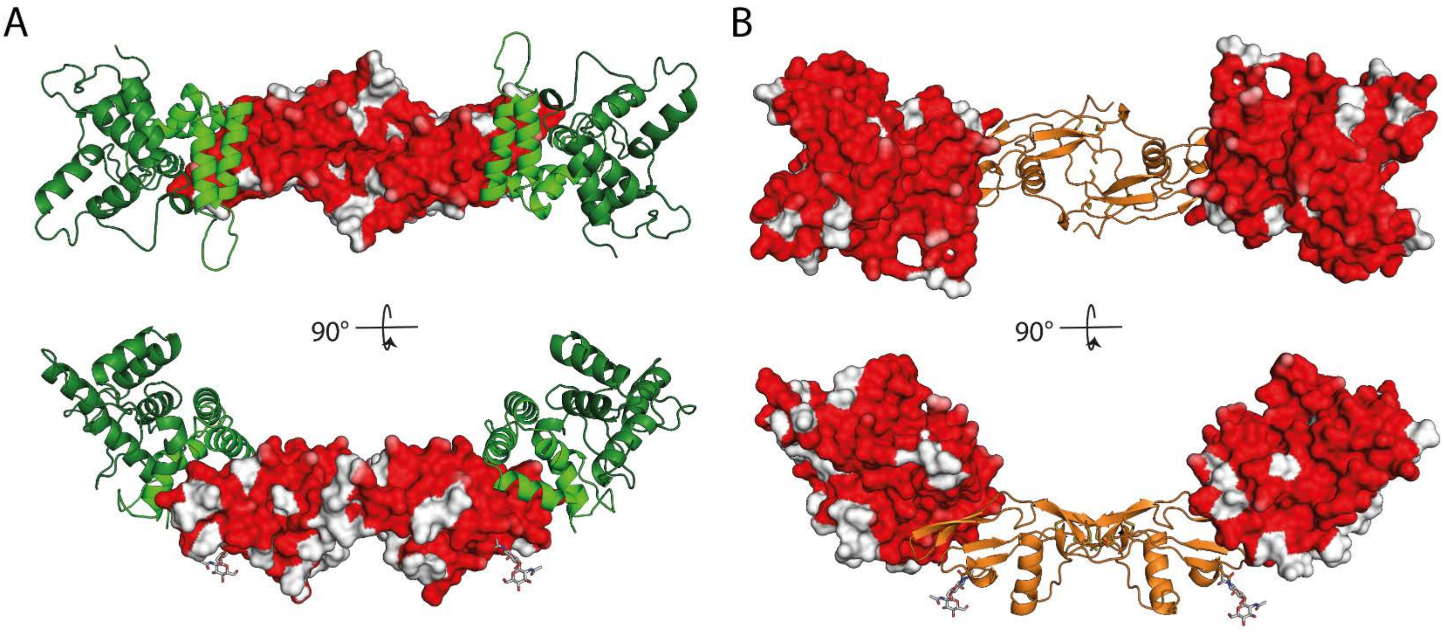

**Supplementary Figure 21: Heat-map of sequence variation between zebrafish and human GFRα1$_2$-GDNF$_2$.** A) Two orthogonal views, of a surface heatmap representation of GDNF coloured by sequence similarity (red highly similar to white, least similar) based on the alignment of zGDNF (Uniprot Q98TU0) and hGDNF (Uniprot P39905). B) Two orthogonal views of a surface heatmap representation of GDNF coloured by sequence similarity (red highly similar to white, least similar) based on the alignment between zGFRα1a (Uniprot Q98TT9) and hGFRα1 (Uniprot P56159). The surface is coloured according to similarity of residue type; aromatic residues (F, W, and Y), aliphatic residues (A, I, L, and V), residues containing an alcohol functional group (S and T), positively charged residues (R and K), negatively charged residues (D and E), and residues with polar sidechains N and Q, and C, G, H and M are counted individually. There are two views, with a 90° rotation Images were rendered in PyMOL (Schrodinger, 2015).

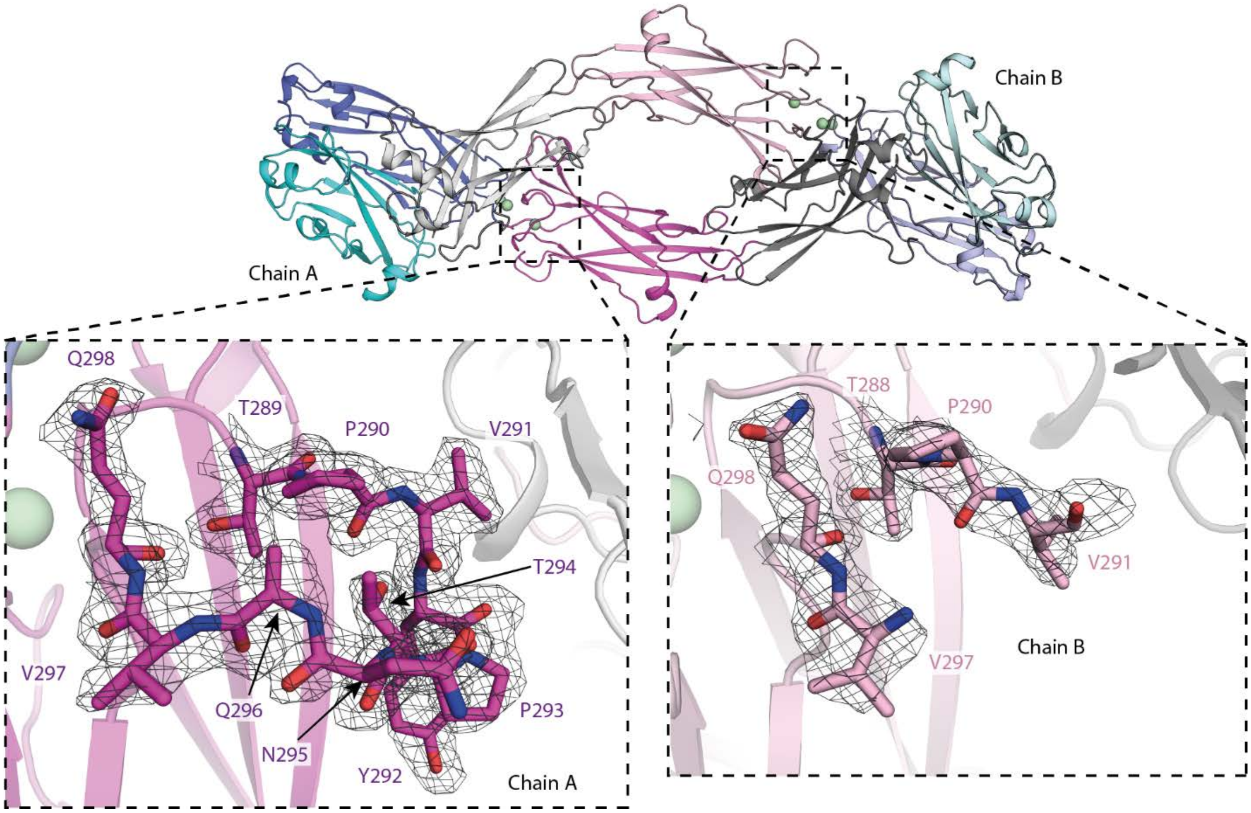

**Supplementary Figure 22: Comparison of CLD3-β2-β3-loop conformations in zRET$^{CLD1\text{-}4}$ chain A and chain B.** A cartoon representation of the intertwined dimer within the crystallographic asymmetric unit, with the two CLD3-β2-β3-loops highlighted in the overall structure. The insets reveal the final electron density calculated using m2Fo-DFc coefficients and contoured at σ=1.0 over the loop shown as sticks. Images rendered in PyMOL (Schrodinger, 2015).

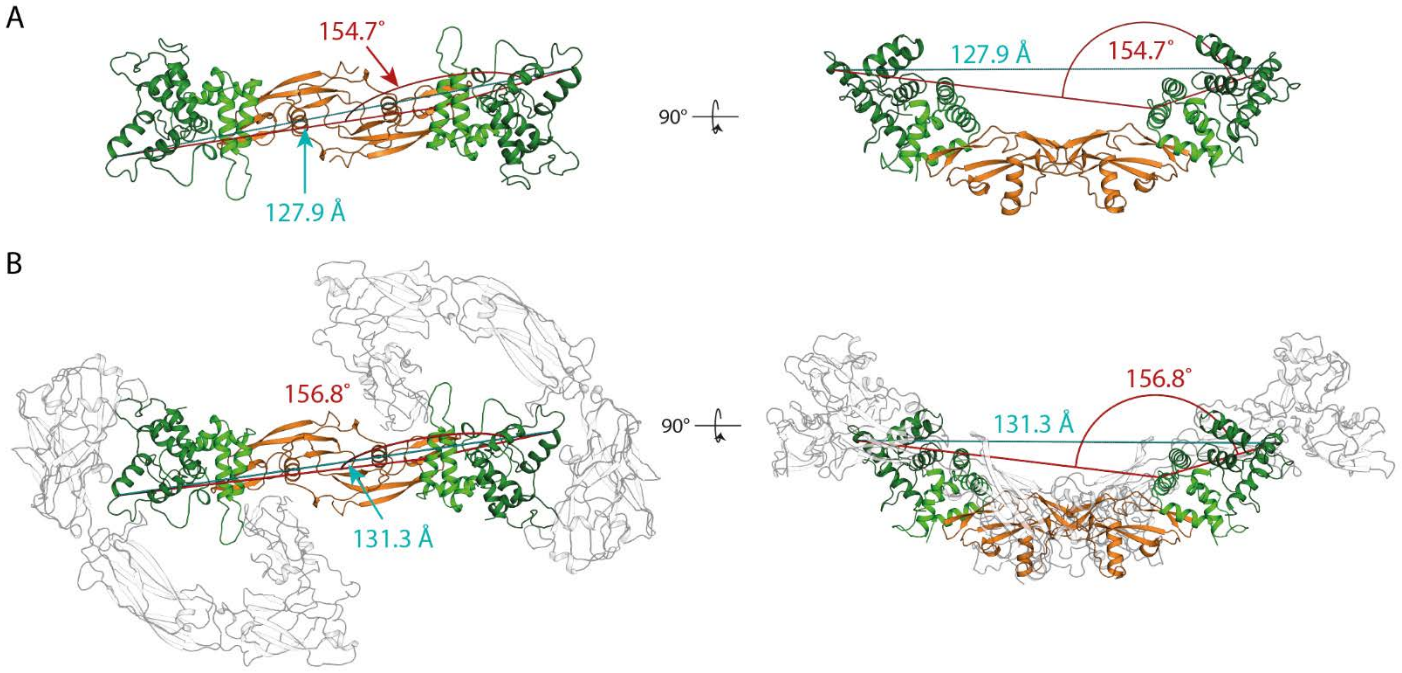

**Supplementary Figure 23: Evidence for limited conformational flexing of zGDNF-zGFRα1a in the presence and absence of zRET$^{ECD}$.** A) The crystal structure of a 2:2 zGDNF-zGFRα1a is shown as a cartoon, with GDNF in light orange, zGFRα1a$^{D2}$ in light green and zGFRα1a$^{D3}$ in green. The distance between K325 from symmetry-related molecules of zGFRα1a is highlighted in teal and the angle between the K325-A172 from one molecule of zGFRα1 and K325 in the second molecule of zGFRα1a in red. B) The 2:2 ligand:co-receptor zGDNF-zGFRα1a$^{D2-D3}$ built into the cryo-EM zRGα1a structure, represented as a cartoon, with zGFRα1a$^{D2}$, zGFRα1a$^{D3}$ and zGDNF in green, forest green and orange, respectively, and zRET$^{ECD}$ and zGFRα1a$^{D1}$ in grey. The distance between K325 from symmetry-related molecules of zGFRα1a highlighted in teal and the angle between the K325-A172 from one molecule of zGFRα1 and K325 in the second molecule of zGFRα1a in purple.
